# Molecular basis behind the isoprene emission diversity in Fagaceae

**DOI:** 10.1101/2025.05.17.654689

**Authors:** Sora Koita, Ryosuke Munakata, Yoko Kamata, Nodoka Shinya, Kenji Fukushima, Atsushi J. Nagano, Yuka Ikezaki, Akiko Satake, Takuya Saito, Kenji Miura, Akifumi Sugiyama, Kazufumi Yazaki

**Affiliations:** Research Institute for Sustainable Humanosphere, Kyoto University, Gokasho, Uji, 611-0011 Japan; Center for Frontier Research, National Institute of Genetics, Yata, Mishima, 451-8540, Shizuoka, Japan; Faculty of Agriculture, Ryukoku University, Seta Oe-cho Yokotani 1-5, Otsu, 520-2194 Japan; Institute for Advanced Biosciences, Keio University, Nipponkoku 403-1, Daihouji, Tsuruoka, 997-0017 Japan; Department of Biology, Kyushu University, Nishi-ku Motooka 744, Fukuoka, 819-0395 Japan; National Institute for Environmental Studies, Onogawa 16-2, Tsukuba, 305-8506 Japan; Graduate School of Life and Environmental Sciences, University of Tsukuba, Tsukuba, 305-8572 Japan; Japan Science and Technology Agency (JST), PRESTO, 4-1-8, Honcho, Kawaguchi-shi, Saitama, 332-0012 Japan

**Keywords:** VOC emission, isoprene synthase, Fagaceae, *Quercus*, terpene synthase

## Abstract

Plants emit a large amount of volatile organic compounds (VOCs) into the atmosphere, reaching approximately 10^9^ tons of carbon per year. These biogenic VOCs exhibit significant chemical diversity, with terpenoids being the dominant group, and isoprene accounting for nearly half of the total biogenic VOCs. Due to its high chemical reactivity, isoprene has a strong impact on atmospheric quality and climate. *Quercus* species (Fagaceae) are known to be the main isoprene emitters in the Northern Hemisphere. However, isoprene synthase is unknown in the entire Fagaceae family. Notably, even within a single genus such as *Quercus*, both isoprene-emitting and non-emitting species are present, yet the molecular basis of this dichotomy remains unclear. Here, we report the identification of the *IspS* gene from the isoprene-emitting species *Quercus serrata* (*QsIspS1*) through seasonal transcriptome analysis and its detailed biochemical characterization. We also identified two genes with high sequence similarity to *QsIspS1* in the genomes of non-emitting species: *Q. glauca* (*QgIspS1-like*) and *Lithocarpus edulis* (*LeIspS1-like*). We discovered mutations in these sequences that likely impair their function. Biochemical analysis revealed that *QgIspS1-like* is a monoterpene synthase, whereas *LeIspS1-like* is a pseudogene incapable of isoprene synthesis, explaining these plants’ inability to emit isoprene. Furthermore, site-directed mutagenesis revealed an amino acid that plays a pivotal role in the substrate and product specificities of isoprene synthase. Our findings provide new insight into the molecular mechanisms of isoprene emission diversity in Fagaceae.

## Introduction

Plants emit a variety of volatile organic compounds (VOCs), numbering approximately 30,000 (Peñuelas & Llusià, 2004). These compounds exhibit high chemical diversity, ranging from terpenes and benzenoids to esters and sulfur-containing metabolites. Plants invest heavily in the production of these VOCs, allocating up to 10 % of the carbon fixed by photosynthesis (Peñuelas & Llusià, 2004). This figure strongly suggests that VOCs play important roles in the plant life cycle. For example, they serve as communication tools with other organisms and have been developed during the long evolutionary processes of plants with a sessile lifestyle. In the context of biological interactions, some VOCs attract pollinators and repel herbivores. They are also involved in abiotic stress responses, such as enhancing thermal stress resilience, which contributes to a plant’s ability to adapt to severe environments (Peñuelas & Llusià, 2003; Sasaki et al., 2007).

On the other hand, plant-derived VOCs strongly impact air quality and the climate by forming ozone and aerosols, which create photochemical smog and rain clouds (Kavouras et al., 1998). The global flux of these biogenic VOCs is estimated to be 10^9^ tons of carbon equivalent per year. Most of this flux consists of low-molecular-weight plant-derived terpenes, such as isoprene (C_5_), monoterpenes (C_10_), and sesquiterpenes (C_15_) (Guenther et al., 1995). Isoprene alone represents nearly half of the total biogenic VOCs emitted into the atmosphere. Due to its high chemical reactivity, isoprene’s half-life is measured in hours in the atmosphere. The resulting decomposed compounds form secondary particles (Crounse et al., 2013; Wennberg et al., 2018). These oxidized particles also contribute to the formation of cloud nuclei (Ehn et al., 2014). Due to their significant impact on atmospheric chemistry, detailed information about plant-derived VOC emissions is necessary for improving concrete, reliable climate modeling (Cao et al., 2021).

Isoprene emission has been observed throughout plant lineages, including mosses (Hanson et al., 1999), ferns (Saito & Yokouchi, 2006; Tingey et al., 1987), gymnosperms, and angiosperms (Sharkey et al., 2008). In angiosperm genera, *Populus* (Salicaceae) and *Quercus* (Fagaceae) are representative emitters in the Northern Hemisphere. In contrast, *Eucalyptus* (Myrtaceae) and *Phoenix* (Arecaceae) are known isoprene emitters in tropical dry areas, such as Australia. The grass *Arundo* (Poaceae) is also reported as an active isoprene emitter. Thus, isoprene emitters are widespread.

However, not all plant species can emit isoprene; isoprene non-emitters are the majority among land plants. Isoprene emitters are found in scattered patterns in the land plant phylogeny, and it has been suggested that the ability to emit isoprene has been gained and lost multiple times during evolution (Monson et al., 2013).

In plant cells, isoprene is synthesized from dimethylallyl diphosphate (DMAPP) in a single reaction by isoprene synthase (IspS), which belongs to the terpene synthase (TPS) family. This is the major pathway (Silver & Fall, 1991). Several plant species, including *Pueraria montana* (Sharkey et al., 2005), *Populus alba* (Sasaki et al., 2005), and *Ficus virgata* (Oku et al., 2015), have been identified as having IspSs. IspS proteins are generally localized in plastids. These angiosperm IspS proteins form the primary IspS clade within the TPS-b subfamily, one of eight TPS subfamilies (TPS-A to H) (Sharkey et al., 2013). Additionally, *IspS* genes arising in distinct phylogenetic positions have been reported, consistent with convergent evolution often observed in the TPS family (Ding et al., 2020; Sharkey et al., 2013). For example, a bifunctional myrcene synthase/IspS from *Humulus lupulus* (Cannabaceae) belongs to the TPS-b subfamily, yet it is positioned outside the major *IspS* clade (Sharkey et al., 2013). Furthermore, *IspS* genes belonging to other TPS subfamilies have been reported. In gymnosperms, 2-methyl-3-buten-2-ol synthase from *Pinus sabiniana* produces isoprene even though it belongs to the TPS-d subfamily (Gray et al., 2011). In the moss species *Calohypnum plumiforme*, an *IspS* belonging to the TPS-c type has been reported (Kawakami et al., 2023). The taxonomic diversity of isoprene emitters and the evolutionary diversity of *IspSs* suggest their importance to plant life. However, the molecular details of isoprene-emitting ability remain largely unknown.

The *Quercus* genus (Fagaceae) includes major isoprene emitters in the Northern Hemisphere, such as *Q. serrata* and *Q. crispula* var. *crispula*. Isoprene emission diversity has been extensively studied at the species level within this family (Monson et al., 2013; Tani & Kawawata, 2008). Notably, even within the same genus, there are species that do not emit isoprene, e.g., *Q. glauca* and *Q. phillyraeoides*. A related fagaceous tree, *L. edulis*, does not emit isoprene despite its taxonomic proximity to isoprene emitters. However, *IspS* in Fagaceae remains unidentified, and the TPS family in Fagaceae is poorly understood, with a few exceptions, such as the myrcene synthase gene from *Q. ilex* (Fischbach et al., 2001).

In this study, we identified an *IspS* gene in *Q. serrata* and performed a detailed functional characterization of its gene product. We obtained orthologous *IspS* genes from non-emitter species in the Fagaceae family and, through site-directed mutagenesis studies, identified amino acid substitutions that contribute to the species-specific isoprene-emitting ability in Fagaceae.

## Results

### In silico screening for IspS candidates in Quercus serrata

To identify isoprene synthase (*IspS*) candidates from the isoprene-emitting representative of the Fagaceae family, *Q. serrata*, we screened transcriptome data from its leaves and buds harvested from May to December. These seasonal transcriptome datasets contained 26 *TPS* genes. Among these, three *QsTPS*s (*TPS15-17*) exhibited higher gene expression in summer leaves than in spring and autumn leaves (Fig. 1a). Similar expression patterns were also observed for these *QsTPS*s in buds (see Supplementary Figs. S1-S3). High isoprene emission from *Q. serrata* leaves has been reported in the summer (Ohta, 1986), so these three *QsTPS*s are considered the IspS candidates. In contrast, the TPSs that are preferentially expressed in the winter are predicted to be sesquiterpene synthases. These enzymes are classified in a different clade from TSP-b. These enzymes localize in the cytosol and are responsible for forming C-15 terpenes from the mevalonate pathway. To investigate the relationship between daily isoprene emissions and gene expression further, we analyzed we analyzed the isoprene emissions from *Q. serrata* leaves using GC-MS and *QsTPS15–17* expression using qRT-PCR. Isoprene emission was highest in the afternoon, followed by the morning and nighttime emissions.

**Figure 1.**
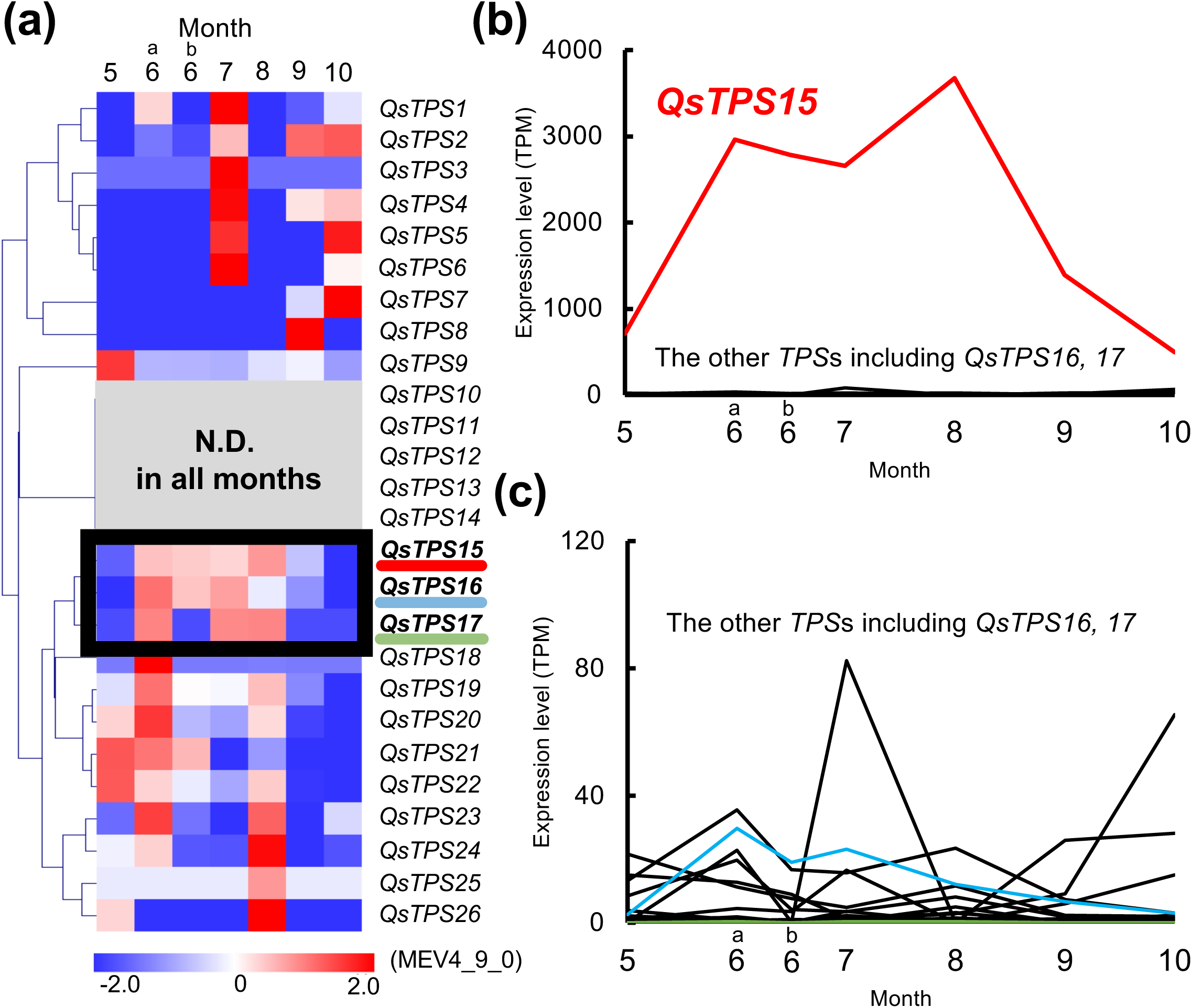
Seasonal changes in the expression pattern of *TPS* genes in *Q. serrata* leaves. (a) Heatmap showing contigs belonging to the *TPS* family based on the seasonal transcriptome dataset of *Q. serrata* leaves (n = 1). The expression of *QsTPS10-14* was undetectable at all data points. N.D., not detected. Samples were collected twice in June: at the beginning (1st) and end (28th). (b) TPM-based expression levels of *QsTPS*s. The individual expression profile of each *TPS* is shown in Supplementary Fig. S2. (c) An enlargement of (b) to highlight the expression of other *TPS*s with *QsTPS*s *16* and *17*.

There were significant differences among the time points (see Supplementary Fig. S4a–c), while monoterpene emissions were an undetectable level in the environmental background. No amplification was detected for *QsTPS16* and *17*. However, *QsTPS15* showed a clear diurnal pattern, with significantly increased expression during the afternoon compared to nighttime (see Supplementary Fig. S4d). These results suggest that the expression level of *QsTPS15* correlates with isoprene emission. Based on its significantly higher expression than the other two *TPS*s in leaves (Fig. 1b,c), we selected this contig as the primary *IspS* candidate, and we isolated its cDNA with full coding sequence (CDS) from RNA prepared from *Q. serrata* leaves.

The predicted polypeptide sequence of QsTPS15 (593 amino acids) contains two aspartate-rich motifs (DDxxD and NSE/DTE) and the N-terminal motif RR(x)8W. These motifs bind to prenyl diphosphate via metal ions (see Supplementary Fig. S5) (Bohlmann et al., 1998; Christianson, 2006). The TargetP2.0 program predicted the presence of a plastid-sorting signal (transit peptide, TP) at the N-terminal region of this candidate (Supplementary Fig. S5). QsTPS15 shows 38-63 % amino acid identity with angiosperm IspSs in the TPS-b subfamily and lower identity with other IspS members: *C. plumiforme* (28 %) in the TPS-c subfamily and *P. sabiniana* (36 %) in the TPS-d1 subfamily (see Supplementary Table S2). QsTPS15 is located close to the angiosperm IspSs in the TPS-b subfamily. It is included in the clade consisting mainly of IspS and ocimene synthase in the phylogenetic tree of the plant TPS family (Fig. 2, Supplementary Fig. S6).

**Figure 2.**
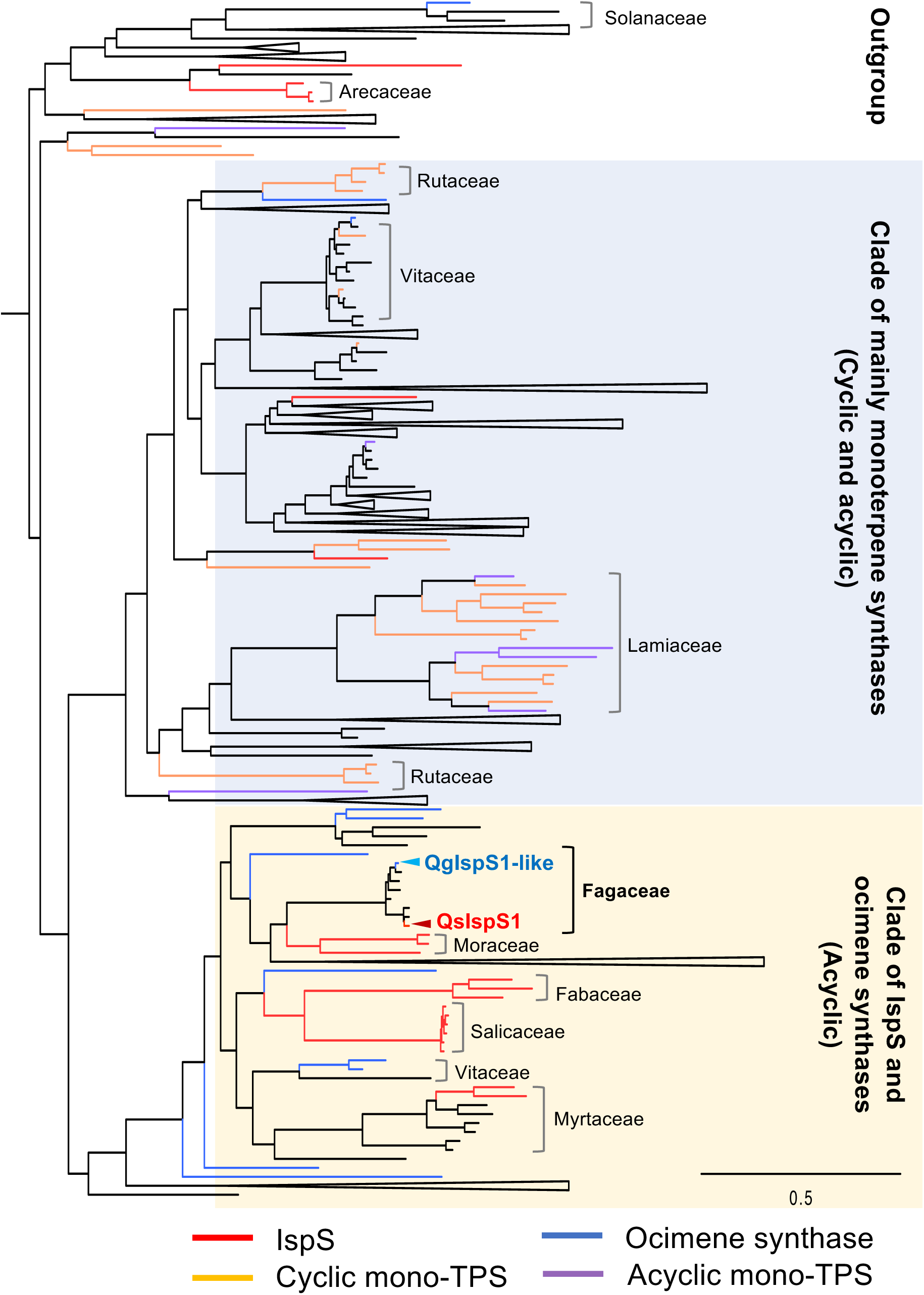
The phylogenetic tree of TPS-b isoprene and monoterpene synthases. A phylogenetic tree of terpene synthases was constructed usig IQ-TREE v2.2.5. Cades close to QsIspS1 are indicated. The IspSs, ocimene synthases, acyclic monoterpene synthases, cyclic monoterpene synthases, and uncharacterized TPSs are highlighted in red, blue, purple, orange, and black, respectively. The branches that include only enzymatically uncharacterized *TPS*s are collapsed. The bar indicates 0.5 substitutions per site. A phylogenetic tree encompassing all plant TPSs is shown in Supplementary Fig. S6.

A detailed comparison of angiosperm TPS amino acid sequences highlighted four amino acid residues (phenylalanine at the 1st site, serine or valine at the 2nd site, phenylalanine at the 3rd site, and asparagine at the 4th site) (see Supplementary Fig. S5). These residues are located close to the catalytic pocket and are frequently present in angiosperm IspSs, including those that have evolved independently (Bergman & Dudareva, 2024; Li et al., 2017; Sharkey et al., 2013). The conservation of these residues is designated the “isoprene score,” which can be used to estimate IspS activity. QsTPS15 achieves a maximum score of 4 based on the number of matching amino acid sites.

### Detection of the isoprene synthase activity of QsTPS15

We first assessed the enzymatic activity of QsTPS15 using a recombinant protein that was transiently expressed in *Nicotiana benthamiana* via agroinfiltration. We used the “Tsukuba system” vector, pTKB3, which enables the production of high levels of exogenous proteins in plant hosts due to a geminiviral replication system and a double terminator (Nozaki et al., 2021). Cell-free extract was prepared from transformed *N. benthamiana* leaves where the recombinant QsTPS15 protein was produced. This extract was then used as crude enzymes for an in vitro IspS assay. After incubating the crude enzymes with DMAPP as the substrate and MgCl₂ as the cofactor in a glass tube, the volatiles in the headspace were trapped using solid-phase microextraction (SPME). The collected volatiles were introduced to GC/MS, and the analyzed data indicated that QsTPS15 produced a clear enzymatic reaction product. Its retention time and MS spectra were identical to those of the isoprene standard (Fig. 3a, b).

**Figure 3.**
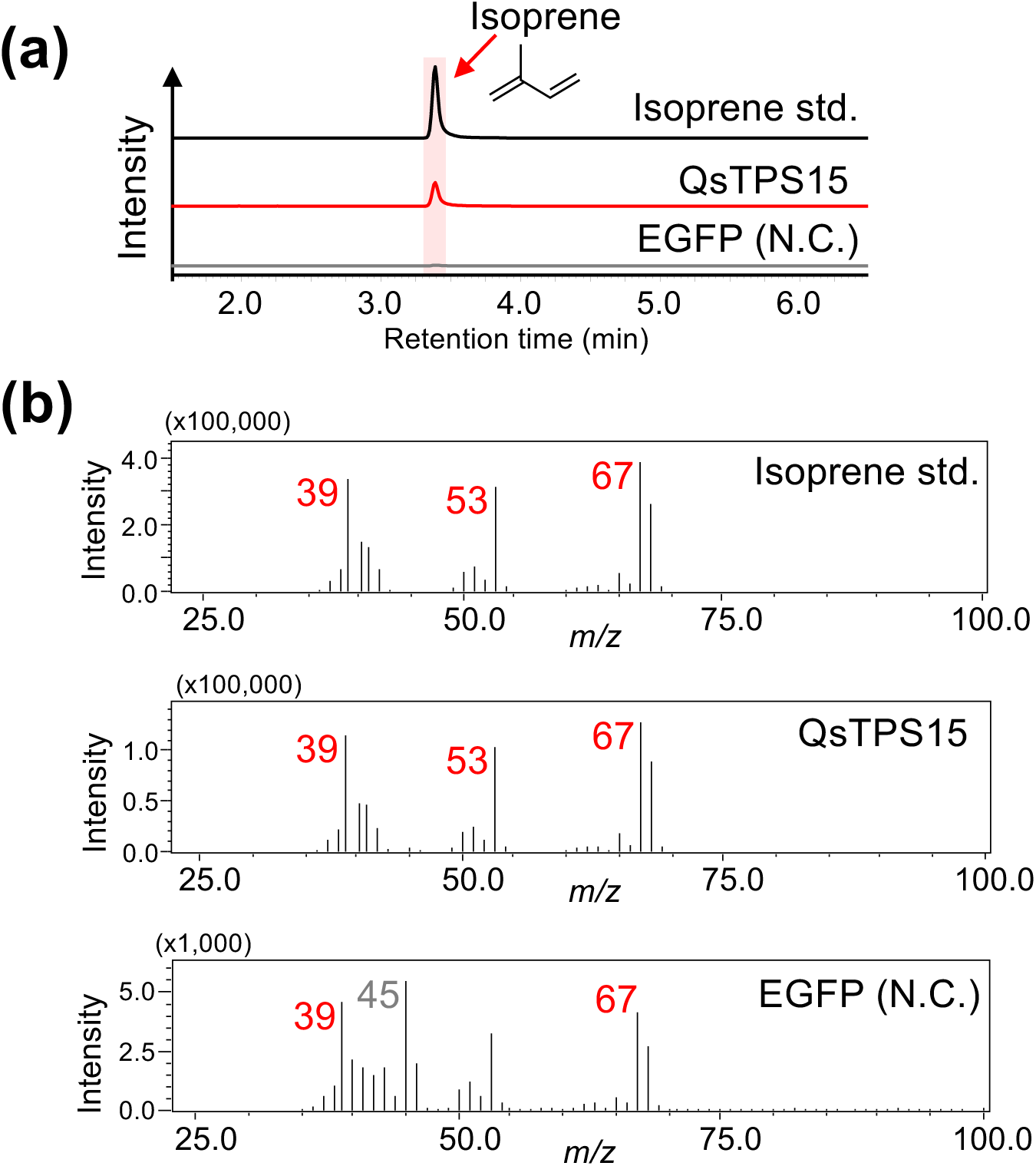
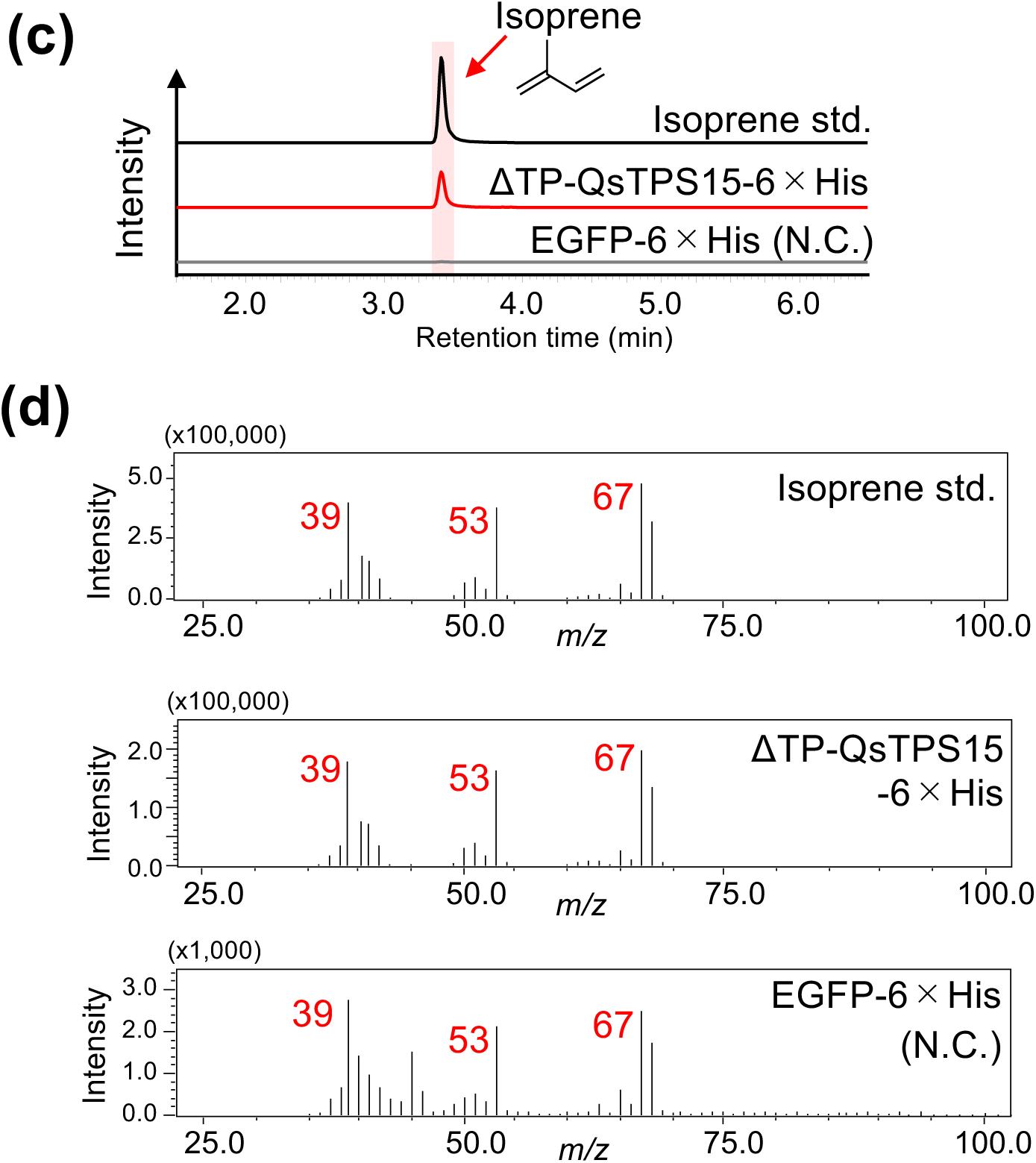
Detection of the enzyme activity of recombinant QsTPS15. (a) GC-MS chromatogram (*m/z* = 67) and (b) MS spectrum (retention time = 3.4 min) of an enzyme reaction mixture using recombinant QsTPS15 protein produced by *N. benthamiana* as a host plant. EGFP was used as a negative control. (c) MS chromatogram (*m/z* = 67) and (d) MS spectrum (retention time = 3.4 min) of an IspS reaction mixture containing recombinant ΔTP-QsTPS15-6×His protein produced by *E. coli*. EGFP-6×His was used as a negative control.

We also performed the in vitro characterization of QsTPS15 in an *Escherichia coli* expression system to facilitate further experimentation. We modified QsTPS15 by removing the N-terminal region containing the putative TP (ΔTP) and adding a 6×His tag at the C-terminus for affinity purification. The recombinant ΔTP-QsTPS15-6×His protein was produced in *E. coli* using a pET-based expression system and purified via Ni-NTA affinity chromatography (see Supplementary Fig. S7). The purified *E. coli* recombinant protein with the histidine tag, ΔTP-QsTPS15-6×His, exhibited identical IspS activity to the recombinant protein from the plant host in the same manner described above (Fig. 3c, d). We performed various negative control assays using the purified recombinant protein, including incubations without crude enzyme, DMAPP, or MgCl₂, as well as incubations with heat-denatured enzyme. As a result, a trace amount of isoprene, most likely formed non-enzymatically from DMAPP, was detected; however, clear isoprene formation was observed in the full assay (Fig. 4, Supplementary Fig. S8). These biochemical results from the plant and bacterial expression systems indicate that the recombinant QsTPS15 exhibits IspS activity.

**Figure 4.**
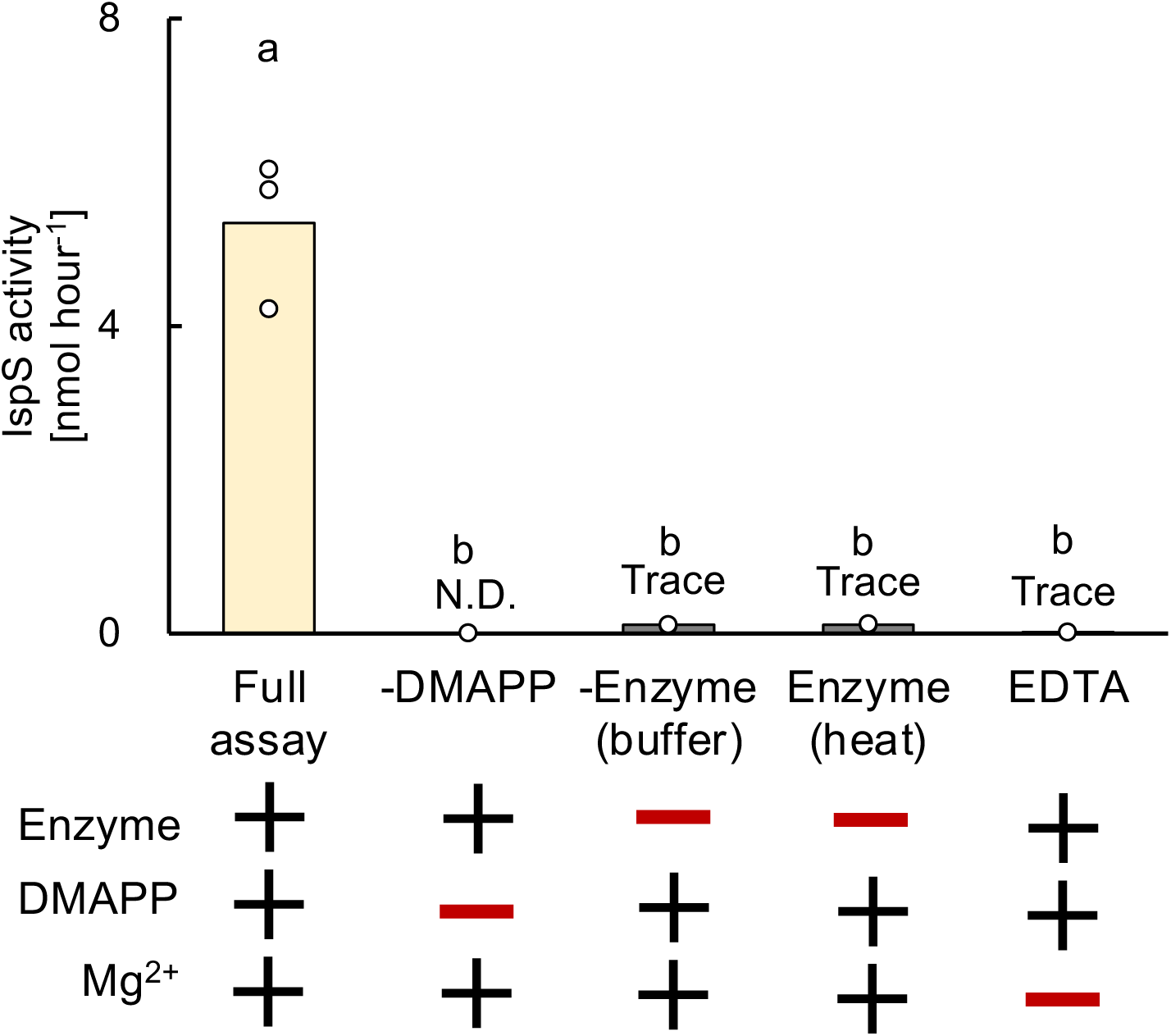
Isoprene synthase assay with QsTPS15. Full and various negative control assays of the recombinant ΔTP-QsTPS15-6×His protein. From left to right: full assay; assays without DMAPP; with reaction buffer instead of enzymes; with heat-denatured enzymes at 95 °C for 20 min; and with excessive EDTA instead of MgCl₂. N.D., not detected. Bars indicate the mean of biological replicates, and dots represent the values of each replicate. Statistical analysis was performed using the Tukey-Kramer test. Different letters indicate statistically significant differences (n = 3).

### Enzymatic characterization of QsTPS15

The substrate specificity of ΔTP-QsTPS15-6×His was evaluated using various prenyl diphosphates of different chain lengths as substrates, such as geranyl diphosphate (GPP: C_10_) and farnesyl diphosphate (FPP: C_15_) (Fig. 5). Using GPP as the substrate, only a trace amount of product was detected with ΔTP-QsTPS15-6×His. Direct comparison with a standard specimen identified the product as β-ocimene, and a similarity search based on the MS spectrum predicted the reaction product to be the cis isomer. However, the yield was much lower than that of isoprene when DMAPP was used as the substrate. With FPP as the substrate, no detectable product was obtained. These results suggest that QsTPS15 primarily functions as an isoprene synthase in *Q. serrata*. Thus, we renamed this enzyme QsIspS1.

**Figure 5.**
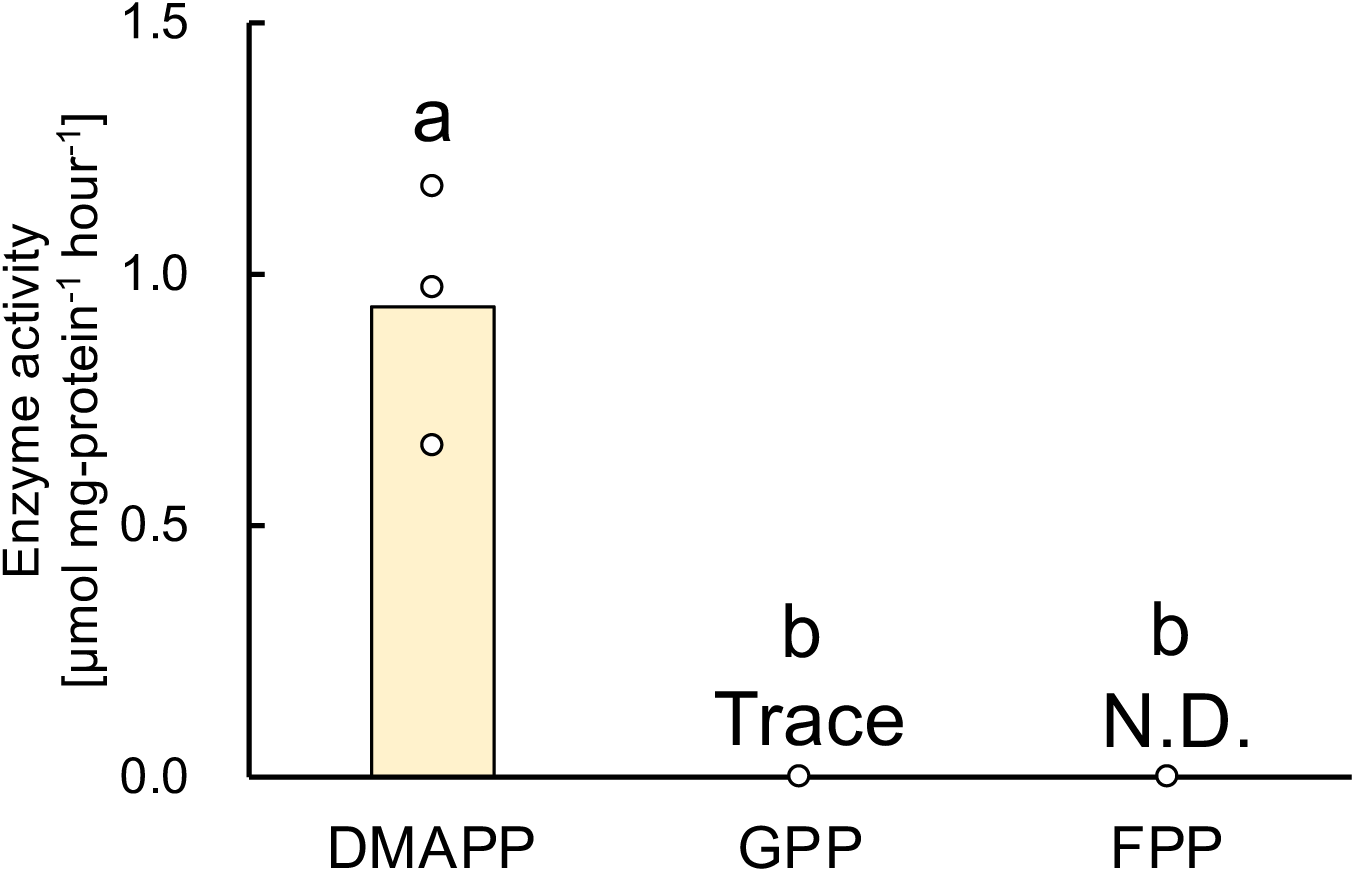
Substrate specificity of QsTPS15. Enzymatic activity of ΔTP-QsTPS15-6×His with prenyl diphosphate substrates of different chain lengths; DMAPP (C_5_), GPP (C_10_), and FPP (C_15_). N.D., not detected. Bars indicate the mean of biological replicates, and dots represent the values of each replicate. Statistical analysis was performed using the Tukey-Kramer test. Different letters indicate statistically significant differences (n = 3).

We conducted a further kinetic analysis. In this study, we use the *K*_½_ value instead of the *K*_m_ value for isoprene synthase because a possible substrate inhibition could not be evaluated due to the extremely high DMAPP concentration in the enzyme assay. The *K*_½_ value is half of the maximum observed rate. The *K*_1/2_ value, the *k*_cat_ value, and the *k*_cat_ *K*_1/2-1_ value of ΔTP-QsIspS1-6×His for DMAPP were calculated to be 12 ± 9 mM, 30 ± 20 min^-1^, and 3 ± 3 min^-1^ mM^-1^, respectively (Table 1, Supplementary Fig. S9). This *k*_cat_ *K*_1/2-1_ value falls within the range reported in previous studies, i.e., *F. septica* IspS (0.2 min^-1^ mM^-1^), *P. alba* IspS (0.2 min^-1^ mM^-1^), *P. montana* IspS (0.7 min^-1^ mM^-1^), *C. equisetifolia* IspS (3 min^-1^ mM^-1^), *P. tremuloides* IspS (13 min^-1^ mM^-1^), and *E. globulus* IspS (59 min^-1^ mM^-1^) (Oku et al., 2015).

**Table 1.** Kinetic parameters of QsIspS1 of *Q. serrata*.

| $k_{\text{cat}}$ (min<sup>-1</sup>) | $K_{1/2}$ (mM) | $k_{\text{cat}} K_{1/2}^{-1}$ (min<sup>-1</sup>mM<sup>-1</sup>) |
| --- | --- | --- |
| $30 \pm 20$ | $12 \pm 9$ | $3 \pm 3$ |

### Subcellular localization of QsIspS1

The online prediction program TargetP 2.0 estimated the plastid localization of the QsIspS1 protein. To confirm this experimentally, a green fluorescent protein (GFP) was fused to the C-terminus of the full QsIspS1 polypeptide sequence (QsIspS1-GFP) and to the partial N-terminal sequence (i.e., the first 40 amino acids containing the predicted transit peptide [TP]) (QsIspS1-TP-GFP). These chimeric proteins were transiently produced in epidermal cells of *N. benthamiana* leaves via agroinfiltration. Confocal microscopic analysis revealed that the GFP fluorescence of both chimeric proteins was present in dotted structures that matched the localization of chlorophyll autofluorescence but differed from the localization of free sGFP, which highlighted the nucleus and cytosol (Fig. 6). These results strongly suggest that QsIspS1 functions in plastids in *Q. serrata* and utilizes DMAPP from the MEP pathway.

**Figure 6.**
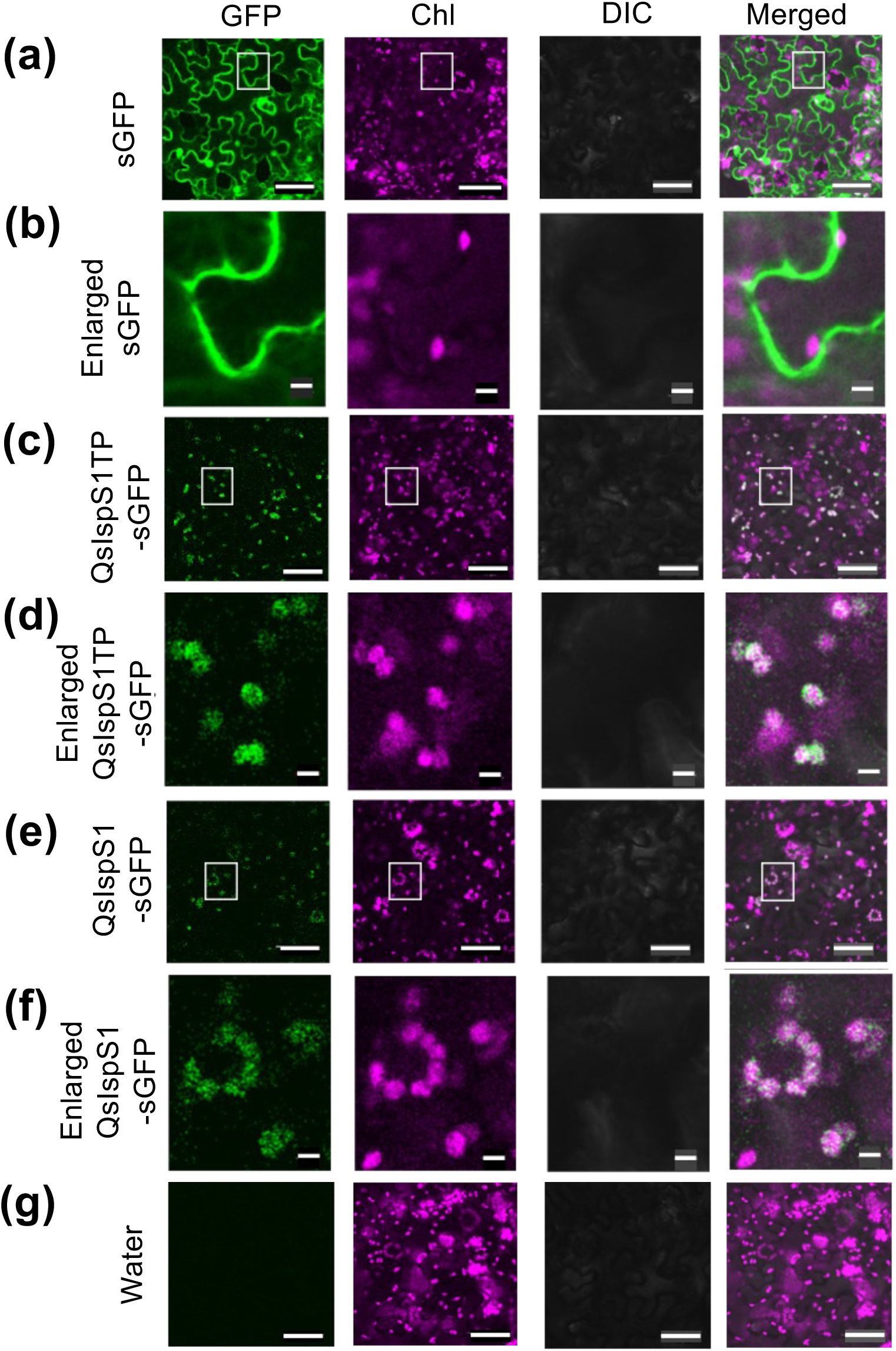
Subcellular localization of QsIspS1. Microscopic observation of the transit peptide of *Q. serrata* isoprene synthase 1 (QsIspS1TP) and the synthetic green fluorescent protein (sGFP), which were expressed in epidermal cells of *N. benthamiana*. sGFP (a), QsIspS1TP-sGFP (c), and QsIspS1-sGFP (e) were transiently expressed in *N. benthamiana* leaves via agroinfiltration. Water infiltrated into the leaves served as the negative control (g). The rows from the top show images of GFP signaling, chlorophyll autofluorescence, differential interference contrast (DIC), and merged images. White squares indicate enlarged areas. Enlarged images (b, d, f) are shown for sGFP (a), QsIspS1TP-sGFP (c) and QsIspS1-sGFP (e). Bars: 50 µm (a, c, e, g); 5 µm (b, d, f). More than three cells were observed per leaf, and three leaves were examined per individual. Observations were performed on two biological replicates.

### Characterization of *IspS*-like genes in isoprene non-emitter species

To gain insight into the molecular basis of isoprene emission diversity in Fagaceae species, we searched the genome sequences of two non-emitting Fagaceae trees, *L. edulis* and *Q. glauca*, using *QsIspS1* as the query (Kudo et al., 2025). Using TPS domains as a guide, we identified TPSs and found the genes most similar to *QsIspS1* in the genomes of the two non-emitter plants. These genes are *LeIspS1-like* and *QgIspS1-like*. LeIspS1-like and QgIspS1-like share 94 % and 93 % amino acid identity with QsIspS1, respectively. While the nucleotide lengths of these cDNAs is the same (1827 bp), we discovered that LeIspS1-like had a nucleotide substitution that caused an in-frame stop codon, resulting in a truncated polypeptide of 169 amino acids (Fig. 7a, Supplementary Figs. S10 and S11). Since this truncated polypeptide lacks the essential substrate-binding motifs for enzymatic function, it is likely nonfunctional as an IspS.

**Figure 7.**
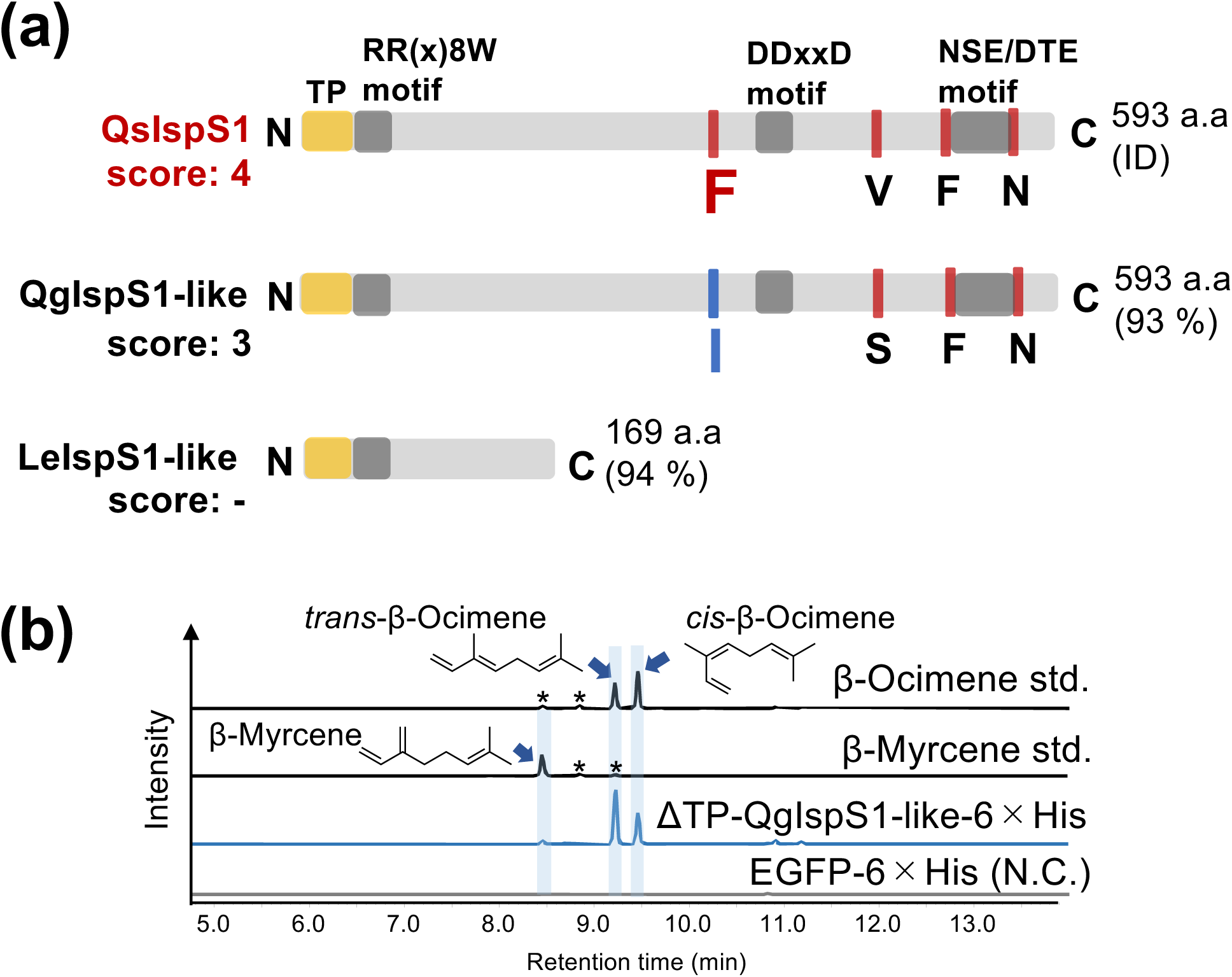
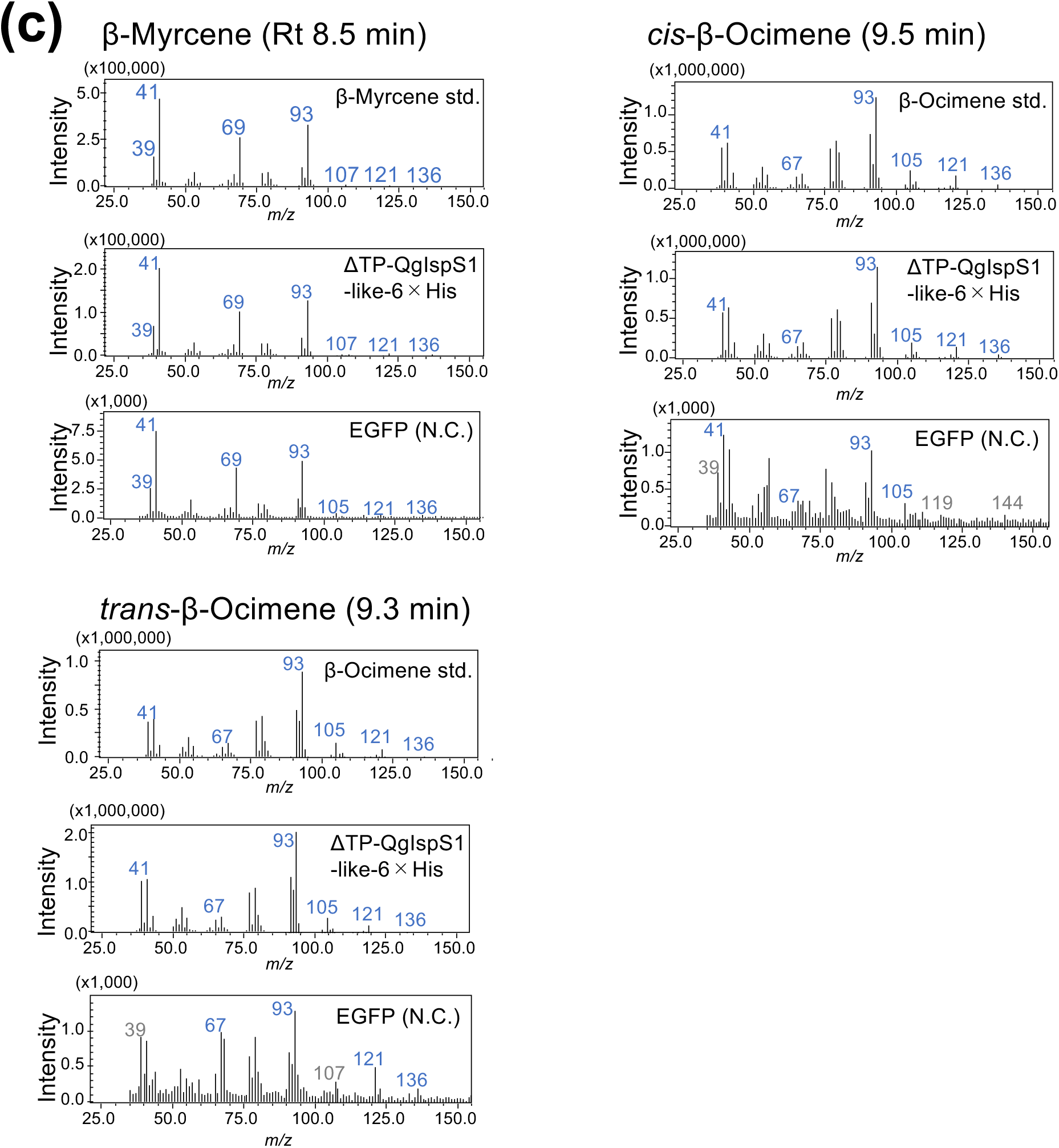
Detection of enzyme activity of QgIspS1-like protein of *Q. glauca*. (a) Comparison of amino acid (aa) sequences of QsIspS1, QgIspS1-like, and LeIspS1-like. TP: transit peptide. (b) The MS chromatogram (*m/z* = 93) and (c) the MS spectrum (retention time = 8.5, 9.3, and 9.5 min) of the reaction mixture of the recombinant ΔTP-QgIspS1-like-6×His protein produced by *E. coli*. EGFP-6×His was used as a negative control. MS chromatogram (*m/z* = 93). The asterisks (*) indicate impurity peaks.

**Figure 8.**
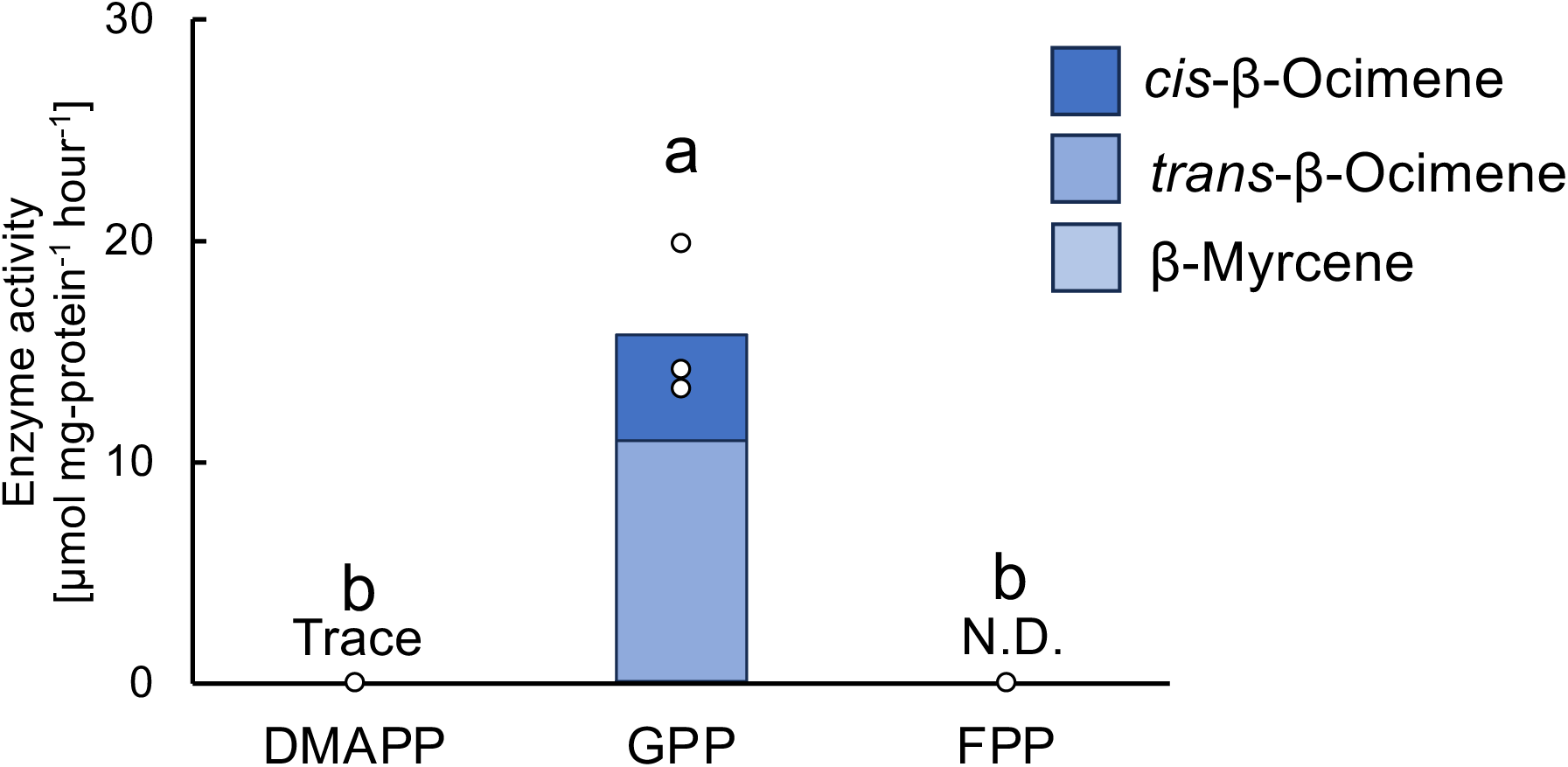
Substrate and product specificity of QgIspS1-like protein. The substrate preference of the QgIspS1-like protein is expressed as ΔTP-QgIspS1-like-6×His. The substrates used were DMAPP (C_5_), GPP (C_10_), and FPP (C_15_). The enzyme reaction products were quantified by GC-MS analysis. N.D., not detected. Bars indicate the mean of biological replicates, and dots represent the values of each replicate. Statistical analysis was performed using the Tukey-Kramer test. Different letters indicate statistically significant differences (n = 3).

Although QgIspS1-like does not have a substitution resulting in the clear loss of IspS function, the above-mentioned isoprene score was 3. In this case, the phenylalanine at the first position of the isoprene score site was replaced with isoleucine (Fig. 7a). Thus, we prepared the recombinant ΔTP-QgIspS1-like-6×His protein and evaluated its enzyme activity biochemically using the purified protein. Analysis of substrate specificity using DMAPP, GPP, and FPP as substrates showed that this enzyme is dedicated to synthesizing the monoterpenes β-myrcene, *cis*-β-ocimene, and *trans*-β-ocimene (Figs. 7b, c, and 8), with *trans*-β-ocimene being the main product. Additionally, we determined the kinetic parameters of the ΔTP-QgIspS1-like-6×His protein (Table 2, Supplementary Figs. S12, S13). These results indicate that QgIspS1-like functions differently as a monoterpene synthase using GPP as its substrate. We then analyzed the VOC emission from *Q. glauca* and found that major VOCs were that α-pinene, β-pinene, and *trans*-β-ocimene, which were identified with standard samples. Other peaks detected in the GC-MS analysis were annotated as α-phellandrene and (+)-sabinene. Isoprene was, however, undetectable at background levels. We also investigated the correlation between total ocimene emissions and *QgIspS1-like* expression. While there were no statistically significant differences in emission levels or gene expression, both tended to peak in the afternoon (Supplementary Fig. S14c, d). Interestingly, a strong correlation was observed between the ocimene emissions and the *QgIspS1-like* expression in the afternoon, with an R² value of 0.98. In contrast, other monoterpenes, such as α-pinene and β-pinene, exhibited only a weak correlation with *QgIspS1-like* expression (Supplementary Fig. S15).

**Table 2.** Kinetic parameters of QgIspS1-like of *Q. glauca*.

| | $k_{\text{cat}}$ (min<sup>-1</sup>) | $K_{\text{m}}$ (μM) | $k_{\text{cat}} K_{\text{m}}^{-1}$ (min<sup>-1</sup> μM<sup>-1</sup>) |
| --- | --- | --- | --- |
| β-myrcene | 1.9 ± 0.2 | 220 ± 60 | 0.009 ± 0.002 |
| *cis*-β-ocimene | 29 ± 2 | 190 ± 40 | 0.15 ± 0.03 |
| *trans*-β-ocimene | 65 ± 5 | 180 ± 40 | 0.35 ± 0.08 |

### Site-directed mutagenesis of QsIspS1 and QgIspS1-like proteins

We conducted a site-directed mutagenesis study with these two enzymes to investigate whether the functional difference between QsIspS1 and QgIspS1-like is attributable to the single amino acid substitution at the first isoprene score site (326th residue). We constructed two reciprocal mutants: phenylalanine at position 326 was substituted with isoleucine in QsIspS1 (ΔTP-QsIspS1(F326I)-6×His), which decreased the isoprene score to 3; and isoleucine at position 326 was substituted with phenylalanine in QgIspS1-like (ΔTP-QgIspS1-like(I326F)-6×His), which increased the isoprene score to 4. We prepared the recombinant proteins of these mutants in an *E. coli* expression system and analyzed them biochemically to compare them with the corresponding wild-type enzymes. The F326I mutation in QsIspS1 significantly reduced IspS activity, generating monoterpene synthase activity and yielding *cis*-and *trans*-β-ocimene (Fig. 9a, b). These results demonstrate that the F326I amino acid substitution shifts the catalytic function of IspS to monoterpene synthase activity (see Supplementary Fig. S16a). Conversely, the I326F point mutation in QgIspS1-like significantly reduced its monoterpene synthase activity (Fig. 9c, d), though this mutant enzyme retains monoterpene synthase activity as its primary function (Supplementary Fig. S16b). Additionally, the IspS activity of QgIspS1-like (I326F) showed an increasing trend, though the difference was not statistically significant. These results suggest that the 1st isoprene score site is important for expressing the IspS activity of QsIspS1 and the monoterpene synthesis activity of QgIspS1-like.

**Figure 9.**
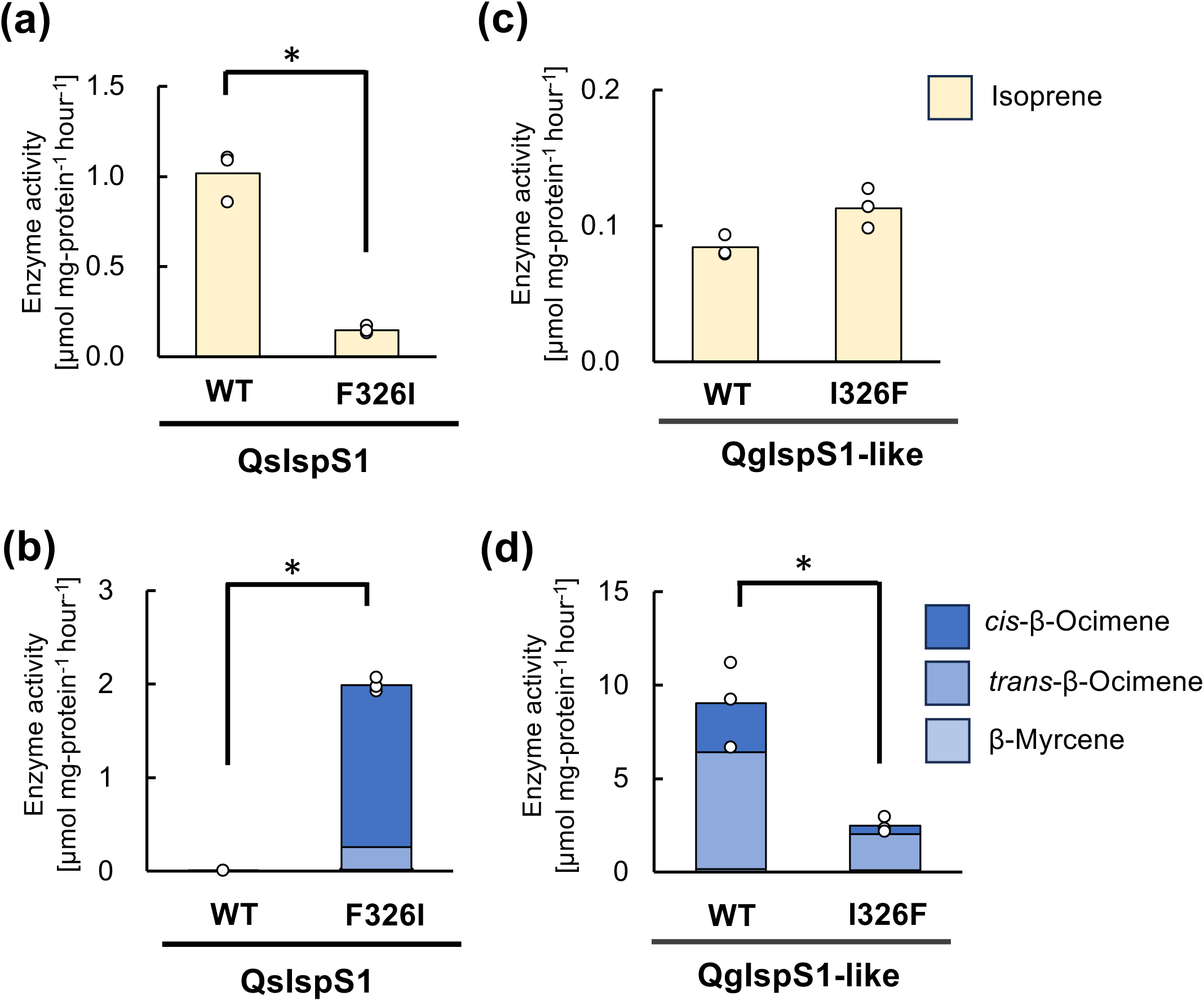
Enzymatic properties of mutant proteins of QsIspS1 and QgIspS1-like. Enzymatic properties of the wild-type and mutant (F326I) forms of QsIspS1 with DMAPP (a) and GPP (b) as substrates. Enzymatic properties of the wild-type and mutant (I326F) QgIspS1-like proteins with DMAPP (c) and GPP (d) as substrates. Bars indicate the mean of biological replicates, and dots represent the values of each replicate. Statistical analysis was performed using a Welch’s t-test (*p*\* < 0.05, n = 3). The asterisk indicates a statistically significant difference.

## Discussion

Isoprene, the most abundant biogenic VOC emitted by a variety of plant species, including trees, herbs, ferns, and mosses, is released into the atmosphere. The total amount of isoprene emitted globally is 0.5 G-ton of carbon. Isoprene emission is intensively studied in the field of atmospheric chemistry because of its strong impact on atmospheric quality. In plant physiology research, independent working groups have studied the physiological and biological roles of isoprene emission for a long time. These studies show that isoprene emission is involved in resistance against various abiotic stresses (Ghirardo et al., 2021; Jud et al., 2016; Loreto & Velikova, 2001; Monson et al., 2021; Sasaki et al., 2007; Sharkey & Singsaas, 1995). Isoprene emission is primarily driven by the IspS function. *IspS* genes have been identified in many plant taxa, but interestingly, they have different evolutionary origins. The widespread occurrence of isoprene emission and the various evolutionary origins of *IspS*s may provide insight into the direct physiological roles of isoprene.

However, isoprene emission is not essential for plant life, as many plant species do not emit isoprene. From a taxonomic point of view, isoprene-emitting plants are scattered. Within one plant genus, some species emit isoprene while others do not. It should be noted that the molecular mechanisms explaining why some closely related plant species exhibit such diversity in their ability to emit isoprene remain unknown. In the present study, we focused on the diversity of isoprene emission in the Fagaceae family, which is one of the major isoprene emitters in the Northern Hemisphere. Since information on *Quercus* isoprene synthase is limited (Li et al., 2016a, 2016b, 2016c), we first identified *IspS* from *Quercus* species, and a detailed analysis of the genetic and biochemical features of the *Quercus IspS* gene was performed.

As a prominent characteristic of angiosperm IspS, this enzyme family belongs to the TPS-b subfamily and exhibits low affinity for its substrate, DMAPP. These enzymes are commonly localized in plastids and are monofunctional, producing isoprene as the sole reaction product. However, some IspSs are exceptional. For example, IspSs of *E. globulus* and *Casuarina equisetifolia* have a higher affinity than most angiosperm IspSs (Oku et al., 2015; Sharkey et al., 2013), and *H. lupulus* has a bifunctional myrcene synthase/IspS. Despite belonging to the TPS-b subfamily, the hop enzyme positions outside the major IspS clade (Sharkey et al., 2013). Phylogenetic analysis and enzymatic characterization revealed that *QsIspS1* belongs to the major *IspS* clade in the TPS-b subfamily. As seen in Fig. 2, the phylogenetic tree shows that the clade containing major IspSs and acyclic ocimene synthases diverges from the clade of cyclic and acyclic representative monoterpene synthases in the TPS-b subfamily. QsIspS1 is included in the same clade as many IspSs and ocimene synthases. The presence of IspSs in multiple branches also suggests that IspS was independently acquired from ocimene synthase through convergent evolution among different plant families.

*QsIspS1* primarily uses DMAPP as its substrate and has a low affinity for DMAPP, with a *k*cat *K*_1/2-1_ of 3 ± 3 min⁻¹ mM⁻¹. Subcellular localization analysis showed that QsIspS1 is localized in the plastid. This suggests that QsIspS1 uses DMAPP provided by the MEP pathway. Some Fagaceae species produce various monoterpenes, sesquiterpenes, and isoprene (Staudt & Bertin, 1998; Welter et al., 2012), but molecular biological studies have not yet been conducted, and only one myrcene synthase gene has been identified in Fagaceae thus far (Fischbach et al., 2001). Our transcriptome analysis revealed 25 *TPS* genes, including a close homolog of the reported myrcene synthase, in addition to *QsIspS1*. Further characterization of these genes will improve our understanding of VOC production in this species at the molecular level.

We identified TPS genes with high sequence similarity to *QsIspS1* in the genomes of two isoprene non-emitting species, *L. edulis* and *Q. glauca*. These genes were named *LeIspS1-like* and *QgIspS1-like*, respectively. Their nucleotide sequence identities to *QsIspS1* were 97 % and 96 %, respectively. However, *LeIspS1-like* contains a stop codon in the middle of the open reading frame (ORF) due to a substitution. These results in a truncated translation product that lacks amino acid sequences critical for catalytic activity, i.e., two aspartate-rich motifs. These findings strongly suggest that the *LeIspS1-like* gene product does not function as an IspS or even a TPS, indicating that it is a pseudogene. The incompleteness of the CDS is the main reason why *L. edulis* does not emit isoprene. In contrast, the predicted QgIspS1-like amino acid sequence of *Q. glauca* has an amino acid substitution at the first site of the isoprene score: phenylalanine is substituted with isoleucine. This substitution is one of the four highly conserved amino acid residues found in plant IspS. This substitution suggests that QgIspS1-like has a different enzymatic function than QsIspS1. Indeed, our functional analysis of the QgIspS1-like recombinant protein revealed that it mainly uses GPP as a substrate to produce monoterpenes, such as *cis*-and *trans*-β-ocimene. Almost no products were produced when DMAPP or FPP were used as substrates. Additionally, the *K*_m_ value of the QgIspS1-like protein was in the same μM range (10–200 μM) as those reported for other monoterpene synthases (Ashaari et al., 2020; Fischbach et al., 2001; Kampranis et al., 2007). The *k*_cat_ *K*_m-1_ values for the QgIspS1-like protein in the production of *cis*-and *trans*-β-ocimene were compared. The *k*_cat_ *K*_m-1_ values for the synthesis of *cis*-β-ocimene and *trans*-β-ocimene were 1.5×10^-1^ ± 3.0×10^-2^ μM⁻¹ min⁻¹ and 3.5×10^-1^ ± 8.0×10^-2^ μM⁻¹ min⁻¹, respectively. These values were more than one order of magnitude higher than the 9.0×10^-3^ ± 2.0×10^-3^ μM⁻¹ min⁻¹ observed for β-myrcene formation catalyzed by this enzyme. These results suggest that QgIspS1-like does not produce isoprene, but rather, is involved in ocimene biosynthesis, unlike QsIspS1. A comparison of the VOC emission profiles of *Q. serrata* and *Q. glauca in vivo* revealed that *Q. serrata* emits only isoprene as the only VOC molecule and does not emit monoterpenes, as reported by Tani and Kawawata (2008) and confirmed in this study (Supplementary Fig. S4a-c). In contrast, *Q. glauca* is known to emit only monoterpenes (Tani et al., 2024), including ocimenes (Supplementary Fig. S14a-c). The expression levels of *QsIspS1* and *QgIspS1-like* were found to correlate with the diurnal emission patterns of isoprene and ocimenes, respectively (Supplementary Figs. S4d, S14d). Notably, the substantial variation observed in ocimene emission among individuals was attributed to the expression level of *QgIspS1-like* in each individual. Overall, QsIspS1 and QgIspS1-like appear to be the primary enzymes that determine the terpenoid VOC emission profiles in *Q. serrata* and *Q. glauca*, respectively.

The amino acid substitution of F326I observed in *Q. glauca* in the present study is probably a natural difference that occurred during evolution. The importance of phenylalanine at this position was reported in a site-directed mutagenesis study of IspS in *Arundo donax* (Poaceae) evaluated in *Arabidopsis thaliana* as the host plant (Li et al., 2017). In their report, they observed a significant decrease in isoprene formation when phenylalanine was substituted with isoleucine, similar to our findings. However, a striking difference was observed in the monoterpene formation ability of the mutant enzyme. In the case of *A. donax* IspS, a significant increase in monoterpene formation was observed when phenylalanine at the 310th position of AdIspS (the first position of the isoprene score) was substituted with alanine. In our study, however, a significant increase in monoterpene synthesis was observed when phenylalanine was substituted with isoleucine, as demonstrated with QsIspS1 (Fig. 9).

In summary, these data suggest that the catalytic mechanisms differ depending on the IspS molecule derived from different plant species. When a mutation (I326F) was introduced to QgIspS1-like in *Q. glauca*, the ability to biosynthesize isoprene was not restored to the same extent as in QsIspS1. Since the mutant enzyme QgIspS1-like (I236F) met the isoprene score criteria, it is hypothesized that, in addition to the four proposed amino acids, other amino acids outside this region contribute to isoprene biosynthesis activity. Furthermore, a significant decrease in monoterpene biosynthesis activity was observed when phenylalanine was substituted with isoleucine in the QgIspS1-like protein (Fig. 9). Together with the increased monoterpene production in the QsIspS1(F326I) mutant, these results suggest that phenylalanine at site 1 is involved in both isoprene and monoterpene biosynthesis. Artificial mutations of a single amino acid in monoterpene synthases have recently been reported to affect product yield and substrate specificity, as in limonene and linalool synthases and sesquiterpene synthases, such as germacradien-4-ol synthase (González Requena et al., 2024; Schiff & Oprian, 2023; T et al., 2023). Together, these findings strongly suggest the high plasticity of the catalytic function of TPS proteins.

In nature, plant VOCs, of which terpenoids are the majority, play various ecological roles, such as attracting pollinators, avoiding enemies, and facilitating tritrophic interactions (e.g., attracting an enemy’s enemy), as well as more complex plant-plant interactions (Takabayashi & Shiojiri, 2019). While the role of isoprene in abiotic stresses in emitter plants is often documented, recent studies have revealed an additional ecological function of isoprene, i.e. it induces resistance against microbial pathogens in neighboring plants (Frank et al., 2021; Zhu et al., 2025). It should be noted that these functions are supported by VOC emission, while the VOC release mechanism is another area of research. Several different modes have been proposed for the mechanism involved in VOC release: transporter-mediated efflux, the vesicle trafficking mechanism, the promotion of secretion by lipid transfer proteins, and direct membrane-membrane interaction (Widhalm et al., 2015). Transporters of different families have been studied extensively in the context of secondary metabolites (Yazaki, 2005, 2006). Notably, members of the ATP-binding cassette transporter family involved in flower scent emission have recently been published (Adebesin et al., 2017; Chang et al., 2023). Despite these examples, the emission mechanisms of the wide variety of terpenoid VOCs are largely unknown. Only isoprene is thought to be released without special molecular machinery, such as transporter proteins. Further intensive studies on VOC emission mechanisms are expected in the future.

The findings of this study have significant implications for atmospheric chemistry (Peñuelas & Llusià, 2004; Satake et al., 2024). The climatic impact of biogenic volatile organic compounds (VOCs) remains highly uncertain (Szopa et al., 2021), in part due to difficulties in estimating their global emissions from terrestrial vegetation (DiMaria et al., 2023). Our study suggests that minor amino acid substitutions in isoprene synthase genes determine whether oak species (*Quercus*) function as isoprene or monoterpene emitters or non-emitters. While more studies are needed, our findings will provide a molecular basis for predicting the biogenic VOC emission potential of *Quercus* species for which emission data are unavailable. This includes savannah and shrub oaks, which remain understudied despite their ecological dominance and the potential for within-genus variation in isoprene emission (Guenther et al., 2020). Addressing a critical gap in emission inventories used in atmospheric models will ultimately enhance model accuracy and reduce uncertainties in the interactions between biogenic VOCs and climate change.

## Materials and methods

### Plant materials and reagents

For RNA-seq analysis and cloning, three leaves and leaf buds of *Q. serrata* were collected from a single tree in the biodiversity reserve field at the Ito campus of Kyushu University, Fukuoka, Japan (33°35′ N, 130°12′ E, 20 to 57 m a.s.l). The sampling date are; 2017/5/3, 6/1, 6/28, 7/26, 8/24, 9/20, 10/18 for leaf and 2017/5/3, 6/1, 6/28, 7/26, 8/24, 9/29, 10/18, 11/15, 12/13, 2018/1/14, 2/8, 3/8 for buds. Samples were taken from the sun-exposed crown (approximately 2–5 m from the ground) using long pruning shears between 11:30 a.m. and 12:30 p.m. The study sites for *Q. glauca* are the biodiversity reserve on the Ito campus of Kyushu University (33°35′ N, 130°12′ E, 20 to 57 m a.s.l). *L. edulis* was studied at the Imajuku Field Activity Center (33°33′ N, 130°16′ E, 84 to 111 m a.s.l). Immediately after harvest, 0.2–0.4 g of tissue from each sample was preserved in a 2-ml microtube containing 1.5 ml of RNA-stabilizing reagent (RNAlater; Thermo Fisher Scientific, Waltham, MA, USA). The samples were transferred to the laboratory within two hours of sampling.

They were stored at 4 °C overnight and stocked at -80 °C until RNA extraction. During transport to the laboratory, the samples were kept on ice in a cooler box to maintain a low temperature. For qRT-PCR, six individuals of *Q. serrata* and *Q. glauca* were purchased from Matsui Nouen (Higashiomi, Japan), and grown at Uji campus of Kyoto University (34°35′ N, 135°48′ E, 23 m a.s.l). The trees were 3 years old and each was 80–100 cm tall.

The substrate compounds such as DMAPP, GPP, and FPP were purchased from Sigma-Aldrich (St. Louis, US), and DMAPP was also purchased from TargetMol (Boston, US). Standard compounds, such as isoprene, β-myrcene, (1R)-(+)-α-pinene, and (-)-β-pinene were purchased from Tokyo Chemical Industry Co., Ltd. (Tokyo, Japan). β-Ocimene was obtained from Cayman Chemical (Ann Arbor, USA). This β-ocimene standard was a mixture of *trans*-and *cis*-β-ocimene.

### Construction of transcriptome data from *Q. serrata*

RNA was extracted from bud and expanded leaf samples of each tree independently. The total RNA from the leaf samples was extracted according to the method described in a previous study (Miyazaki et al., 2014). The total RNA from the bud samples was extracted using PureLink™ Plant RNA Reagent (ThermoFisher Scientific) and was subsequently processed with DNase using the TURBO DNA-free Kit (ThermoFisher Scientific), according to their protocols. The RNA was then purified using the RNeasy Plant Mini Kit (QIAGEN, Hilden, Germany), according to the RNA cleanup protocol. The RNA integrity was examined using an Agilent RNA 6000 Nano Kit on a 2100 Bioanalyzer (Agilent Technologies, Santa Clara, California, USA), and the RNA concentration was determined using a NanoDrop ND-2000 Spectrophotometer (ThermoFisher Scientific). Five to six micrograms of RNA extracted from each leaf sample was sent to Hokkaido System Science Co., Ltd., where a cDNA library was prepared using a NovaSeq 6000 S4 Reagent Kit. Then, 151-base paired-end transcriptome sequencing of each sample was conducted using an Illumina NovaSeq 6000 sequencer. RNA samples were extracted from *Q. glauca* and *L. edulis* using the same protocol.

### Transcriptome assembly and gene expression analysis

The Illumina RNA-seq reads of *Q. serrata* were preprocessed for quality control using FastP v0.23.2 (https://github.com/OpenGene/fastp). The cleaned reads were then assembled using Trinity v2.15.1 (https://github.com/trinityrnaseq/trinityrnaseq). Open reading frames (ORFs) of at least 50 bp were predicted using TransDecoder v5.5.0, and the longest ORFs among the isoforms were selected using the “aggregate” function of CDSKIT v0.10.10. Transcript abundance (transcripts per million, TPM) was estimated using the “quant” function of AmalgaKit v0.10.28 (https://github.com/kfuku52/amalgkit), which utilizes Kallisto v0.48.0 (https://github.com/pachterlab/kallisto) internally. All contigs were clustered using a hierarchical algorithm in MultiExperiment Viewer (MeV, version 4.9.0) (J. Craig Venter Institute), based on log₂ values of transcript abundance. The minimum value among all expression values was assigned to datapoints with no detectable expression.

### RNA extraction for cDNA cloning

We isolated the cDNAs of *QsIspS1, QgIspS1-like* and *LeIspS1-like* from the leaves of the respective tree species. For cDNA cloning, total RNA was extracted using a cetyltrimethylammonium bromide (CTAB) procedure, independently from the above protocol. *Q. serrata* leaf tissue (2-4 pieces, each 5 mm) was immersed in 500 µL of extraction buffer (3 % CTAB, 100 mM Tris-HCl pH 8.0, 1.4 M NaCl, and 20 mM EDTA) and homogenized with MN Bead Tubes Type G (Takara Bio Inc., Kusatsu, Japan) for 1 min. The mixture was then incubated at 65 °C for 60 min. After adding 500 µL of chloroform, the mixture was shaken and centrifuged to separate the organic phase. The total RNA in the aqueous phase was then precipitated with 330 µL of isopropanol and collected by centrifugation. The resulting pellet was washed with 500 µL of 75 % ethanol, air-dried, and resuspended in 50 µL of RNase-free water. Purification was performed using an RNase-free DNase set and a RNeasy Plant Mini Kit (Qiagen). First-strand cDNA was synthesized using the SuperScript IV First-Strand Synthesis System (ThermoFisher Scientific). For *Q. glauca* and *L. edulis*, cDNA was synthesized from 500 ng of total RNA per sample using a High-Capacity RNA-to-cDNA Kit (Applied Biosystems, Foster City, CA), following the manufacturer’s instructions.

### Cloning of QsIspS1, QgIspS1-like and LeIspS1-like genes

The cDNA sequences containing the complete open reading frames (ORFs) of *QsIspS1* (1,827 bp), *QgIspS1-like* (1,827 bp), and *LeIspS1-like* (1,827 bp) were amplified by polymerase chain reaction (PCR) using KOD One® (Toyobo, Osaka, Japan) as a template. The primer pairs used for each cDNA were as follows: QsIspS1_5’UTR_Fw and QsIspS1_3’UTR_Rv for *QsIspS1*; QgIspS1-like_5’UTR_Fw and QgIspS1-like_3’UTR_Rv for *QgIspS1-like*; and LeIspS1-like_5’UTR_Fw and LeIspS1-like_3’UTR_Rv for *LeIspS1-like* (see Supplementary Table S1). *QsIspS1* cDNA amplification was performed with an initial denaturation step at 94 °C for 1 min, followed by 40 cycles of denaturation at 98 °C for 10 sec, annealing at 50 °C for 5 sec, and extension at 68 °C for 10 sec, concluding with a final extension step at 68 °C for 5 min. The amplifications of *QgIspS1-like* and *LeIspS1-like* cDNAs were performed with an initial denaturation at 94 °C for 1 min, followed by 40 cycles of denaturation at 98 °C for 10 sec, annealing at 51 °C for 5 sec, and extension at 68 °C for 10 sec, followed by a final extension at 68 °C for 5 min. These amplicons were inserted independently into the pGEM T-easy vector (Promega, Madison, WI) by TA cloning to confirm the sequence.

### Homology analysis of QsIspS1

The IspS sequences were collected from GenBank (https://www.ncbi.nlm.nih.gov/), the Sol Genomics Network (https://solgenomics.net), and UniProt (https://www.uniprot.org/) (see Supplementary Table S3). The amino acid identity of these sequences with QsIspS1 was calculated using Local BLAST (see Supplementary Table S2). The sequences were also aligned using Clustal Omega (Madeira et al., 2022; https://www.ebi.ac.uk/jdispatcher/msa/clustalo). TargetP (Nielsen et al., 1997; Emanuelsson et al., 2000) and DeepLoc (Ødum et al., 2024) were used to predict the TP.

### Phylogenetic analysis

TBLASTX (v2.14.0) searches were conducted to identify TPS homologs (E-value ≤ 0.01 and >50 % query coverage) against the coding sequences of 24 species (see Supplementary Table S4). A set of 63 pre-selected TPS sequences from *Arabidopsis thaliana* and *Solanum lycopersicum* were used as queries. The retrieved coding sequences were supplemented with 26 known IspS sequences from 26 species, and 50 known monoterpene synthase sequences from 29 species, as well as an additional *Q. serrata* 26 *TPS* genes, myrcene synthase from *Q. ilex*, and *QgIspS1-like* from *Q. glauca* (see Supplementary Table S5). Near-duplicate sequences were removed by clustering with MMseqs2 easy-cluster (v16.747c6, https://github.com/soedinglab/MMseqs2) at a minimum sequence identity of 0.99, with one representative sequence retained per cluster. All sequences were converted to in-frame codons using the “pad” function of CDSKIT, and stop and ambiguous codons were masked as gaps using the “mask” function. The amino acid sequences translated from these coding sequences were aligned using MAFFT v7.520 with the —auto option, trimmed using ClipKIT v2.1.1, and back-translated using the “backtrim” function in CDSKIT. A maximum-likelihood gene tree was reconstructed using IQ-TREE v2.2.5 (https://github.com/iqtree/iqtree2) with the GTR+G4 substitution model. Branch support was assessed via ultrafast bootstrapping with 1,000 replicates (-ufboot 1000) and optimization by hill-climbing nearest-neighbor interchange (-bnni). The root position was inferred using the non-reversible amino acid substitution model for plants (-m NQ.plant).

### Construction of plasmids for expression in plant host

The full-length CDS of *QsIspS1* was amplified by PCR using KOD One® with the primer pair QsIspS1_Nb_Fw and QsIspS1_Nb_Rv (see Supplementary Table S1) and pGEM-*QsIspS1* as the template. The amplification process included an initial denaturation step at 94 °C for 1 min, followed by 40 cycles of denaturation at 98 °C for 10 sec, annealing at 50 °C for 5 sec, and extension at 68 °C for 10 sec. A final extension step was performed at 68 °C for 5 min. The PCR product was inserted into the SalI site of the pTKB3 vector (Nosaki et al., 2021) via an in-fusion reaction, creating the pTKB3-*QsIspS1* construct.

To analyze subcellular localization, the 5’ terminal region containing the predicted translation initiation site (TP) of *QsIspS1*, *QsIspS1TP* (120 bp), was amplified using primers QsIspS1TP_Fw and QsIspS1_CDS_Rv, as well as QsIspS1TP_Rv (see Supplementary Table S1). The amplification process included an initial denaturation step at 94 °C for 1 min, followed by 40 cycles of denaturation at 98 °C for 10 sec, annealing at 50 °C for 5 sec, and extension at 68 °C for 10 sec for *QsIspS1* and 5 sec for *QsIspS1TP*. A final extension step was performed at 68 °C for 5 min. The PCR product was inserted into the pENTR™/D-TOPO® vector (ThermoFisher Scientific) via a directional TOPO reaction. It was then introduced into the binary vector pGWB505 (Nakagawa et al., 2007) via LR recombination, resulting in a plasmid containing *35Spro-QsIspS1TP-sGFP-Tnos*.

### Transient expression of recombinant protein in *N. benthamiana* for enzymatic characterization

pTKB3-*QsIspS1* and pTKB3-*EGFP* (Nozaki et al., 2021) were introduced into *Agrobacterium tumefaciens* GV3101 individually. The resulting transformants were pre-cultured in 3 mL of Luria-Bertani (LB) liquid medium containing 25 μg mL^-1^ rifampicin, 28 μg mL^-1^ gentamicin, and 50 μg mL^-1^ kanamycin for two days. Then, 20 μL of the pre-cultures were inoculated into 20 mL of LB liquid medium containing 25 μg mL^-1^ rifampicin, 28 μg mL^-1^ gentamicin, and 50 μg mL^-1^ kanamycin. The cultures were incubated until the OD_600_ reached 0.8-1.2. After washing the bacterial pellet with deionized water containing 100 μM acetosyringone, the *Agrobacterium* cells were resuspended in deionized water containing 100 μM acetosyringone and adjusted to an OD_600_ of 0.2. The cell suspensions were infiltrated into six-week-old *N. benthamiana* leaves using a 1-ml syringe (Terumo, Tokyo, Japan). Three days post-agroinfiltration, the treated leaves were collected and mashed on ice with 2.5 mL extraction buffer (100 mM KPi buffer at pH 7.0 containing 10 mM dithiothreitol (DTT), one tablet of Complete Mini EDTA-free (Sigma-Aldrich), and 0.1 g polyvinylpolypyrrolidone). The extract was then centrifuged at 9,200 × g for 20 minutes at 4 °C to remove debris. The supernatant was then passed through a PD-10 column (GE Healthcare, Illinois, USA), which had been equilibrated with reaction buffer (50 mM Tris-HCl, pH 7.5, containing 1 mM DTT).

### Site-directed mutagenesis of *QsIspS1* and *QgIspS1-like*

The full-length CDSs of *QsIspS1* and *QgIspS1-like* were amplified by PCR using KOD One®, with the primer pairs QsIspS1_F326I_Fw and QsIspS1_F326I_Rv for *QsIspS1*, whereas QgIspS1-like_I326F_Fw and QgIspS1-like_I326F_Rv for *QgIspS1-like* (Supplementary Table S1), in which pGEM-*QsIspS1* and pGEM-*QgIspS1-like* as templates, respectively. Amplifications of *QsIspS1 (F326I)* and *QgIspS1-like (I326F)* were carried out with an initial denaturation at 94 ℃ for 1 min followed by 40 cycles of denaturation at 98 ℃ for 10 sec, annealing at 51 ℃ for 25 sec, and extension at 68 ℃ for 10 sec. A final extension was performed at 68 °C for 5 min. The amplicon was inserted into the pGEM T-easy vector (Promega, Madison, WI) via TA cloning to confirm the sequence.

### Construction of plasmids for *E. coli* expression

The *QsIspS1, QgIspS1-like, QsIspS1(F326I)* and *QgIspS1-like(I326F)* lacking their 5’-terminal 93 bp were amplified by PCR using KOD One® with primer pairs ΔTP-QsIspS1_Fw and ΔTP-QsIspS1_Rv for *QsIspS1* and *QsIspS1(F326I)*, ΔTP-QgIspS1-like_Fw and ΔTP-QgIspS1-like_Rv for *QgIspS1-like* and *QgIspS1-like(I326F)* (Supplementary Table S1). As templates of the PCR, pGEM-*QsIspS1*, pGEM-*QgIspS1-like,* pGEM-*QsIspS1(F326I)*, pGEM-*QgIspS1-like(I326F)* were used. Amplification was carried out with an initial denaturation at 94 ℃ for 1 min followed by 40 cycles of denaturation at 98 ℃ for 10 sec, annealing at 51 ℃ for 5 sec, and extension at 68 ℃ for 10 sec. A final extension was performed at 68 °C for 5 min. The *EGFP* was also amplified by PCR using PrimeSTAR Max and the primer pairs EGFP_Fw and EGFP_Rv (Supplementary Table S1). pTKB3-*EGFP* was used as a template for the PCR. The amplification process involved an initial denaturation step at 96 ℃ for 2 min followed by 38 cycles of denaturation at 98 ℃ for 30 sec, annealing at 55 ℃ for 5 sec, and extension at 72 ℃ for 15 sec. A final extension was then performed at 72 °C for 5 min. The PCR product was then inserted into the pET22b(+) vector (Novagen, Madison, WI) via an “In-Fusion” reaction (Takara Bio) using the NdeI and BamHI sites.

### Enzymatic characterization with recombinant proteins expressed in *E. coli*

pET22b-*ΔTP*-*QsIspS1-6×His*, *ΔTP*-*QgIspS1-like-6×His, ΔTP*-*QsIspS1(F326I)-6×His*, and *ΔTP*-*QgIspS1-like(I326F)-6×His* were individually introduced into *E. coli* BL21 (DE3) for recombinant protein production. Transformants were cultured in 125 mL LB liquid medium containing 80 μg mL^-1^ ampicillin with shaking at 180 rpm at 37 °C. At a range of OD_600_ 0.5-0.8, recombinant protein production was induced by the addition of 250 µL 1 M isopropyl-β-D-thiogalactopyranoside and cultured further for 3 h at 34 °C. Cells were collected by centrifugation (9,200 × *g*, 10 min, 4 °C), and the bacterial pellet was resuspended in 650 μL lysis buffer (50 mM NaPi buffer, 10 mM imidazole, 300 mM NaCl), which was sonicated by Branson Sonifier 250 (duty cycle 40 %, output 3) (Emerson Electric Co., Missouri, US) followed by centrifugation (13,000 × *g*, 5 min, 4 °C) to remove the cell debris. The supernatant was subjected to affinity purification via His-tag using Ni-NTA Agarose (QIAGEN). The purified protein was used for enzyme assays. The purity of these enzymes was checked by SDS-PAGE (Supplementary Fig. S7). The concentration of the protein is quantified using a Qubit® 2.0 Fluorometer and the Qubit Protein Assay Kit, both from Thermo Fisher Scientific.

### In vitro assays

To detect QsIspS1 enzyme activity, in vitro assays were performed using the crude enzyme extracted from *N. benthamiana* leaves and the purified ΔTP-QsIspS1-6×His from *E. coli*. The standard reaction mixture (200 µL) contains 0.5 mM DMAPP, 20 mM MgCl₂, and 160 µL of enzymes. The reaction mixtures were then incubated in 2-mL glass vials (Thermo Fisher Scientific). The negative control assays shown in Fig. 4 contain an incubation mixture with 6.6 μg of ΔTP-QsIspS1-6×His, except for the buffer line. The following substitutions were made: milli-Q water was used instead of DMAPP; reaction buffer was used instead of enzymes; the enzymes were heat-denatured at 95 °C for 20 min and used in the enzyme (heat) line; and excessive EDTA was added instead of MgCl₂. In the negative control assays shown in Supplementary Figs. S8 and S12, the incubation mixtures contain 4.5-6.8 μg of enzymes for DMAPP assays, and 1.6-1.7 μg of enzymes for GPP assays, except for the buffer line. The standard reaction mixture for substrate specificity assays comprises 20 mM MgCl₂, 0.5 mM DMAPP, GPP, or FPP, and 160 μL of purified ΔTP-QsIspS1-6×His, ΔTP-QgIspS1-like-6×His, ΔTP-QsIspS1(F326I)-6×His, or ΔTP-QgIspS1-like(I326F)-6×His. For the kinetic analysis of ΔTP-QsIspS1-6×His and ΔTP-QgIspS1-like-6×His, the standard reaction mixture contains 20 mM MgCl₂, 0, 1, 2, 4, 6, or 10 mM DMAPP, or 0, 25, 50, 100, 200, 400, or 600 μM GPP and 160 μL of the purified protein to a final volume of 200 μL. For mutagenesis assays, the standard reaction mixture contained 20 mM MgCl₂, 0.5 mM DMAPP or GPP, and 160 µL of purified ΔTP-QsIspS1-6×His, ΔTP-QgIspS1-like-6×His, ΔTP-QsIspS1(F326I)-6×His, or ΔTP-QgIspS1-like(I326F)-6×His. All incubation steps were performed at 37 °C for 30 min.

### GC/MS analysis

Volatile compounds in the headspace of an enzyme incubation mixture were absorbed with an SPME fiber, (df 85 μm, CAR/PDMS, needle size 24-gauge, StableFlex; Sigma-Aldrich). The VOCs absorbed by the SPME fiber were analyzed using a Shimadzu GC-2030 equipped with a photoionization detector. Separation was performed with a DB-624 column (30 m, 0.25 mm ID, 1.4 μm df; Agilent Technologies, Santa Clara, CA). The temperature gradient program was conducted from 35 ℃ to 200 ℃ over 16 min, with an initial 35 ℃ for 5 min, and helium was used as a carrier gas with a flow rate of 1.22 mL min^-1^. For monoterpene, the gradient program was conducted from 50 ℃ to 200 ℃ over 15 min, with the flow rate at 2.53 mL min^-1^. For sesquiterpenes, the gradient program was conducted from 100 ℃ to 250 ℃ with the flow rate at 2.38 mL min^-1^. However, no peaks were observed in the retention time region where the sesquiterpenes were detected. All analyses were performed in splitless mode. The enzymatic reaction products were quantified using the same headspace analysis with standard specimens of isoprene, β-ocimene, β-myrcene, (1R)-(+)-α-pinene, and (-)-β-pinene dissolved in aqueous methanol. The peak area of non-enzymatic products (detected in buffer-treated lines) was subtracted from that of each sample before the quantification by standards. All peaks detected in the assays were verified as falling within the range of the calibration curve (Supplementary Fig. S17). To detect both isoprene and monoterpenes in the VOC analysis, the temperature gradient program was conducted from 35 ℃ to 200 ℃ over 21.5 min. The initial temperature was 35 °C for five minutes. Helium was used as the carrier gas at a flow rate of 1.22 mL min^-1^. The annotation of the *trans*-or *cis*-isomer of β-ocimene, as well as the annotations for α-phellandrene and (+)-sabinene, were based on the GC-MS library.

### Subcellular localization of QsIspS1

pGWB505-*QsIspS1*, pGWB505-*QsIspS1TP*, and pHKN29 (Kumagai et al., 2003), which contains *P35S-sGFP-Tnos* (control), were introduced individually into *A. tumefaciens* LBA4404. The competent LBA4404 cells and bacterial transformants were cultured in a selection medium containing 50 μg mL^-1^ streptomycin and 100 μg mL^-1^spectinomycin. The transformants were then infiltrated into four-week-old *N. benthamiana* leaves using the agroinfiltration method. Two days after agroinfiltration, leaf epidermal cells were observed using a FV3000 confocal laser scanning microscope (Olympus, Tokyo, Japan) with a UPlanSApo objective lens (Olympus). The excitation wavelengths were 488 nm for GFP fluorescence using an OBIS 488-20 LS CW laser (Coherent, Santa Clara, CA, USA) and 664 nm for chlorophyll autofluorescence using an OBIS LX 640-nm, 40-mW laser (Coherent). The collection bandwidth and gains were set to 500–540 nm for GFP fluorescence and 650–750 nm for chlorophyll autofluorescence. The laser intensities for GFP and chlorophyll autofluorescence were 3.6 % and 1.0 %, respectively. Images were captured using an Olympus FV31S-SW and analyzed using an Olympus FV31S-DT. For each treatment, more than three cells were observed per leaf, and three leaves were examined per individual. Observations were performed on two biological replicates.

### qRT-PCR

Six individuals of *Q. serrata* and *Q. glauca* were sampled on three sunny days in August (August 5^th^, 6^th^, and 8^th^, 2025). The sampling times were in the morning (7:45-9:30 AM), the afternoon (1:30-3:00 PM), and the evening (6:30-8:30 PM). Sampling temperature and humidity were measured using a digital thermo-hygrometer CR-1100B (CRECER Co., Ltd., Tokyo, Japan), and light intensity was measured using a digital separate-type illuminance meter Model 78747 (Shinwa Rules Co., Ltd., Niigata, Japan). These measurements are shown in Supplementary Tables S6-S9. Two individuals of *Q. serrata* and *Q. glauca* were collected each day. Total RNA was extracted from the leaves of *Q. serrata* and *Q. glauca* at different times of day Then, cDNA was synthesized using a standard protocol with ReverTra Ace qPCR RT Master Mix with gDNA Remover (Toyobo). RT-PCR experiments were performed using a CFX96 Deep Well real-time system (Bio-Rad) with a Thunderbird SYBR qPCR Mix (Toyobo) and primer pairs (QsTPS15_qPCR_Fw/Rv, QgIspS1-like_qPCR_Fw/Rv, and Actin_Fw/Rv as shown in Supplementary Table S1) with the synthesized cDNA serving as the template. The *Quercus*-conserved actin gene was used as a reference gene (Marum et al., 2012). The amplification protocol consisted of an initial denaturation step at 95 °C for 1 min, followed by 40 cycles of denaturation at 95 °C for 15 sec and annealing and extension at 62 °C for 60 sec for *QsTPS15* and 35 cycles for *QgIspS1-like*. Primer specificity was confirmed by melting curve analysis, which involved additional heating at additional heating at 65 °C and 95 °C for 5 sec each. Relative expression levels were analyzed using the corresponding standard curve and normalized to the reference gene.

### Quantification of VOC emitted from leaves of *Q. serrata* and *Q. glauca*

The VOC trapping apparatus was constructed using a plastic Petri dish (90 mm diameter x 19 mm height, Koryo Chemical Industry, Nara, Japan) to which two holes were drilled on the side. One hole was fitted with a silicone septum for SPME, and the other served as a leaf petiole (Supplementary Fig. S18). The dish was then used to sandwich an expanded leaf of *Q. serrata* and *Q. glauca* that had been grown in a pot, and the gap was sealed. The headspace of the transformed *Q. serrata* and *Q. glauca* leaves was captured using an SPME fiber. The fiber was inserted into the VOC trapping apparatus for 15 min, then subjected to GC-MS analysis. As a negative control, VOCs were trapped and analyzed from an empty Petri dish. Meanwhile, the leaf area was determined using ImageJ (Schneider et al., 2012). Quantification was performed by normalizing the peak areas of *m/z* = 67 for isoprene and *m/z* = 93 for monoterpenes to the leaf area.

## Acknowledgements

We thank Dr. Thomas D. Sharkey for providing an *E. coli* expression plasmid containing *E. globulus IspS*, which was used in a preliminary test as a positive control. We also thank Dr. Tsuyoshi Nakagawa for pGWB505 vector, Dr. Hiroshi Kouchi for the plasmid pHKN29, Dr. Hirobumi Yamamoto for providing DMAPP, Dr. Yuki Tobimatsu and Dr. Senri Yamamoto for technical support for GC-MS analysis. We thank Ms. Kaori Kanazawa RISH of Kyoto University for technical assistance. We also thank the Development and Assessment of Sustainable Humanosphere (DASH) system of the Research Institute for Sustainable Humanosphere (RISH), Kyoto University for a plant growth room.

## Author Contributions

S.K., R.M., A.J.N., and K.Y. conceived and designed the research. R.M. A. Sugiyama and K.Y. supervised experiments. Y.I. and A. Satake obtained plant samples and conducted RNA-seq experiment. K.F., N.S., and S.K. analyzed RNA-seq data. K.M. provided Tsukuba system vectors. S.K., T.S., and Y.K. performed enzymatic characterization. S.K. performed subcellular localization analysis. S.K., R.M., and K.Y. wrote the manuscript with contributions from all the authors.

## Supplementary data

**Supplementary Table S1.** List of primers used in this study.

**Supplementary Table S2.** Amino acid identities among the gene products of QsTPS15 and those reported as IspSs. The accession numbers of the putative IspSs are listed in Supplementary Table S3. The alphabets represent the TPS subfamily to which each gene belongs. * Reported enzyme activity, † described as IspS in reports, and ‡ predicted as IspS in databases.

**Supplementary Table S3.** Accession numbers of reported IspSs and *Quercus* TPS (myrcene synthase of *Q. ilex*). The IspS sequences were collected from Genbank (http://www.ncbi.nlm.nih.gov/genbank/), and UniProt (https://www.uniprot.org/).

**Supplementary Table S4.** File names, sources, and species of *TPS* sequences used to construct the phylogenetic tree in Fig. 2 and Supplementary Fig. S6.

**Supplementary Table S5.** Sequence labels and species names of IspS used to construct a phylogenetic tree in Fig. 2 and Supplementary Fig. S6.

**Supplementary Table S6.** The temperature at each sampling time for *Q. serrata* and *Q. glauca*.

**Supplementary Table S7.** The humidity at each sampling time for *Q. serrata* and *Q. glauca*.

**Supplementary Table S8.** The illuminance at each sampling time for *Q. serrata* and *Q. glauca*. Illuminance was measured under sunlight. The lux scores were converted to µmol photons m^-2^s^-1^ using the conversion coefficients reported by Thimijan and Heins (1983).

**Supplementary Table S9.** The times of sunrise and sunset on each sampling day.

**Supplementary Fig. S1.** Leaves and buds of *Q. serrata* Leaves (a) and buds (b) of *Q. serrata were* sampled at Kyushu University. The buds are indicated by blue circles.

**Supplementary Fig. S2.** Seasonal changes in the expression patterns of *Q. serrata TPS*s in leaf buds. (a) Heatmap showing contigs belonging to the TPS family based on the seasonal transcriptome dataset of *Q. serrata* leaf buds (n = 1). The expression of *QsTPS26* was undetectable at all data points. N.D., not detected. (b) TPM-based expression levels of *QsTPS*s (n = 1). The expression profile of each contig is shown individually in Supplementary Fig. S2b.

**Supplementary Fig. S3.** Seasonal expressions of each *QsTPS* in leaves and leaf buds. The Seasonal expression data of each *QsTPS* in leaves (a) and buds (b) were individually shown (n = 1). N.D., not detected.

**Supplementary Fig. S4.** Diurnal isoprene emission and *QsIspS1* expression in *Q. serrata* leaves. GC-MS chromatograms of isoprene (*m/z* = 67) and monoterpenes (*m/z* = 93) emitted from *Q. serrata* leaves are shown in (a) and (b), respectively. (c) and (d) show the amount of isoprene emission and *QsIspS1* expression in *Q. serrata l*eaves at morning (7:45-9:30 AM), noon (1:30-3:00 PM), and evening (6:30-8:30 PM). Bars indicate the mean of biological replicates, and dots represent the values of each replicate. An outlier was excluded from the morning sample by Dixon’s Q test before statistical analysis, and the same individual was excluded from the VOC emission analysis. Games-Howell’s test, *p\** < 0.05, n = 5 (morning) and 6 (afternoon and evening). Different alphabets represent significance. Specific information on climate and diurnal conditions during the sampling days is provided in Supplementary Tables S6-9.

**Supplementary Fig. S5.** Sequence alignment of the gene product of *QsTPS15* alongside reported IspS polypeptide sequences. The IspS sequences were aligned using Clustal Omega. The predicted transit peptide sequences, the D-rich Mg^2+^-binding motifs (DDxxD and NSE/DTE), the RR(x)8W motif, and the four amino acids representing the “isoprene score” are highlighted in yellow, green, purple, and orange, respectively. The transit peptide sequences were predicted using the TargetP2.0 and DeepLoc2.1 programs.

**Supplementary Fig. S6.** Phylogenetic tree of plant TPSs, including QsTPSs The phylogenetic tree was constructed using QsTPSs, reported IspSs, and TPSs obtained from other plant taxa. The TPS-a, -b, -c, and -g subfamilies are colored blue, red, yellow, and green, respectively. The branches of the IspSs in subfamilies TPS-b, -c, and -d are highlighted in red, yellow, and purple, respectively. The branches of the 26 QsTPSs are colored light blue. The bar indicates 0.5 substitutions per site. CpIspS: *Calohypnum plumiforme* IspS, PsIspS: *Pinus sabiniana* IspS.

**Supplementary Fig. S7.** SDS-PAGE of recombinant proteins expressed in *E. coli* that are affinity-purified with Ni-agarose column. The following proteins are shown: EGFP, ΔTP-QsIspS1-6×His, ΔTP-QgIspS1-like-6×His, ΔTP-QsIspS1(F326I) -6×His and ΔTP-QgIspS1-like(I326F) -6×His.

**Supplementary Fig. S8.** Negative control assays of QsIspS1. The following proteins are shown:ΔTP-QsIspS1-6×His, EGFP-6×His, transiently expressed in *E. coli* (a) or QsIspS1, EGFP, transiently expressed in *N. benthamiana* (b). The reaction buffer was also assayed with DMAPP. Bars indicate the mean of biological replicates, and dots represent the values of each replicate. Dunnett’s test (*p\** < 0.05, n = 3). The asterisk indicates a statistically significant difference. N.S., not significant.

**Supplementary Fig. S9.** Kinetic analysis of recombinant QsIspS1 with DMAPP as its substrate. Kinetic analysis of QsIspS1 for DMAPP using ΔTP-QsIspS1-6×His as the enzyme. Reaction mixtures were incubated for 30 min with different concentrations of DMAPP (0–20 mM). The *K*_1/2_ of QsIspS1 was estimated to be 12 ± 9 mM, which was calculated using nonlinear fitting in SigmaPlot 14.5.

**Supplementary Fig. S10.** Sequence alignment of the gene product of *QsIspS1, QgIspS1-like,* and *LeIspS1-like*. Sequences were aligned using Clustal Omega. Predicted transit peptide sequences, D-rich Mg^2+^-binding motifs (DDxxD and NSE/DTE), RR(x)8W motif, and the four amino acids representing the “isoprene score” are highlighted in yellow, green, purple, and orange, respectively. The transit peptide sequences were predicted using the TargetP2.0 program.

**Supplementary Fig. S11.** Sequence alignment of the gene of *QsIspS1* and *LeIspS1-like*. Sequences were aligned using Clustal Omega. A nucleotide that induces a stop codon is highlighted in blue.

**Supplementary Fig. S12.** Negative control assays of QgIspS1-like. ΔTP-QgIspS1-like-6×His, EGFP-6×His, transiently expressed in *E. coli*, and the reaction buffer were assayed with GPP. Bar graphs show the amount of β-myrcene (a), *cis*-β-ocimene (b), and *trans*-β-ocimene (c). Bars indicate the mean of biological replicates, and dots represent the values of each replicate. Dunnett’s test (*p\** < 0.05, n = 3). The asterisk indicates a statistically significant difference. N.S., not significant.

**Supplementary Fig. S13.** Kinetic analysis of recombinant QgIspS1-like using GPP as its substrate. Microsomes were incubated for 30 min with various concentrations of GPP (0–600 μM) and recombinant ΔTP-QgIspS1-like-6×His was used as the enzyme. The *K*_m_ values for QgIspS1-like were estimated to be 220 ± 60 μM for β-myrcene (a), 190 ± 40 μM for *cis*-β-ocimene (b), and 180 ± 40 μM for *trans*-β-ocimene (c). These *K*_m_ values were calculated using nonlinear fitting in SigmaPlot 14.5.

**Supplementary Fig. S14.** Diurnal monoterpene emission and *QgIspS1-like* expression in *Q. glauca* leaves. Monoterpene emissions are shown in the GC-MS chromatogram (*m/z* = 93) and the MS spectrum of the major five peaks (a). Isoprene emissions are shown in the chromatogram (*m/z* = 67) (b). The amounts of total ocimene emissions (c) and *QgIspS1-like* expression (d) in *Q. glauca* leaves were measured at morning (7:45‒9:30 AM), afternoon (1:30‒3:00 PM), and evening (6:30‒8:30 PM) were measured. Bars indicate the mean of biological replicates, and dots represent the values of each replicate. Games-Howell’s test, *p\** < 0.05, n = 6 (morning, afternoon, and evening). The different alphabets represent significance. Specific information on climate and diurnal conditions during the sampling days is provided in Supplementary Tables S6-9.

**Supplementary Fig. S15.** Correlation of monoterpene emission and expression levels in the afternoon. Correlation of *QgIspS1-like* expression with the amounts of total ocimene (a), α-pinene (b), β-pinene (c), α-phellandrene (d), and (+)-sabinene (e) emissions.

**Supplementary Fig. S16.** Enzymatic characterization of mutant QsIspS1-like (F326I) and QgIspS1-like (I326F). Recombinant proteins of ΔTP-QsIspS1-like(F326I)-6×His (a) and ΔTP-QgIspS1-like(I326F)-6×His, which were expressed in *E. coli*, were used after affinity purification. The substrate preferences of QsIspS1-like(F326I) (a) and QgIspS1-like(I326F) (b) are shown with DMAPP, GPP, and FPP as substrates. N.D., not detected. Bars indicate the mean of biological replicates, and dots represent the values of each replicate. Statistical analysis was performed using the Tukey-Kramer test. Different letters indicate statistically significant differences (n = 3).

**Supplementary Fig. S17.** Standard curves of isoprene and monoterpenes analyzed in this study. Standard curves of isoprene (a), *trans*-β-ocimene (b), *cis*-β-ocimene (c), and myrcene (d).

**Supplementary Fig. S18.** VOC trapping from the leaves of *Quercus* trees. A leaf of either *Q. serrata* (the photo image) or *Q. glauca* was sandwiched with a Petri dish (90 mmφ x19 mm) through a hole that was sized for the leaf petiole. The SPME fiber was inserted from the other side of the hole. The VOCs in the headspace were trapped for 15 min.

## Data and Materials Availability

The accession numbers of QsIspS1, QgIspS1-like, and LeIspS1 are LC833859, LC870167, and LC870166. Transcriptome data generated in this work have been deposited in DDBJ and are available from the Sequence Read Archive under project PRJDB18102.

