## Supplemental Tables and Figures for "Molecular basis behind the isoprene emission diversity in Fagaceae"

Supplementary Table S1. List of primers used in this study.

| Primer name | Sequence (5'-3') |
|---|---|
| QsIspS1_5'UTR_Fw | TGGCATATACACCTGTCACAAA |
| QsIspS1_3'UTR_Rv | CACTAAGCATAGGCAGCTCAAC |
| QgIspS1-like_5'UTR_Fw | TGGCATATATACCTGTCACAAA |
| QgIspS1-like_3'UTR_Rv | AACTAAGCATAGGCAGCTCAAC |
| LeIspS1-like_5'UTR_Fw | TGGCATATATATCTGTCACAAA |
| LeIspS1-like_3'UTR_Rv | AACTAAGCAAAGGCAGCTCAAC |
| QsIspS1_Nb_Fw | CTATTTACAAGTCGAATGGCAGCACTCCAAGTC |
| QsIspS1_Nb_Rv | ATTCAGAATTGTCGACTAAAGGTGGATCTGGCTG |
| QsIspS1TP_Fw | CACCATGGCAGCACTCCAAGT |
| QsIspS1TP_Rv | TGTATTAGAAAGCACTTGTTT |
| QsIspS1_CDS_Rv | AAGGTGGATCTGGCTGTGGA |
| QsIspS1_F326I_Fw | CAAAAATGATTGCATTTATAACCGTCATTGATG |
| QsIspS1_F326I_Rv | ATGCAATCATTTTTGTTACCGCCTTACG |
| QgIspS1-like_I326F_Fw | CAAAAGTGTTTTCACTAATAACCATCATTGATGA |
| QgIspS1-like_I326F_Rv | GTGAAAACACTTTTGTTAGCTCCTTACGGCA |
| EGFP_Fw | AAGGAGATATACATAATGGTGAGCAAGGGCGAGG |
| EGFP_Rv | GCTCGAATTCGGATCGGACTTGTACAGCTCGTCCATGC |
| ΔTP-QsIspS1_Fw | AAGGAGATATACATAATGGCGAGCAAACAAGTGC |
| ΔTP-QsIspS1_Rv | GCTCGAATTCGGATCAAGGTGGATCTGGCTGTG |
| ΔTP-QgIspS1-like_Fw | AAGGAGATATACATAATGGCGAGCAAACAAGTTCTTTCC |
| ΔTP-QgIspS1-like_Rv | GCTCGAATTCGGATCAAGGTGGATATGGCCGTGG |
| QsTPS15_qPCR_Fw | ATGGGTTTGGCGTCTCTCAA |
| QsTPS15_qPCR_Rv | AGCCTCATGTAGACTCAGCA |
| QgIspS1-like_qPCR_Fw | GCCGTGAAAGATCTCCCTGA |
| QgIspS1-like_qPCR_Rv | AACGCCTTGCATAAATCCCC |
| Actin_qPCR_Fw | GCTGGTCGTGATCTCACTG |
| Actin_qPCR_Rv | CTTGGCAGTCTCAAGTTCCT |

Supplementary Table S2. Amino acid identities among the gene products of QsTPS15 and those reported as IspSs.

| Family | Species | Length (a.a.) | a.a. Identity (%) | TPS subfamily |
|---|---|---|---|---|
| Myrtaceae | *Melaleuca alternifolia* | 583 | 51 | b* |
| | *Eucalyptus globulus* | 582 | 52 | b* |
| | *Eugenia uniflora* | 581 | 54 | b‡ |
| | *Eucalyptus grandis* | 624 | 49 | b‡ |
| Salicaceae | *Populus alba* | 596 | 57 | b* |
| | *Populus tremuloides* | 595 | 57 | b† |
| | *Populus trichocarpa* | 558 | 58 | b† |
| | *Populus canescens* | 595 | 57 | b* |
| | *Populus balsamifera* | 593 | 57 | b* |
| | *Populus fremontii* | 595 | 58 | b* |
| | *Populus deltoides* | 599 | 56 | b* |
| | *Populus grandidentata* | 595 | 57 | b* |
| | *Populus nigra* | 595 | 57 | b* |
| | *Salix sp.* | 595 | 58 | b* |
| | *Populus euphratica* | 596 | 56 | b‡ |
| Leguminosae | *Pueraria montana var. lobata* | 608 | 55 | b* |
| | *Robinia pseudoacacia* | 534 | 55 | b* |
| | *Wisteria sp.* | 541 | 53 | b* |
| | *Glycine soja* | 606 | 53 | b‡ |
| | *Cajanus cajan* | 603 | 49 | b‡ |
| | *Mucuna pruriens* | 604 | 54 | b‡ |
| | *Arachis hypogaea* | 605 | 53 | b‡ |
| | *Glycine max* | 606 | 53 | b† |
| Hypericidae | *Humulus lupulus* | 613 | 45 | b* |
| Moraceae | *Ficus septica* | 592 | 61 | b* |
| | *Ficus virgata* | 591 | 63 | b* |
| | *Artocarpus heterophyllus* | 591 | 61 | b* |
| Poaceae | *Arundo donax* | 564 | 41 | b* |
| | *Arundo plinii* | 564 | 41 | b‡ |
| Arecaceae | *Bismarckia nobilis* | 585 | 50 | b† |
| | *Howea forsteriana* | 585 | 51 | b* |
| | *Phoenix canariensis* | 585 | 49 | b* |
| | *Sabal minor* | 585 | 49 | b* |
| | *Trachycarpus oreophilus* | 585 | 50 | b† |
| | *Washingtonia filifera* | 585 | 50 | b† |
| Casuarinaceae | *Casuarina equisetifolia* | 592 | 50 | b* |
| Anacardiaceae | *Mangifera indica* | 598 | 58 | b‡ |
| Convolvulaceae | *Ipomoea batatas* | 584 | 51 | b‡ |
| Pedaliaceae | *Sesamum indicum* | 588 | 49 | b‡ |
| Solanaceae | *Solanum lycopersicum* | 562 | 38 | b* |
| Hypnaceae | *Calohypnum plumiforme* | 651 | 28 | c* |
| Pinaceae | *Pinus sabiniana* | 614 | 36 | d-1* |

Accession numbers of putative IspSs are listed in Supplementary Table S3. Alphabets represent which TPS subfamily each gene belongs to. * Reported enzyme activity, and † described as IspS in reports, ‡ predicted as IspS in databases.

Supplementary Table S3. Accession numbers of reported IspSs and *Quercus* TPS (myrcene synthase of *Q. ilex*).

| Species | accession No. |
|---|---|
| *Melaleuca alternifolia* | AAP40638 |
| *Eucalyptus globulus* | BAF02831 |
| *Eugenia uniflora* | AIK19222 |
| *Eucalyptus grandis* | XP_039160987 |
| *Populus alba* | BAD98243 |
| *Populus tremuloides* | AAQ16588 |
| *Populus trichocarpa* | XP_006373002 |
| *Populus canescens* | Uniprot: Q50L36 |
| *Populus balsamifera* | AEK70964 |
| *Populus fremontii* | AEK70967 |
| *Populus deltoides* | AEK70966 |
| *Populus grandidentata* | AEK70965 |
| *Populus nigra* | CAL69918 |
| *Salix sp.* | AEK70969 |
| *Populus euphratica* | XP_011006402 |
| *Pueraria montana var. lobata* | AAQ84170 |
| *Robinia pseudoacacia* | AEK70968 |
| *Wisteria sp.* | AEK70970 |
| *Glycine soja* | XP_028221489 |
| *Cajanus cajan* | XP_020216726 |
| *Mucuna pruriens* | RDX57825 |
| *Arachis hypogaea* | XP_025679837 |
| *Glycine max* | XP_003533120 |
| *Humulus lupulus* | ACI32638 |
| *Ficus septica* | BAS30550 |
| *Ficus virgata* | BAS30551 |
| *Artocarpus heterophyllus* | UUF44715 |
| *Arundo donax* | ASF20076 |
| *Arundo plinii* | AXT98924 |
| *Bismarckia nobilis* | QUQ60718 |
| *Howea forsteriana* | QUQ60719 |
| *Phoenix canariensis* | QUQ60720 |
| *Sabal minor* | QUQ60721 |
| *Trachycarpus oreophilus* | QUQ60722 |
| *Washingtonia filifera* | QUQ60723 |
| *Casuarina equisetifolia* | BAS30549 |
| *Mangifera indica* | BDN86180 |
| *Ipomoea batatas* | GMC70530 |
| *Solanum lycopersicum* | tomato genome |
| *Calohypnum plumiforme* | BEH00593 |
| *Pinus sabiniana* | AEB53064 |
| *Quercus lobata* | XP_030931870 |
| *Quercus robur* | XP_050248080 |
| *Quercus ilex* | CAC41012 |

*IspS* sequences were collected in Genbank (http://www.ncbi.nlm.nih.gov/genbank/), Sol Genomics Network (tomato genome) (https://solgenomics.net), and Uniprot (https://www.uniprot.org/)

Supplementary Table S4. File names, sources, and species of *TPS* sequences used to construct the phylogenetic tree in Fig. 2 and Supplementary Fig. S6.

| Species | Source | File name |
|---|---|---|
| *Amborella trichopoda* | Phytozome | Amborella_trichopoda_Atrichopoda_291_v1.0.cds_primaryTranscriptOnly.fa.gz |
| *Ananas comosus* | Phytozome | Ananas_comosus_Acomosus_321_v3.cds_primaryTranscriptOnly.fa.gz |
| *Aquilegia coerulea* | Phytozome | Aquilegia_coerulea_Acoerulea_322_v3.1.cds_primaryTranscriptOnly.fa.gz |
| *Arabidopsis thaliana* | Phytozome | Arabidopsis_thaliana_Athaliana_447_Araport11.cds_primaryTranscriptOnly.fa.gz |
| *Betula platyphylla* | Phytozome | Betula_platyphylla_Bplatyphylla_679_v1.1.cds_primaryTranscriptOnly.fa.gz |
| *Carpinus fangiana* | NCBI | Carpinus_fangiana_GCA_006937295.1.fa.gz |
| *Carya illinoinensis* | Phytozome | Carya_illinoinensis_Cillinoinensis_573_v1.1.cds_primaryTranscriptOnly.fa.gz |
| *Castanea dentata* | Phytozome | Castanea_dentata_Cdentata_673_v1.1.cds_primaryTranscriptOnly.fa.gz |
| *Castanea mollissima* | NCBI | Castanea_mollissima_GCA_000763605.2.fa.gz |
| *Eucalyptus grandis* | Phytozome | Eucalyptus_grandis_Egrandis_297_v2.0.cds_primaryTranscriptOnly.fa.gz |
| *Juglans regia* | NCBI | Juglans_regia_GCF_001411555.2.fa.gz |
| *Lotus japonicus* | Phytozome | Lotus_japonicus_Ljaponicus_571_Lj1.0v1.cds_primaryTranscriptOnly.fa.gz |
| *Morella rubra* | NCBI | Morella_rubra_GCA_003952965.2.fa.gz |
| *Nymphaea colorata* | Phytozome | Nymphaea_colorata_Ncolorata_566_v1.2.cds_primaryTranscriptOnly.fa.gz |
| *Oryza sativa* | Phytozome | Oryza_sativa_Osativa_323_v7.0.cds_primaryTranscriptOnly.fa.gz |
| *Populus trichocarpa* | Phytozome | Populus_trichocarpa_Ptrichocarpa_533_v4.1.cds_primaryTranscriptOnly.fa.gz |
| *Quercus gilva* | NCBI | Quercus_gilva_GCA_023621385.1.fa.gz |
| *Quercus lobata* | NCBI | Quercus_lobata_GCF_001633185.2.fa.gz |
| *Quercus robur* | NCBI | Quercus_robur_GCF_932294415.1.fa.gz |
| *Quercus serrata* | NCBI | Quercus_serrata_longestCDS.fa.gz |
| *Quercus suber* | NCBI | Quercus_suber_GCA_002906115.4.fa.gz |
| *Quercus variabilis* | NCBI | Quercus_variabilis_CNA0051893.fa.gz |
| *Solanum lycopersicum* | Phytozome | Solanum_lycopersicum_Slycopersicum_691_ITAG4.0.cds_primaryTranscriptOnly.fa.gz |
| *Vitis vinifera* | Phytozome | Vitis_vinifera_Vvinifera_457_v2.1.cds_primaryTranscriptOnly.fa.gz |

Supplementary Table S5. Sequence labels and species names of *IspS*s used to construct a phylogenetic tree in Fig. 2 and Supplementary Fig. S6.

| Species | Sequence label | Species | Sequence label |
|---|---|---|---|
| *Artocarpus heterophyllus* | Artocarpus_heterophyllus_MZ493337.1_IspS | *Agastache rugosa* | Agastache_rugosa_AY055214.1 |
| *Arundo donax* | Arundo_donax_KX906604.1_IspS | *Artemisia annua* | Artemisia_annua_AF154124.1 |
| *Calohypnum plumiforme* | Calohypnum_plumiforme_LC765855.1_IspS | *Camellia sinensis* | Camellia_sinensis_MN135992.1 |
| *Casuarina equisetifolia* | Casuarina_equisetifolia_LC006089.1_IspS | *Cannabis sativa* | Cannabis_sativa_DQ839404.1 |
| *Eucalyptus globulus* | Eucalyptus_globulus_AB266390.1_IspS | *Cannabis sativa* | Cannabis_sativa_DQ839405.1 |
| *Ficus septica* | Ficus_septica_LC006090.1_IspS | *Citrus limon* | Citrus_limon_AF514289.1 |
| *Ficus virgata* | Ficus_virgata_LC006091.1_IspS_1 | *Citrus limon* | Citrus_limon_AF514288.1 |
| *Howea forsteriana* | Howea_forsteriana_MT512619.1_IspS | *Citrus limon* | Citrus_limon_AF514287.1 |
| *Humulus lupulus* | Humulus_lupulus_EU760349.1_IspS | *Citrus limon* | Citrus_limon_AF514286.1 |
| *Melaleuca alternifolia* | Melaleuca_alternifolia_AY279379.1_IspS | *Citrus unshiu* | Citrus_unshiu_AB110638.1 |
| *Phoenix canariensis* | Phoenix_canariensis_MT512620.1_IspS | *Citrus unshiu* | Citrus_unshiu_AB110636.1 |
| *Pinus sabiniana* | Pinus_sabiniana_JF719039.1_IspS | *Citrus unshiu* | Citrus_unshiu_AB110641.1 |
| *Populus alba* | Populus_alba_AB198180.1_IspS | *Citrus unshiu* | Citrus_unshiu_AB110642.1 |
| *Populus balsamifera* | Populus_balsamifera_JN173037.1_IspS | *Cucumis melo* | Cucumis melo |
| *Populus canescens* | Populus_canescens_AJ294819.1_IspS | *Lavandula angustifolia* | Lavandula_angustifolia_DQ263741.1 |
| *Populus deltoides* | Populus_deltoides_JN173039.1_IspS | *Lavandula angustifolia* | Lavandula_angustifolia_DQ263740.1 |
| *Populus fremontii* | Populus_fremontii_JN173040.1_IspS | *Litsea cubeba* | Litsea_cubeba_HQ651180.1 |
| *Populus grandidentata* | Populus_grandidentata_JN173038.1_IspS | *Litsea cubeba* | Litsea_cubeba_HQ651178.1 |
| *Populus nigra* | Populus_nigra_AM410988.1_IspS | *Magnolia grandiflora* | Magnolia_grandiflora_EU366430.1 |
| *Pueraria montana* | Pueraria_montana_var._lobata_AY316691.1_IspS | *Malus domestica* | Malus_domestica_JX848733.1 |
| *Quercus ilex* | Quercus_ilex_AJ304839.1_Myrcene_synthase | *Matricaria chamomilla var. recutita* | Matricaria_chamomilla_JQ255378.1 |
| *Quercus lobata* | Quercus_lobata_XM_031076010.1_IspS | *Medicago truncatula* | Medicago_truncatula_AY766248.1 |
| *Quercus robur* | Quercus_robur_XM_050392123.1_IspS | *Mentha citrata* | Mentha_citrata_AY083653.1 |
| *Robinia pseudoacacia* | Robinia_pseudoacacia_JN173041.1_IspS | *Mentha spicata* | Mentha_spicata_L13459.1 |
| *Sabal minor* | Sabal_minor_MT512621.1_IspS | *Nicotiana attenuata* | Nicotiana_attenuata_MN400708.1 |
| *Salix sp.* | Salix_sp._JN173043.1_IspS | *Ocimum basilicum* | Ocimum_basilicum_AY693647.1 |
| *Wisteria sp.* | Wisteria_sp._JN173042.1 | *Ocimum basilicum* | Ocimum_basilicum_AY693650.1 |
| *Quercus serrata* | Quercus_serrata_DN11064_c0_g1 | *Ocimum basilicum* | Ocimum_basilicum_AY693649.1 |
| *Quercus serrata* | Quercus_serrata_DN1140_c0_g1 | *Osmanthus fragrans* | Osmanthus_fragrans_KT591182.1 |
| *Quercus serrata* | Quercus_serrata_DN13219_c0_g1 | *Perilla frutescens* | Perilla_frutescens_AF271259.1 |
| *Quercus serrata* | Quercus_serrata_DN15502_c0_g1 | *Perilla frutescens* | Perilla_frutescens_AF241793.1 |
| *Quercus serrata* | Quercus_serrata_DN1562_c0_g1 | *Perilla frutescens var. acuta* | Perilla_frutescens_D49368.1 |
| *Quercus serrata* | Quercus_serrata_DN16154_c0_g1 | *peruviana cv. Sweet Laura* | Alstroemeria_peruviana_FR822739.1 |
| *Quercus serrata* | Quercus_serrata_DN17344_c1_g1 | *Phaseolus lunatus* | Phaseolus_lunatus_EU194553.1 |
| *Quercus serrata* | Quercus_serrata_DN20099_c0_g1 | *Salvia fruticosa* | Salvia_fruticosa_DQ785793.1 |
| *Quercus serrata* | Quercus_serrata_DN21870_c0_g1 | *Salvia officinalis* | Salvia_officinalis_AF051901.1 |
| *Quercus serrata* | Quercus_serrata_DN21977_c0_g1 | *Salvia officinalis* | Salvia_officinalis_AF051900.1 |
| *Quercus serrata* | Quercus_serrata_DN22894_c0_g1 | *Salvia officinalis* | Salvia_officinalis_AF051899.1 |
| *Quercus serrata* | Quercus_serrata_DN24395_c0_g1 | *Salvia pomifera* | Salvia_pomifera_DQ785794.1 |
| *Quercus serrata* | Quercus_serrata_DN25533_c0_g1 | *Salvia stenophylla* | Salvia_stenophylla_AF527416.1 |
| *Quercus serrata* | Quercus_serrata_DN25533_c1_g1 | *Santalum album* | Santalum_album_EU798692.1 |
| *Quercus serrata* | Quercus_serrata_DN28119_c0_g1 | *Schizonepeta tenuifolia* | Schizonepeta_tenuifolia_AF282875.2 |
| *Quercus serrata* | Quercus_serrata_DN28500_c0_g1 | *Vitis vinifera* | Vitis_vinifera_HM807386.1 |
| *Quercus serrata* | Quercus_serrata_DN37502_c0_g1 | *Vitis vinifera* | Vitis_vinifera_HM807385.1 |
| *Quercus serrata* | Quercus_serrata_DN37963_c0_g1 | *Vitis vinifera* | Vitis_vinifera_AY572987.1 |
| *Quercus serrata* | Quercus_serrata_DN40354_c0_g1 | *Vitis vinifera* | Vitis_vinifera_HM807387.1 |
| *Quercus serrata* | Quercus_serrata_DN4452_c0_g1 | *Vitis vinifera* | Vitis_vinifera_HM807390.1 |
| *Quercus serrata* | Quercus_serrata_DN4452_c0_g4 | *Vitis vinifera* | Vitis_vinifera_HM807383.1 |
| *Quercus serrata* | Quercus_serrata_DN47593_c0_g1 | *Vitis vinifera* | Vitis_vinifera_AY572986.1 |
| *Quercus serrata* | Quercus_serrata_DN5248_c0_g1 | *Quercus glauca* | Quercus_glauca_QgIspS1like |
| *Quercus serrata* | Quercus_serrata_DN6183_c1_g1 | | |
| *Quercus serrata* | Quercus_serrata_DN8111_c0_g1 | | |

Supplementary Table S6. The temperature at each sampling time for *Q. serrata* and *Q. glauca.*

| Individual | Temperature (℃) | | |
| --- | --- | --- | --- |
| | Morning | Afternoon | Evening |
| Serrata1 | 30.5 | 42.6 | 33.6 |
| Serrata2 | 31.0 | 43.7 | 32.3 |
| Serrata3 | 27.6 | 45.8 | 27.2 |
| Serrata4 | 28.0 | 42.0 | 26.8 |
| Serrata5 | 30.6 | 45.8 | 33.5 |
| Serrata6 | 30.6 | 45.8 | 33.5 |
| Glauca1 | 31.5 | 41.3 | 32.1 |
| Glauca2 | 32.4 | 47.7 | 31.6 |
| Glauca3 | 29.4 | 38.0 | 26.2 |
| Glauca4 | 30.1 | 39.1 | 26.1 |
| Glauca5 | 30.5 | 46.1 | 31.5 |
| Glauca6 | 30.5 | 46.1 | 31.5 |

Supplementary Table S7. The humidity at each sampling time for *Q. serrata* and *Q. glauca*.

| Individual | Humidity (%) | | |
|---|---|---|---|
| | Morning | Afternoon | Evening |
| Serrata1 | 75 | 37 | 54 |
| Serrata2 | 74 | 33 | 58 |
| Serrata3 | 82 | 28 | 75 |
| Serrata4 | 82 | 33 | 77 |
| Serrata5 | 73 | 30 | 55 |
| Serrata6 | 73 | 30 | 55 |
| Glauca1 | 73 | 33 | 64 |
| Glauca2 | 71 | 26 | 69 |
| Glauca3 | 79 | 38 | 78 |
| Glauca4 | 76 | 42 | 79 |
| Glauca5 | 73 | 29 | 68 |
| Glauca6 | 73 | 29 | 68 |

Supplementary Table S8. The illuminance at each sampling time for *Q. serrata* and *Q. glauca*. Illuminance was measured under sunlight. The lux scores were converted to µmol photons $m^{-2}s^{-1}$ using the conversion coefficients reported by Thimijan and Heins (1983).

| Individual | Illuminance (Lux/ µmol photons $m^{-2}s^{-1}$) | | |
|---|---|---|---|
| | Morning | Afternoon | Evening |
| Serrata1 | 2700/ 50.0 | 52600/ 974 | 83/ 1.54 |
| Serrata2 | 2360/ 43.7 | 69100/ 1280 | 0/ 0 |
| Serrata3 | 3800/ 70.4 | 54300/ 1010 | 0/ 0 |
| Serrata4 | 4010/ 74.3 | 45600/ 844 | 0/ 0 |
| Serrata5 | 3170/ 58.7 | 50100/ 928 | 150/ 2.78 |
| Serrata6 | 3170/ 58.7 | 50100/ 928 | 150/ 2.78 |
| Glauca1 | 3140/ 58.1 | 61900/ 1150 | 0/ 0 |
| Glauca2 | 5700/ 106 | 24500/ 454 | 0/ 0 |
| Glauca3 | 3580/ 66.3 | 4920/ 91.1 | 0/ 0 |
| Glauca4 | 4930/ 91.3 | 61100/ 1130 | 0/ 0 |
| Glauca5 | 2460/ 45.6 | 61700/ 1140 | 0/ 0 |
| Glauca6 | 2460/ 45.6 | 61700/ 1140 | 0/ 0 |

Supplementary Table S9. The times of sunrise and sunset on each sampling day.

| Date | Sunrise time | Sunset time |
|---|---|---|
| August 5th | 5:09 AM | 6:56 PM |
| August 6th | 5:10 AM | 6:55 PM |
| August 8th | 5:11 AM | 6:53 PM |

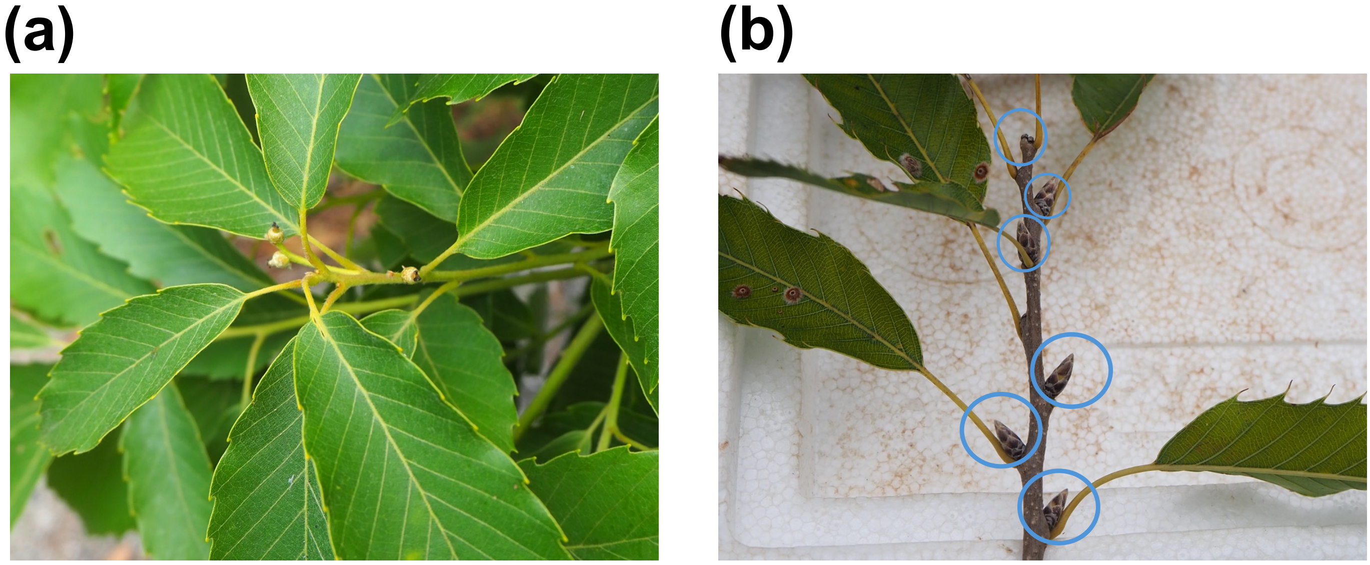


Supplementary Fig. S1. Leaves and buds of *Q. serrata*
Leaves (a) and buds (b) of *Q. serrata were* sampled at Kyushu University. The buds are indicated by blue circles.

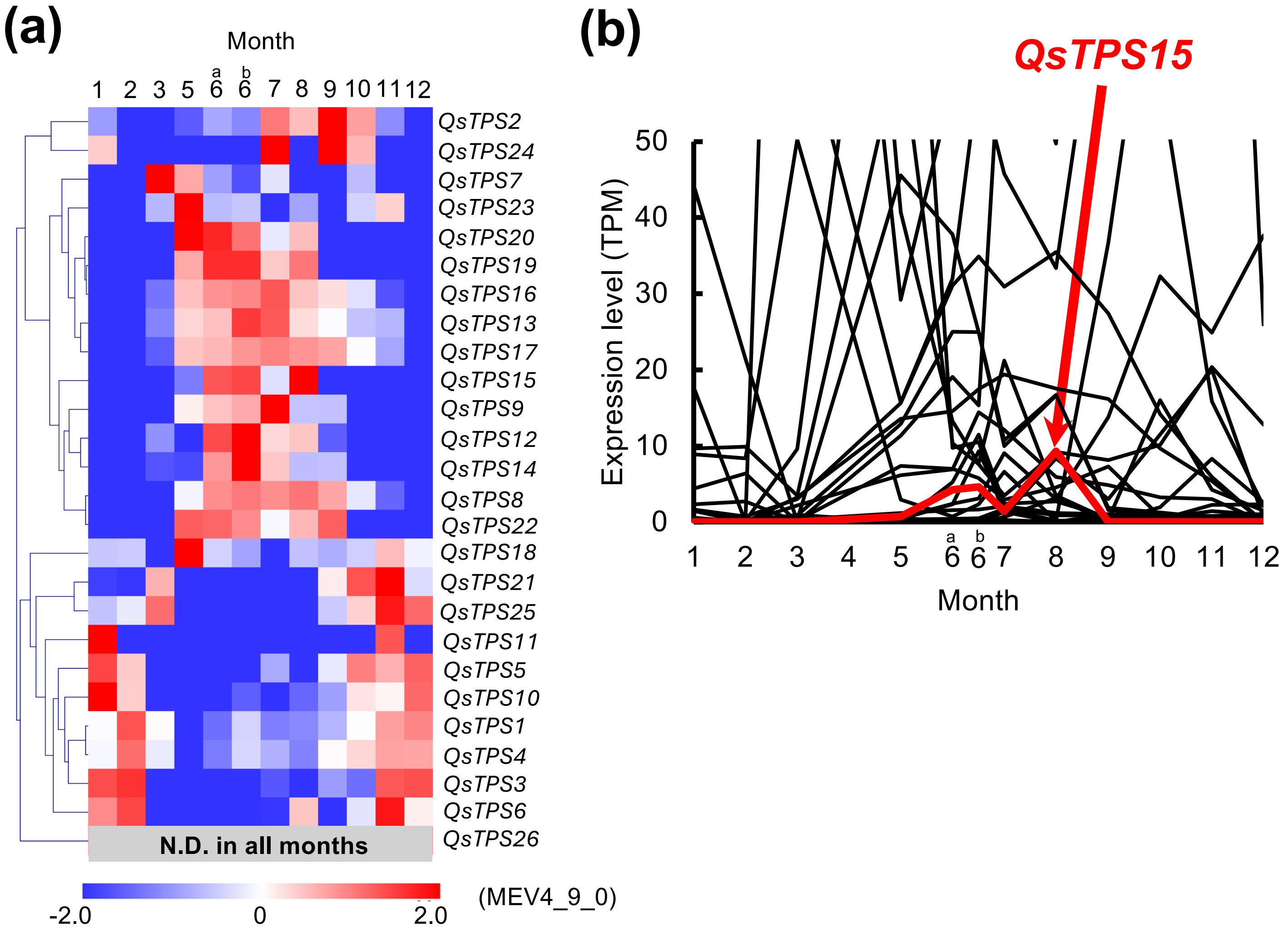


Supplementary Fig. S2. Seasonal changes in the expression patterns of *Q. serrata TPS*s in leaf buds. (a) Heatmap showing contigs belonging to the TPS family based on the seasonal transcriptome dataset of *Q. serrata* leaf buds (n = 1). The expression of *QsTPS26* was undetectable at all data points. N.D., not detected. (b) TPM-based expression levels of *QsTPS*s (n = 1). The expression profile of each contig is shown individually in Supplementary Fig. S2b.

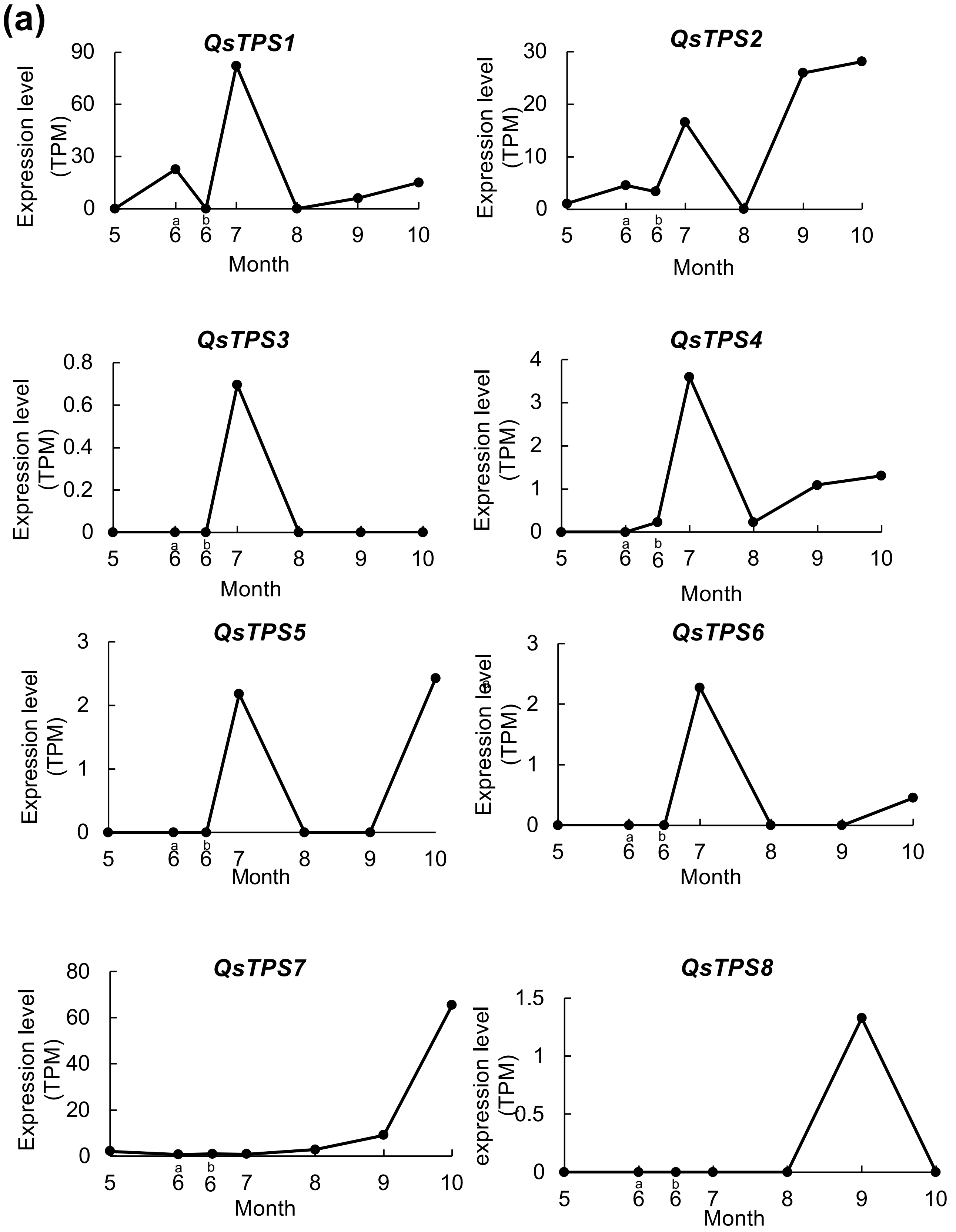


Supplementary Fig. S3. Seasonal expressions of each *QsTPS* in leaves and leaf buds. The Seasonal expression data of each *QsTPS* in leaves (a) and buds (b) were individually shown (n = 1). N.D., not detected.

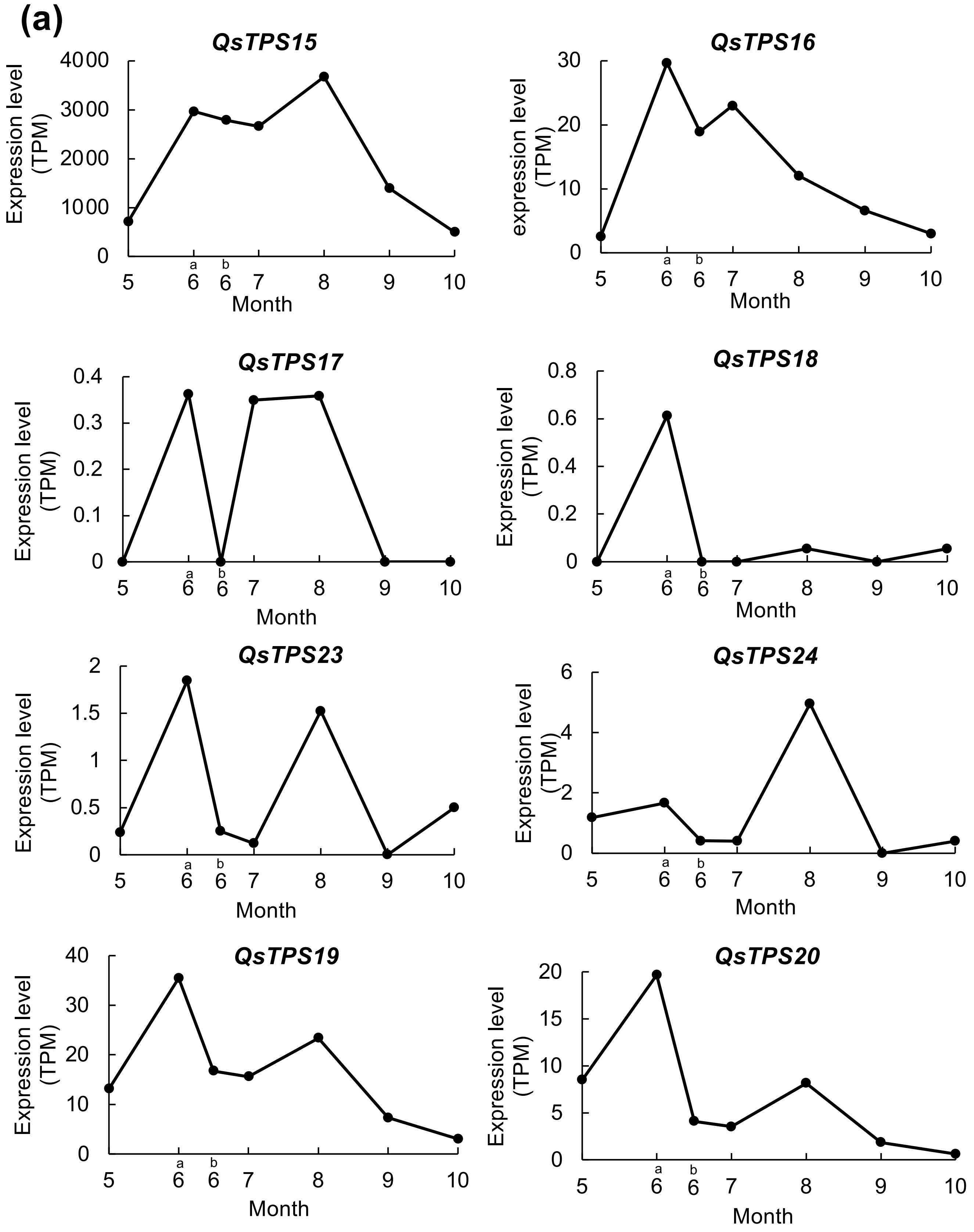


Supplementary Fig. S3. Seasonal expressions of each *QsTPS* in leaves and leaf buds. (*-continued*) The Seasonal expression data of each *QsTPS* in leaves (a) and buds (b) were individually shown (n = 1). N.D., not detected.

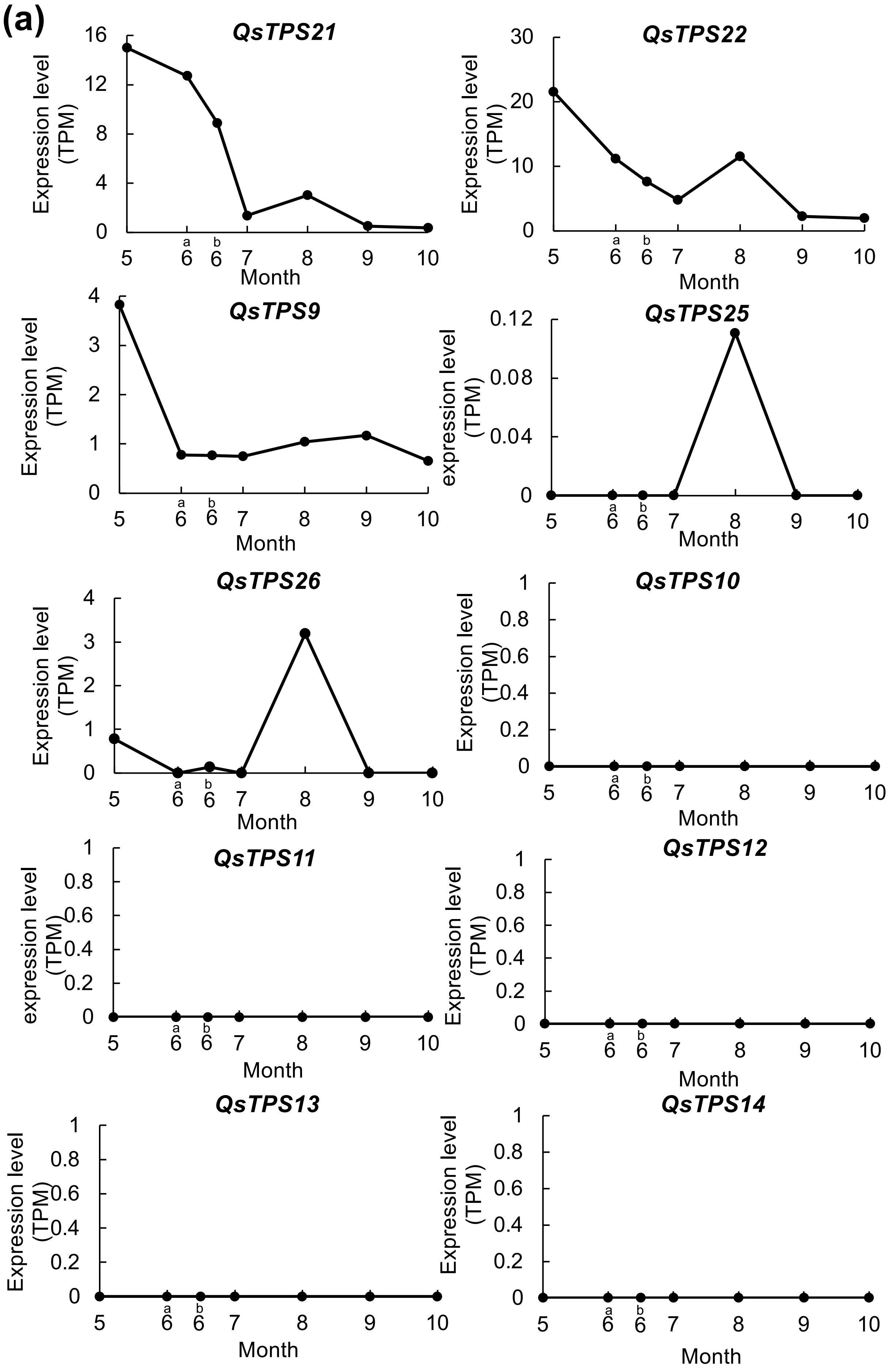


Supplementary Fig. S3. Seasonal expressions of each *QsTPS* in leaves and leaf buds. (-*continued*) The Seasonal expression data of each *QsTPS* in leaves (a) and buds (b) were individually shown (n = 1). N.D., not detected.

Koita *et al.*

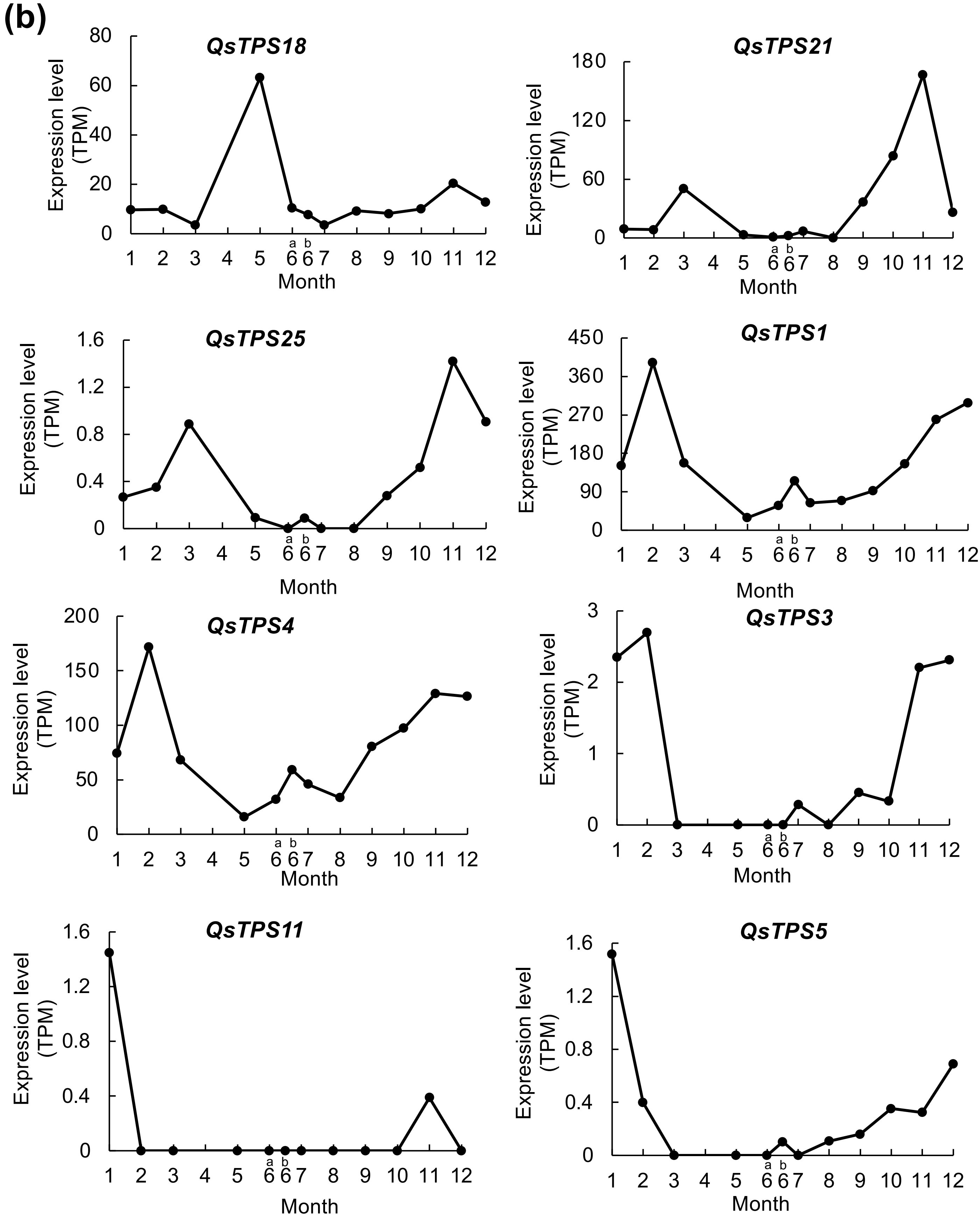


Supplementary Fig. S3. Seasonal expressions of each *QsTPS* in leaves and leaf buds. (*-continued*) The Seasonal expression data of each *QsTPS* in leaves (a) and buds (b) were individually shown (n = 1). N.D., not detected.

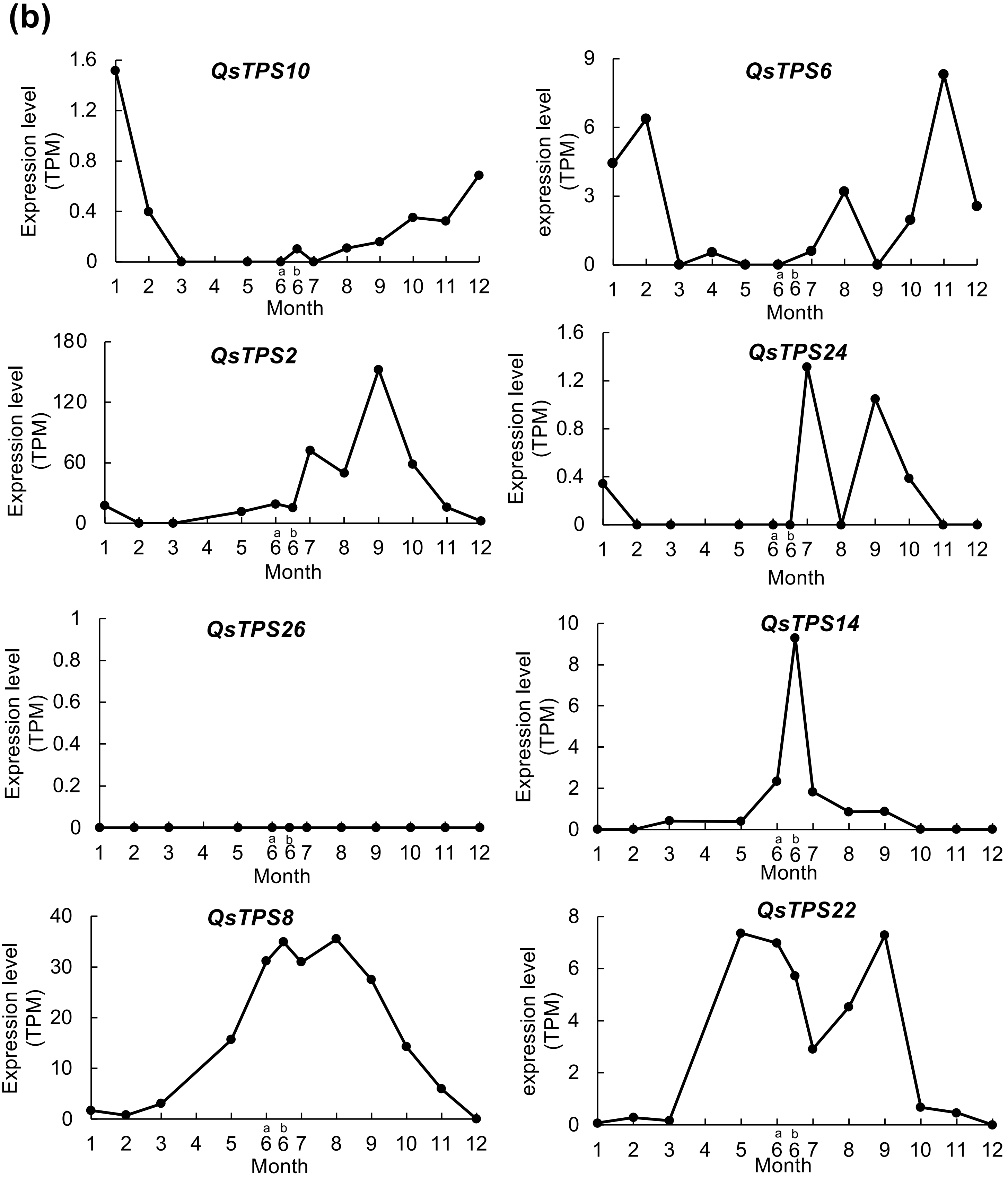


Supplementary Fig. S3. Seasonal expressions of each *QsTPS* in leaves and leaf buds. (-*continued*) The Seasonal expression data of each *QsTPS* in leaves (a) and buds (b) were individually shown (n = 1). N.D., not detected.

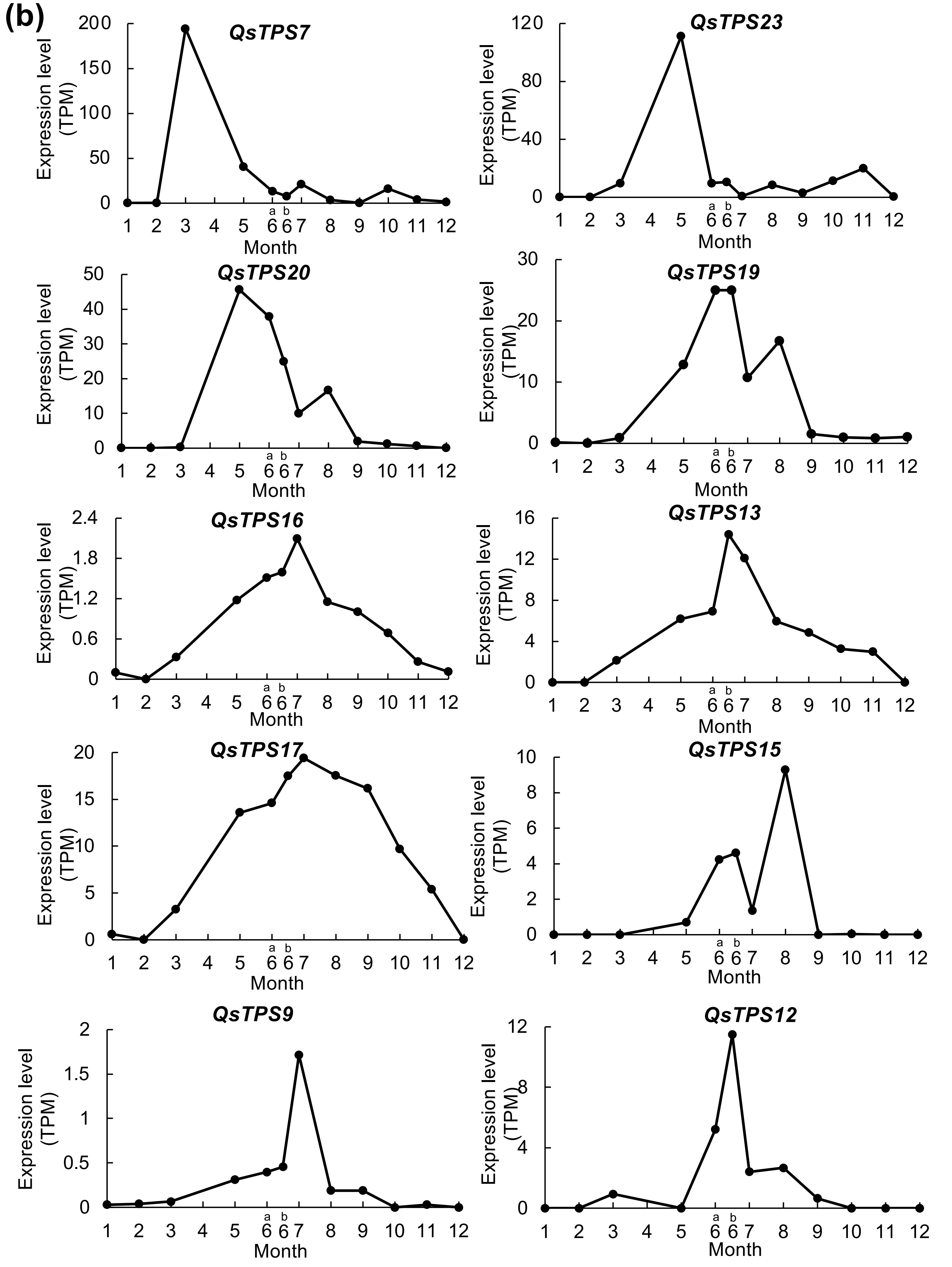


Supplementary Fig. S3. Seasonal expressions of each *QsTPS* in leaves and leaf buds. (*-continued*) The Seasonal expression data of each *QsTPS* in leaves (a) and buds (b) were individually shown (n = 1). N.D., not detected.

(a)

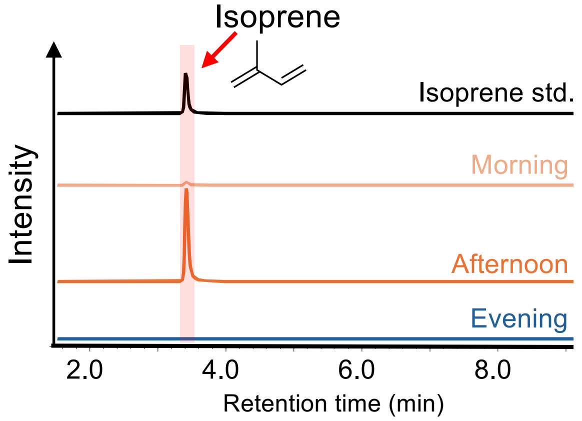


(b)

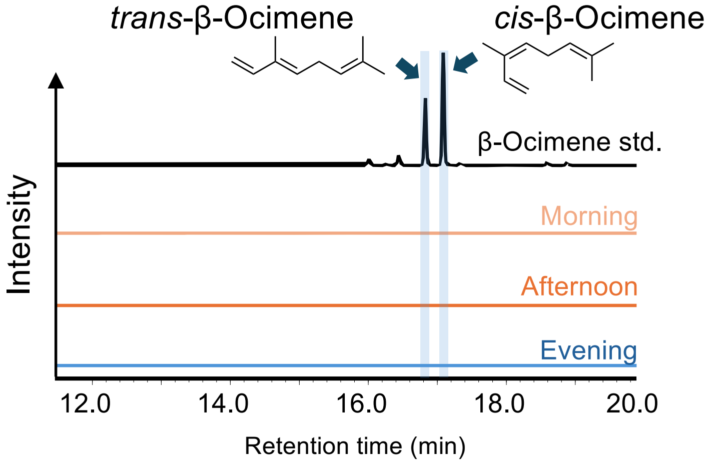


Supplementary Fig. S4. Diurnal isoprene emission and *QsIspS1* expression in *Q. serrata* leaves.
GC-MS chromatograms of isoprene ($m/z$ = 67) and monoterpenes ($m/z$ = 93) emitted from *Q. serrata* leaves are shown in (a) and (b), respectively. (c) and (d) show the amount of isoprene emission and *QsIspS1* expression in *Q. serrata l*eaves at morning (7:45-9:30 AM), noon (1:30-3:00 PM), and evening (6:30-8:30 PM). Bars indicate the mean of biological replicates, and dots represent the values of each replicate. An outlier was excluded from the morning sample by Dixon's Q test before statistical analysis, and the same individual was excluded from the VOC emission analysis. Games-Howell's test, $p^* < 0.05$, n = 5 (morning) and 6 (afternoon and evening). Different alphabets represent significance. Specific information on climate and diurnal conditions during the sampling days is provided in Supplementary Tables S6-9.

(c)

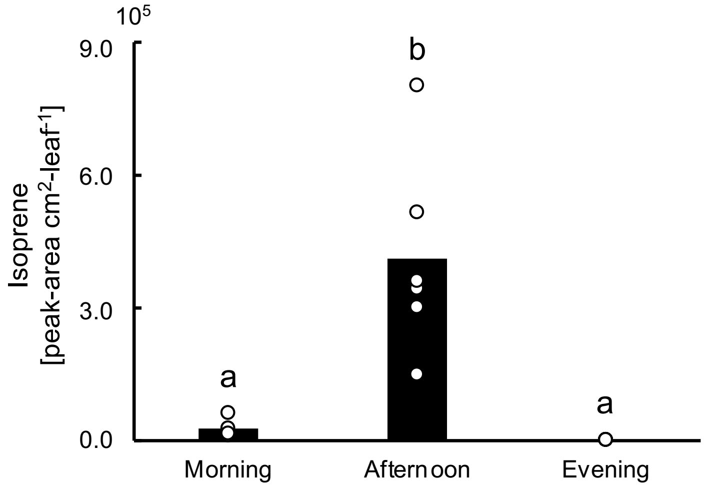


(d)

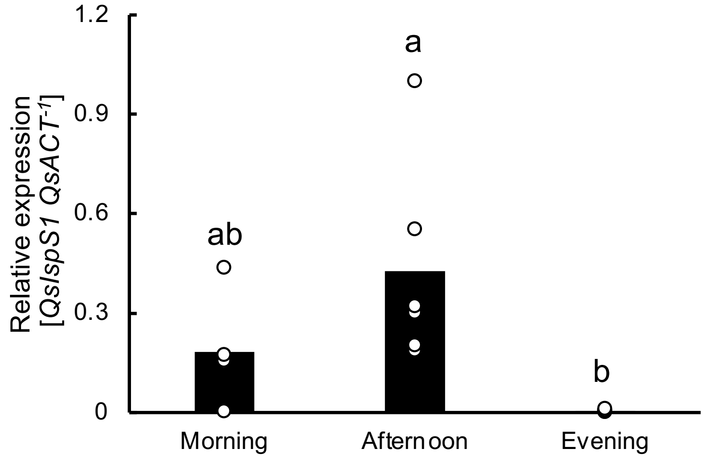


Supplementary Fig. S4. Diurnal isoprene emission and *QsIspS1* expression in *Q. serrata* leaves. (-*continued*)
GC-MS chromatograms of isoprene (*m/z* = 67) and monoterpenes (*m/z* = 93) emitted from *Q. serrata* leaves are shown in (a) and (b), respectively. (c) and (d) show the amount of isoprene emission and *QsIspS1* expression in *Q. serrata l*eaves at morning (7:45-9:30 AM), noon (1:30-3:00 PM), and evening (6:30-8:30 PM). Bars indicate the mean of biological replicates, and dots represent the values of each replicate. An outlier was excluded from the morning sample by Dixon's Q test before statistical analysis, and the same individual was excluded from the VOC emission analysis. Games-Howell's test, $p* < 0.05$, n = 5 (morning) and 6 (afternoon and evening). Different alphabets represent significance. Specific information on climate and diurnal conditions during the sampling days is provided in Supplementary Tables S6-9.

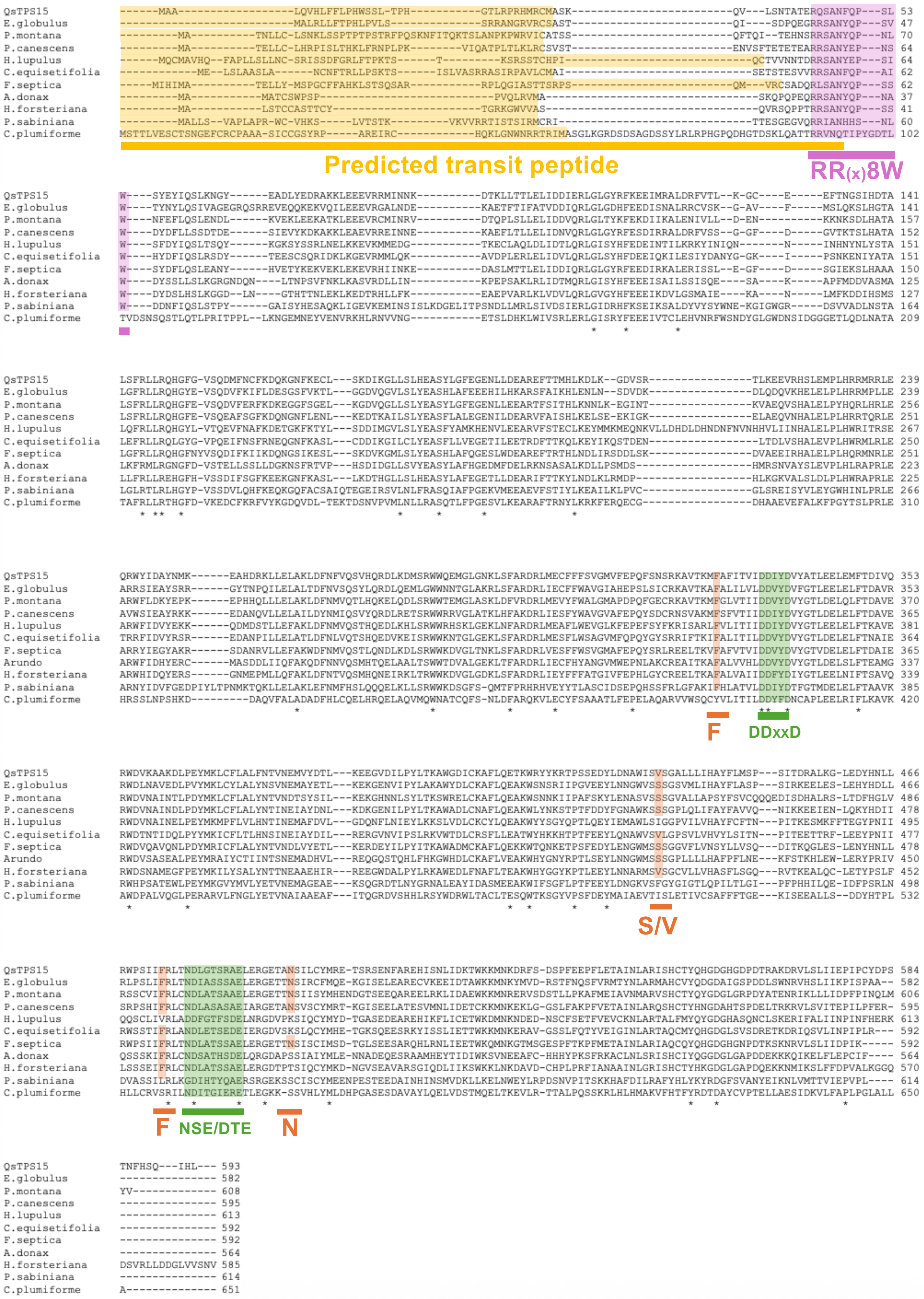


Supplementary Fig. S5. Sequence alignment of the gene product of *QsTPS15* alongside reported IspS polypeptide sequences.

The IspS sequences were aligned using Clustal Omega. The predicted transit peptide sequences, the D-rich $Mg^{2+}$-binding motifs (DDxxD and NSE/DTE), the RR(x)8W motif, and the four amino acids representing the "isoprene score" are highlighted in yellow, green, purple, and orange, respectively. The transit peptide sequences were predicted using the TargetP2.0 and DeepLoc2.1 programs.

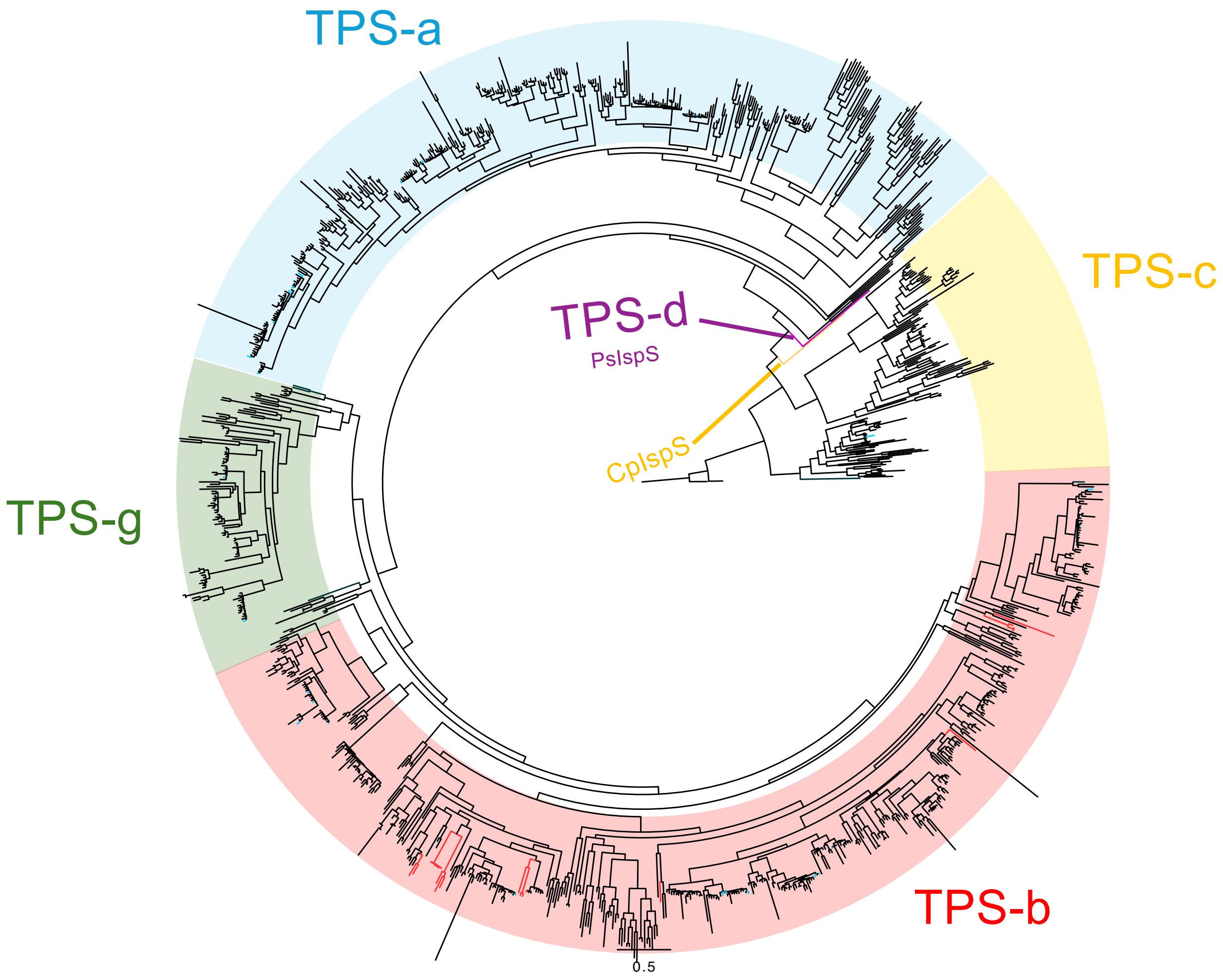


Supplementary Fig. S6. Phylogenetic tree of plant TPSs, including QsTPSs

The phylogenetic tree was constructed using QsTPSs, reported IspSs, and TPSs obtained from other plant taxa. The TPS-a, -b, -c, and -g subfamilies are colored blue, red, yellow, and green, respectively. The branches of the IspSs in subfamilies TPS-b, -c, and -d are highlighted in red, yellow, and purple, respectively. The branches of the 26 QsTPSs are colored light blue. The bar indicates 0.5 substitutions per site. CpIspS: *Calohypnum plumiforme* IspS, PsIspS: *Pinus sabiniana* IspS.

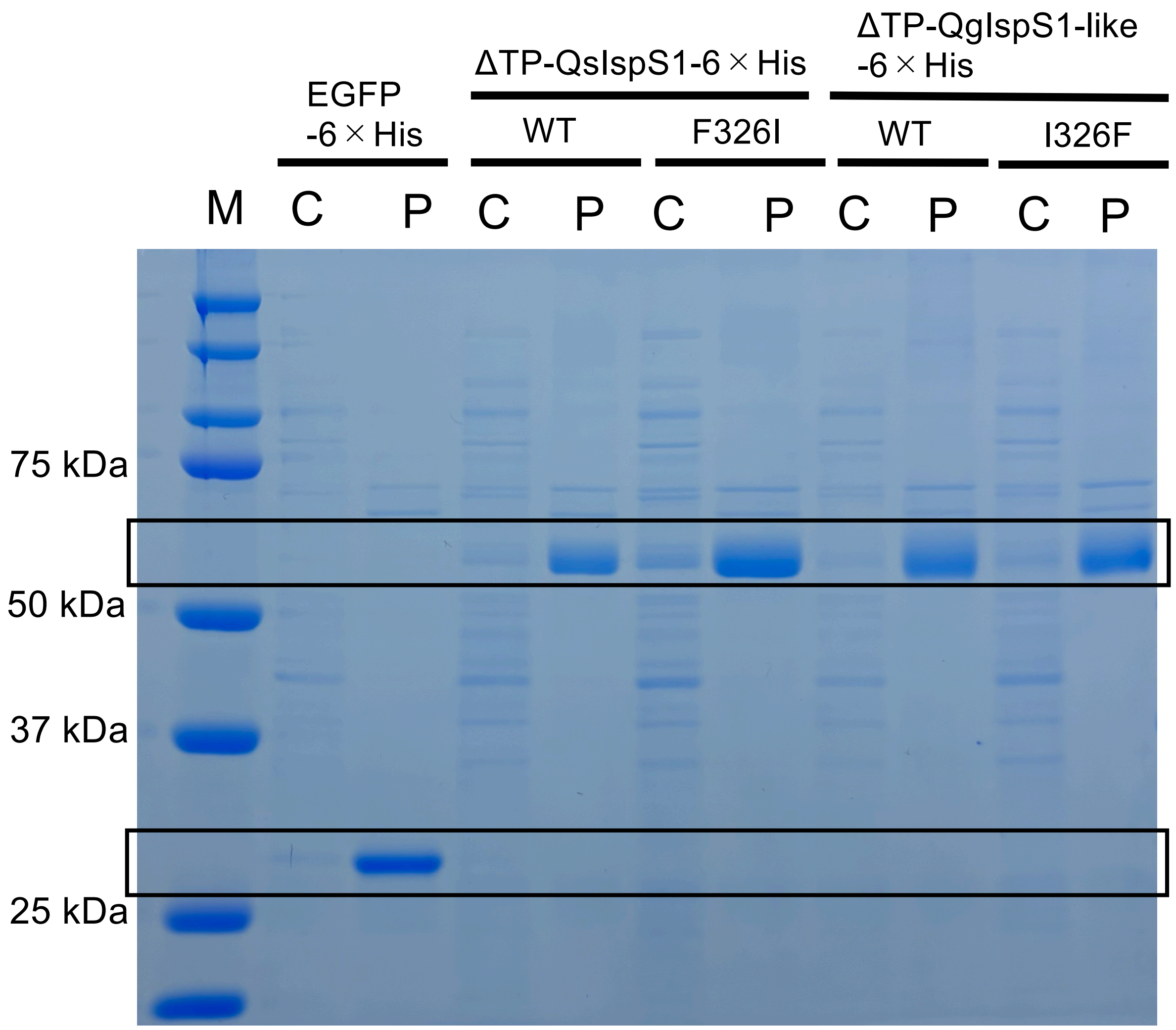


Supplementary Fig. S7. SDS-PAGE of recombinant proteins expressed in *E. coli* that are affinity-purified with Ni-agarose column. The following proteins are shown: EGFP, ΔTP-QsIspS1-6 × His, ΔTP-QgIspS1-like-6 × His, ΔTP-QsIspS1(F326I) -6 × His and ΔTP-QgIspS1-like(I326F) -6 × His.

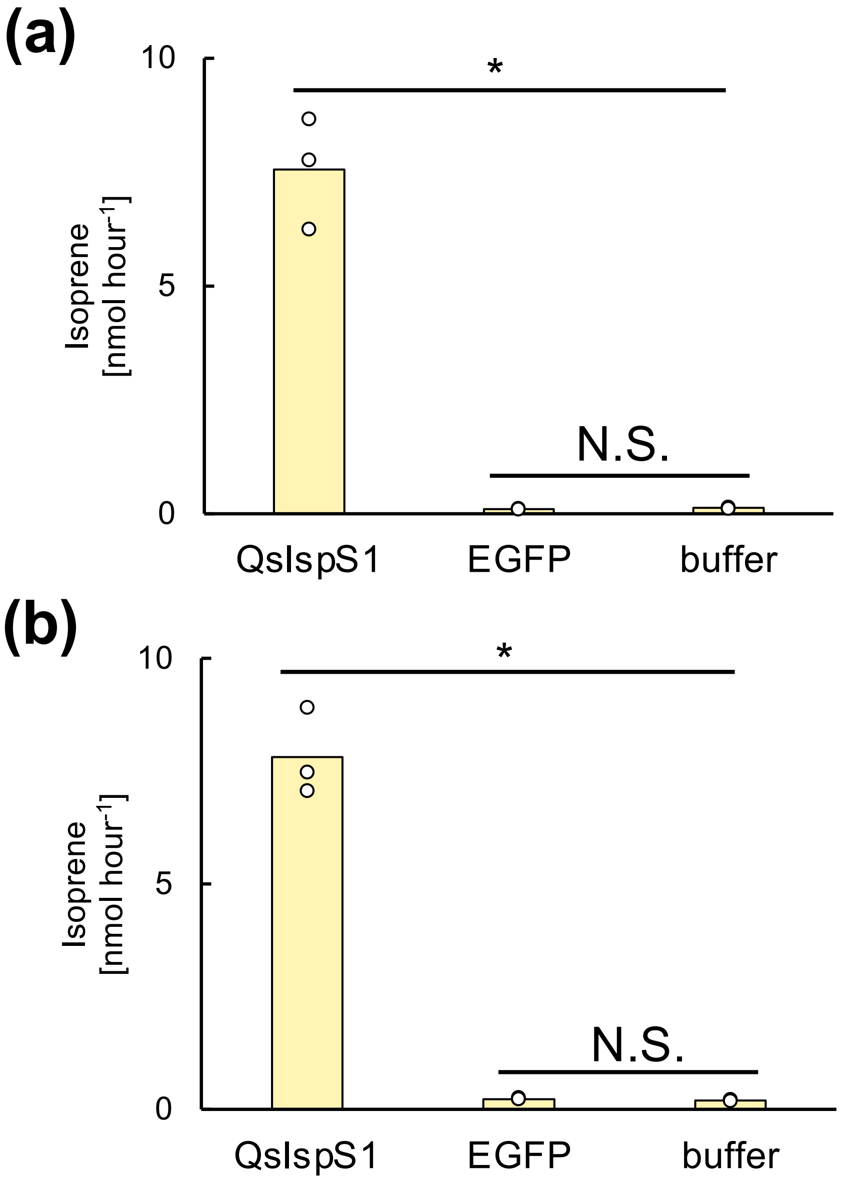


Supplementary Fig. S8. Negative control assays of QsIspS1. The following proteins are shown:ΔTP-QsIspS1-6 × His, EGFP-6 × His, transiently expressed in *E. coli* (a) or QsIspS1, EGFP, transiently expressed in *N. benthamiana* (b). The reaction buffer was also assayed with DMAPP. Bars indicate the mean of biological replicates, and dots represent the values of each replicate. Dunnett's test ($p^* < 0.05$, n = 3). The asterisk indicates a statistically significant difference. N.S., not significant.

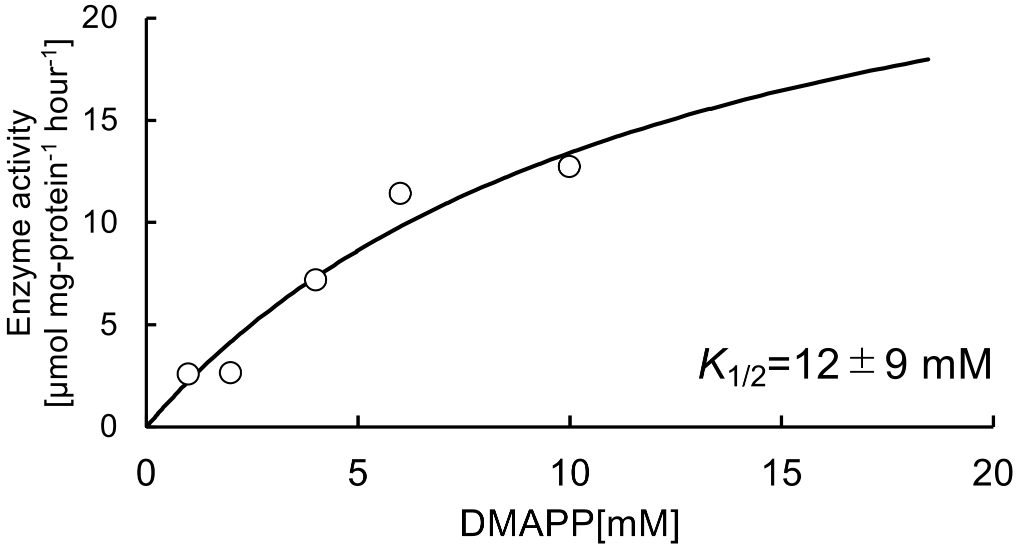


Supplementary Fig. S9. Kinetic analysis of recombinant QsIspS1 with DMAPP as its substrate.

Kinetic analysis of QsIspS1 for DMAPP using ΔTP-QsIspS1-6×His as the enzyme. Reaction mixtures were incubated for 30 min with different concentrations of DMAPP (0–20 mM). The $K_{1/2}$ of QsIspS1 was estimated to be 12 ± 9 mM, which was calculated using nonlinear fitting in SigmaPlot 14.5.

```
QsIspS1         MAALQVHLFFLPHWSSLTPHGTLRPRHMRCMASKQVLSNTATERQSANFQPSLWSYEYIQ 60
QgIspS1-like    MAALQVHLFFLPHWSSLTPHGTLRPRHMRCMASKQVLSNTATKRQSANYLPSLWSYEYIQ 60
LeIspS1-like    MAALQVHLFSLPHWSSFTPHGTLRPRHMRCMASKQVLSNTATKRQSANYLPSLWSYEYIQ 60
                ********* ******.*************************:*****. **********
                   Predicted transit peptide                  RR(x)8W

QsIspS1         SLKNGYEADLYEDRAKKLEEEVRRMINNKDTKLLTTLELIDDIERLGLGYRFKEEIMRAL 120
QgIspS1-like    SLKNGYEADLYEDRAKKVEEEVRRMINNKDAKLLTTLELIDDIERLGLGYRFKEEIMRAL 120
LeIspS1-like    SLKNGYEADLYKDRAKKLEEEVRRMINNKDTKLLTTLELIDDIERLGLGYRFKEEIIRAL 120
                ***********:*****:************:*************************:***

QsIspS1         DRFVTLKGCEEFTNGSIHDTALSFRLLRQHGFGVSQDMFNCFKDQKGNFKECLSKDIKGL 180
QgIspS1-like    DRFVTLKGCEEFTNASIHDTALSFRLLRQHGFGVSQDVFNCFKDQKGNFKECLSKDIKGL 180
LeIspS1-like    DRFVTLKGCKEFTNGSIHDTALSFRLLRQHGFGVSQDIFNCFKDQKENF*---------- 169
                *********:****.**********************:******** **

QsIspS1         LSLHEASYLGFEGENLLDEAREFTTMHLKDLKGDVSRTLKEEVRHSLEMPLHRRMRRLEQ 240
QgIspS1-like    LSLHEASYLGFEGENLLDEAMEFTTMHLKDLKGDVSRTLKEEVRHSLEMPLHRRMRRLEQ 240
LeIspS1-like    ------------------------------------------------------------ 169

QsIspS1         RWYIDAYNMKEAHDRKLLELAKLDFNFVQSVHQRDLKDMSRWWQEMGLGNKLSFARDRLM 300
QgIspS1-like    RWYIDAYNMKEAHNRKLLELAKLDFNIVQSVHQRDLKDMSRWWKEMGLGNKLSFARDRLM 300
LeIspS1-like    ------------------------------------------------------------ 169

QsIspS1         ECFFFSVGMVFEPQFSNSRKAVTKMFAFITVIDDIYDVYATLEELEMFTDIVQRWDVKAA 360
QgIspS1-like    ECFFWAVGMVFEPQFSNCRKELTKVISLITIIDDVYDVYATLEELEMFTDIVQRWDVKAV 360
LeIspS1-like    ------------------------------------------------------------ 169
                                         F        DDxxD

QsIspS1         KDLPEYMKLCFLALFNTVNEMVYDTLKEEGVDILPYLTKAWGDICKAFLQETKWRYYKRT 420
QgIspS1-like    KDLPEYMKLCFLALYNTVNEMVYDTLKEQGVDILPYLTKAWGDLCKAFLQETKWRYYKHT 420
LeIspS1-like    ------------------------------------------------------------ 169

QsIspS1         PSSEDYLDNAWISVSGALLLIHAYFLMSPSITDRALKGLEDYHNLLRWPSIIFRLTNDLG 480
QgIspS1-like    PSSEDYLDNAWMSSSGALLLVHAYFLMSPSITNRALEGLEDYHNLLRWPSIIFRLCNDLV 480
LeIspS1-like    ------------------------------------------------------------ 169
                             S/V                                  F  NSE/DTE

QsIspS1         TSRAELERGETANSILCYMRETSRSENFAREHISNLIDKTWKKMNKDRFSDSPFEEPFLE 540
QgIspS1-like    TSMTELERGETANSILCYMRETSLSEDFAREHISNLIDKTWKKMNKDRFSDSPFEEPFLE 540
LeIspS1-like    ------------------------------------------------------------ 169
                             N

QsIspS1         TAINLARISHCTYQHGDGHGDPDTRAKDRVLSLIIEPIPCYDPSTNFHSQIHL* 593
QgIspS1-like    TAINLARISHCTYQHGDGHSAPDSRSKDRVLSLIIEPIPCYDPSTNFHGHIHL* 593
LeIspS1-like    ------------------------------------------------------ 169
```

Supplementary Fig. S10. Sequence alignment of the gene product of *QsIspS1, QgIspS1-like,* and *LeIspS1-like*.

Sequences were aligned using Clustal Omega. Predicted transit peptide sequences, D-rich $Mg^{2+}$-binding motifs (DDxxD and NSE/DTE), RR(x)8W motif, and the four amino acids representing the “isoprene score” are highlighted in yellow, green, purple, and orange, respectively. The transit peptide sequences were predicted using the TargetP2.0 program.

```
QsIspS1        ATGGCAGCACTCCAAGTCCATCTCTTCTTCTTGCCCCATTGGAGCTCCTTAACACCTCAT     60
LeIspS1-like   ATGGCAGCACTCCAAGTCCATCTCTTCTCCTTGCCCCATTGGAGCTCCTTCACACCTCAT     60
               **************************** ********************* *********

QsIspS1        GGAACACTGAGGCCTAGGCATATGCGGTGCATGGCGAGCAAACAAGTGCTTTCTAATACA     120
LeIspS1-like   GGAACACTGAGGCCTAGGCACATGCGGTGCATGGCGAGCAAACAAGTGCTTTCCAATACA     120
               ******************** ******************************** ******

QsIspS1        GCAACTGAAAGGCAGTCGGCCAATTTCCAGCCGAGCCTCTGGAGTTACGAATATATACAG     180
LeIspS1-like   GCAACTAAAAGACAGTCGGCCAATTACCTGCCGAGCCTCTGGAGTTATGAATATATACAG     180
               ****** **** ************* ** ****************** ************

QsIspS1        TCATTGAAGAATGGTTATGAGGCTGACCTGTATGAAGATAGGGCAAAGAAGCTGGAGGAA     240
LeIspS1-like   TCATTGAAGAATGGTTATGAGGCTGATCTGTATAAAGATAGGGCAAAGAAGCTGGAGGAA     240
               ************************** ****** **************************

QsIspS1        GAAGTGAGGAGAATGATCAATAACAAAGATACGAAGTTATTGACCACACTTGAATTAATT     300
LeIspS1-like   GAAGTGAGGAGAATGATCAATAACAAAGATACGAAGTTATTGACTACACTTGAATTAATT     300
               ******************************************** ***************

QsIspS1        GATGACATTGAACGGCTCGGTTTGGGATACCGGTTCAAGGAAGAAATAATGAGAGCCCTT     360
LeIspS1-like   GATGACATTGAACGGCTCGGTTTGGGATACCGGTTCAAGGAAGAAATAATTAGAGCCCTT     360
               ************************************************** *********

QsIspS1        GACAGATTTGTCACTTTGAAAGGATGTGAAGAGTTTACAAATGGTAGTATCCATGACACT     420
LeIspS1-like   GACAGATTTGTCACTTTGAAAGGATGTAAAGAGTTTACAAATGGTAGTATCCATGACACT     420
               ************************** *********************************

QsIspS1        GCCCTCAGCTTCAGGCTCCTTAGACAACATGGGTTTGGCGTCTCTCAAGATATGTTTAAT     480
LeIspS1-like   GCCCTCAGCTTCAGGCTCCTTAGACAACATGGGTTTGGTGTCTCTCAAGATATATTTAAT     480
               ************************************** ************** ******

QsIspS1        TGTTTCAAGGACCAAAAGGGAAATTTCAAGGAGTGCCTTAGCAAGGACATCAAAGGATTG     540
LeIspS1-like   TGTTTCAAGGACCAAAAGGAAAATTTCTAGGAGTGCCTTAGCAAGGACAATAAAGGATTG     540
               ******************* ******** ********************  *********
                                           stop
                                           codon
```

Supplementary Fig. S11. Sequence alignment of the gene of *QsIspS1* and *LeIspS1-like*. Sequences were aligned using Clustal Omega. A nucleotide that induces a stop codon is highlighted in blue.

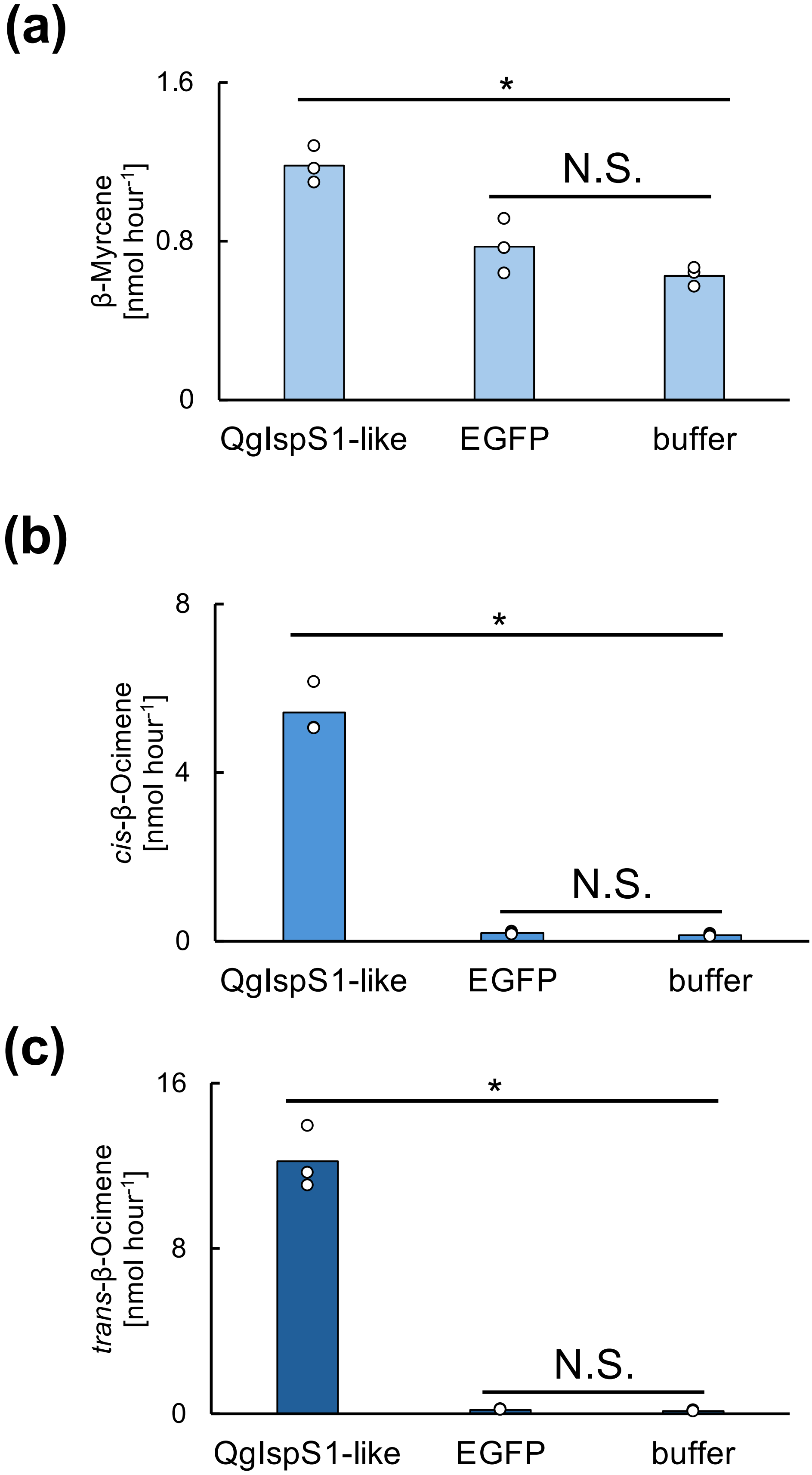


Supplementary Fig. S12. Negative control assays of QgIspS1-like. ΔTP-QgIspS1-like-6 × His, EGFP-6 × His, transiently expressed in *E. coli*, and the reaction buffer were assayed with GPP. Bar graphs show the amount of β-myrcene (a), *cis*-β-ocimene (b), and *trans*-β-ocimene (c). Bars indicate the mean of biological replicates, and dots represent the values of each replicate. Dunnett's test ($p^* < 0.05$, n = 3). The asterisk indicates a statistically significant difference. N.S., not significant.

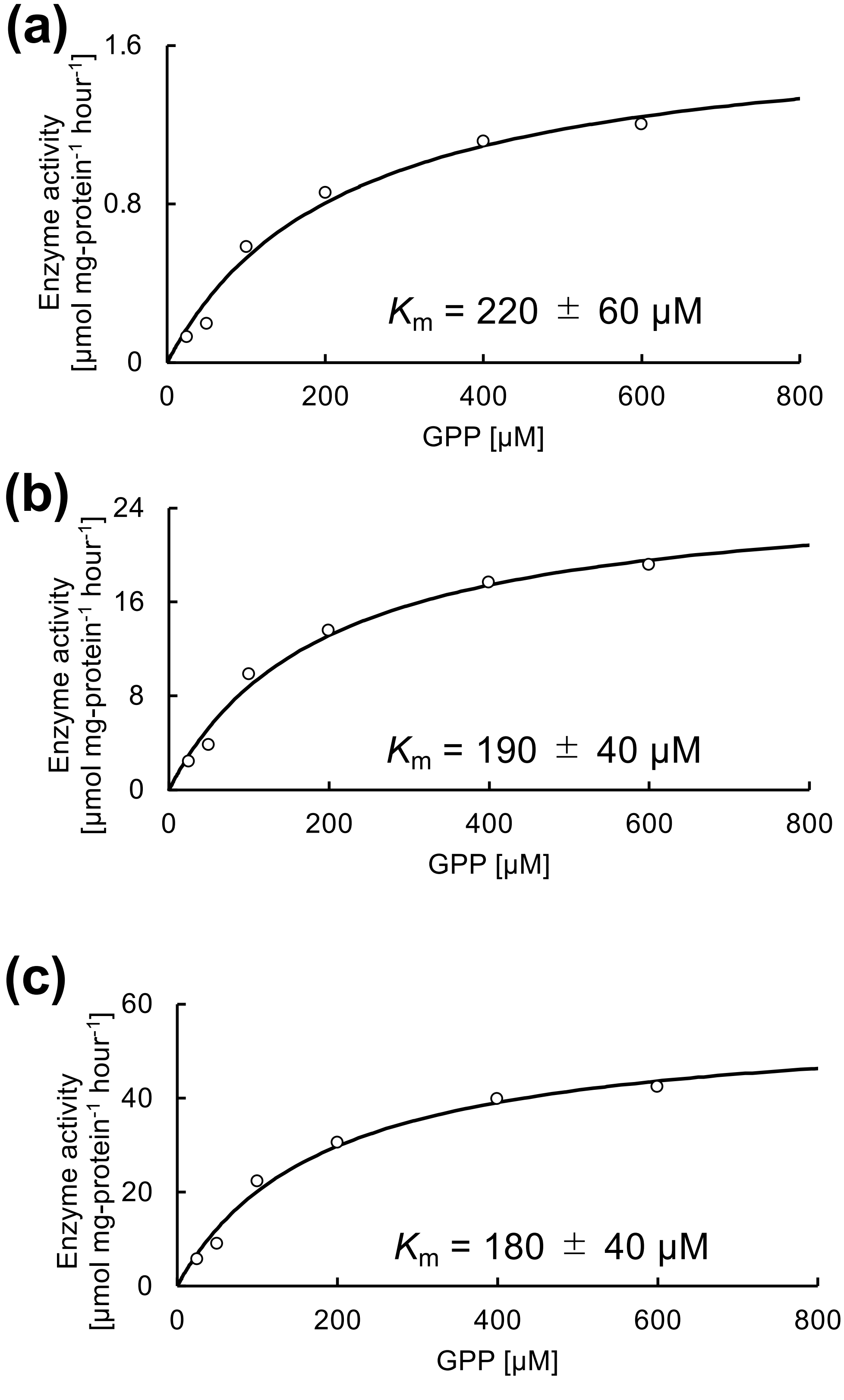


Supplementary Fig. S13 Kinetic analysis of recombinant QgIspS1-like using GPP as its substrate.

Microsomes were incubated for 30 min with various concentrations of GPP (0–600 μM) and recombinant ΔTP-QgIspS1-like-6×His was used as the enzyme. The $K_m$ values for QgIspS1-like were estimated to be 220 ± 60 μM for β-myrcene (a), 190 ± 40 μM for *cis*-β-ocimene (b), and 180 ± 40 μM for *trans*-β-ocimene (c). These $K_m$ values were calculated using nonlinear fitting in SigmaPlot 14.5.

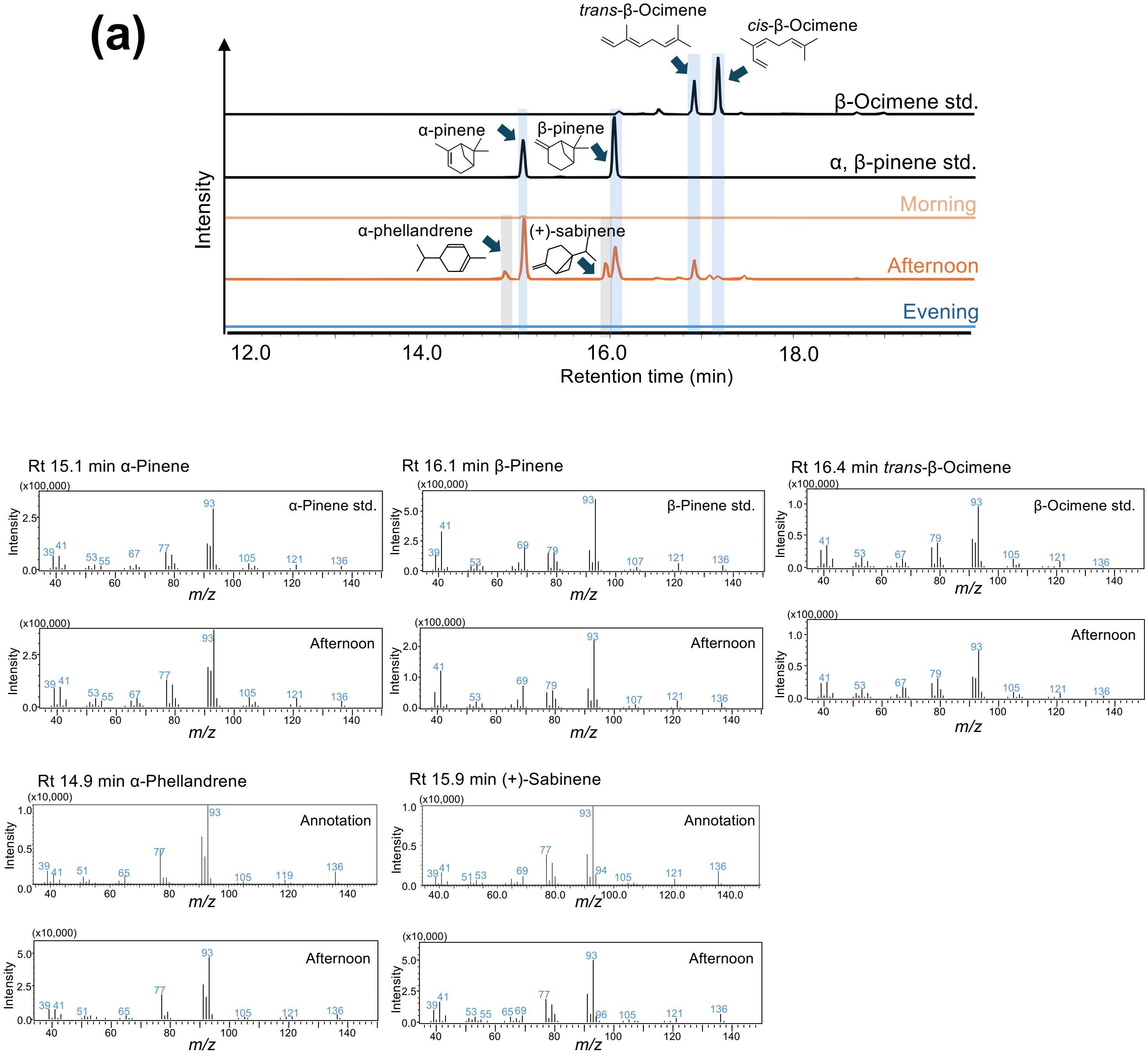


Supplementary Fig. S14. Diurnal monoterpene emission and *QgIspS1-like* expression in *Q. glauca* leaves.

Monoterpene emissions are shown in the GC-MS chromatogram (*m/z* = 93) and the MS spectrum of the major five peaks (a). Isoprene emissions are shown in the chromatogram (*m/z* = 67) (b). The amounts of total ocimene emissions (c) and *QgIspS1-like* expression (d) in *Q. glauca* leaves were measured at morning (7:45–9:30 AM), afternoon (1:30–3:00 PM), and evening (6:30–8:30 PM) were measured. Bars indicate the mean of biological replicates, and dots represent the values of each replicate. Games-Howell's test, $p^* < 0.05$, n = 6 (morning, afternoon, and evening). The different alphabets represent significance. Specific information on climate and diurnal conditions during the sampling days is provided in Supplementary Tables S6-9.

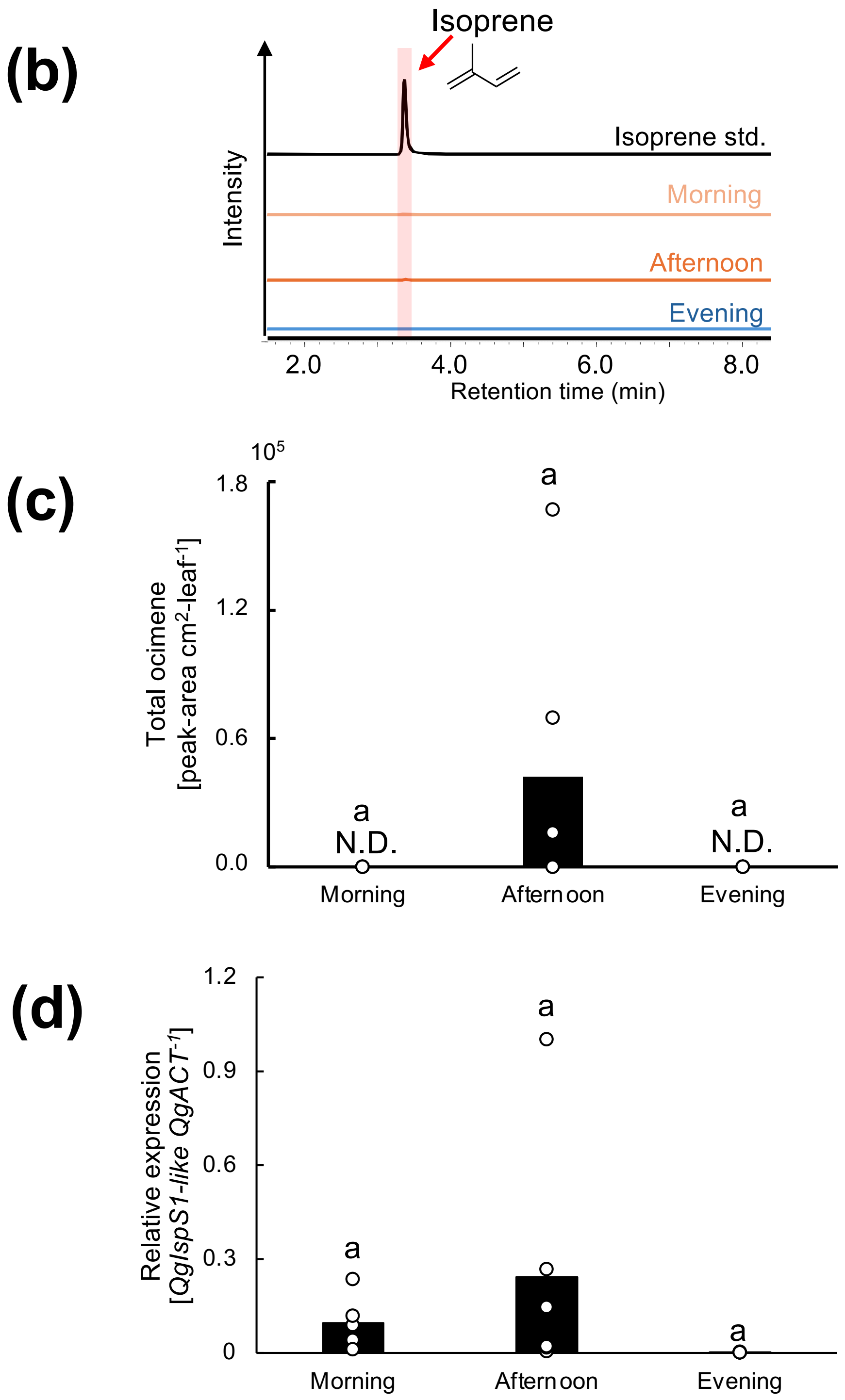


Supplementary Fig. S14. Diurnal monoterpene emission and *QgIspS1-like* expression in *Q. glauca* leaves. (*-continued*)
Monoterpenes emissions are shown in the GC-MS chromatogram ($m/z$ = 93) and MS spectrum of the major 5 peaks (a), and isoprene emission in the chromatogram ($m/z$ = 67) (b). The amounts of total ocimene emission (c) and *QgIspS1-like* expression (d) in *Q. glauca* leaves at morning (7:45-9:30 AM), afternoon (1:30-3:00 PM), and evening (6:30-8:30 PM) were measured. Bars indicate the mean of biological replicates, and dots represent the values of each replicate. Games-Howell's test, $p* < 0.05$, n = 6 (morning, afternoon, and evening). The different alphabets represent significance. Specific information on climate and diurnal conditions during the sampling days is described in Supplementary Tables S6-9.

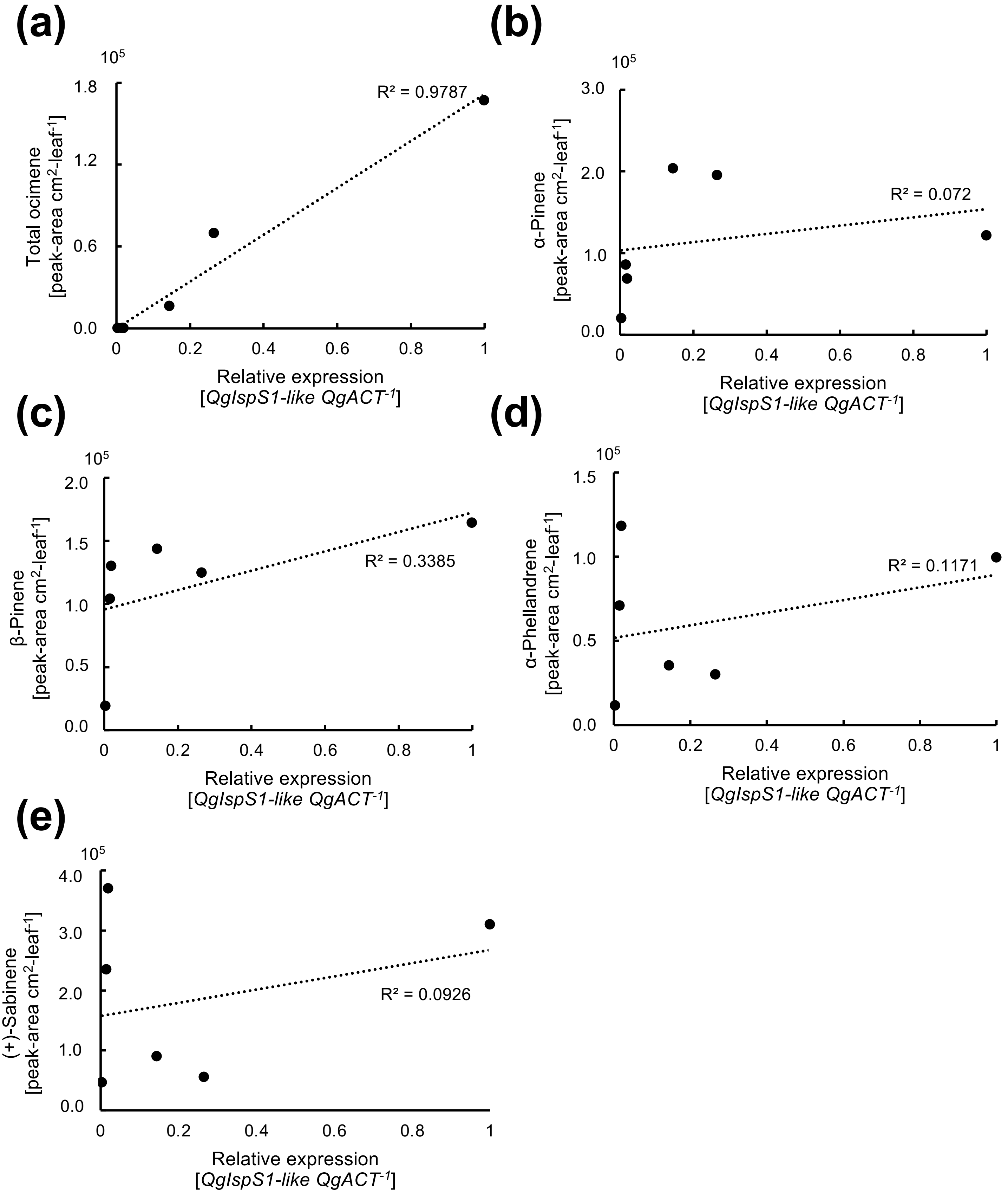


Supplementary Fig. S15. Correlation of monoterpenes emission and expression levels in the afternoon. Correlation of *QgIspS1-like* expression with the amounts of total ocimene (a), α-pinene (b), β-pinene (c), α-phellandrene (d), and (+)-sabinene (e) emissions.

Koita *et al.*

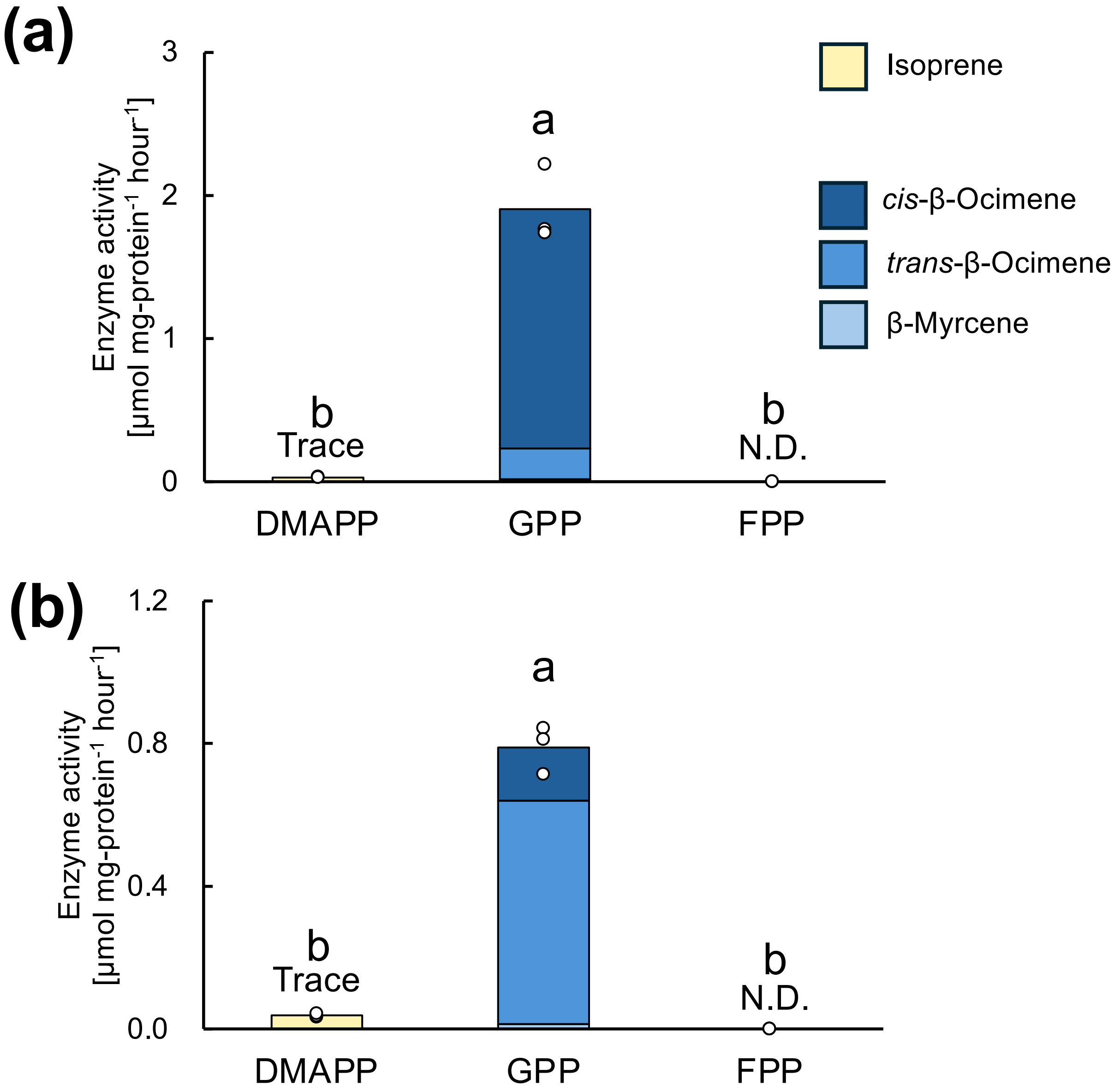


Supplementary Fig. S16. Enzymatic characterization of mutant QsIspS1-like (F326I) and QgIspS1-like (I326F).
Recombinant proteins of ΔTP-QsIspS1-like(F326I)-6 × His (a) and ΔTP-QgIspS1-like(I326F)-6 × His, which were expressed in *E. coli*, were used after affinity purification. The substrate preferences of QsIspS1-like(F326I) (a) and QgIspS1-like(I326F) (b) are shown with DMAPP, GPP, and FPP as substrates. N.D., not detected. Bars indicate the mean of biological replicates, and dots represent the values of each replicate. Statistical analysis was performed using the Tukey-Kramer test. Different letters indicate statistically significant differences (n = 3).

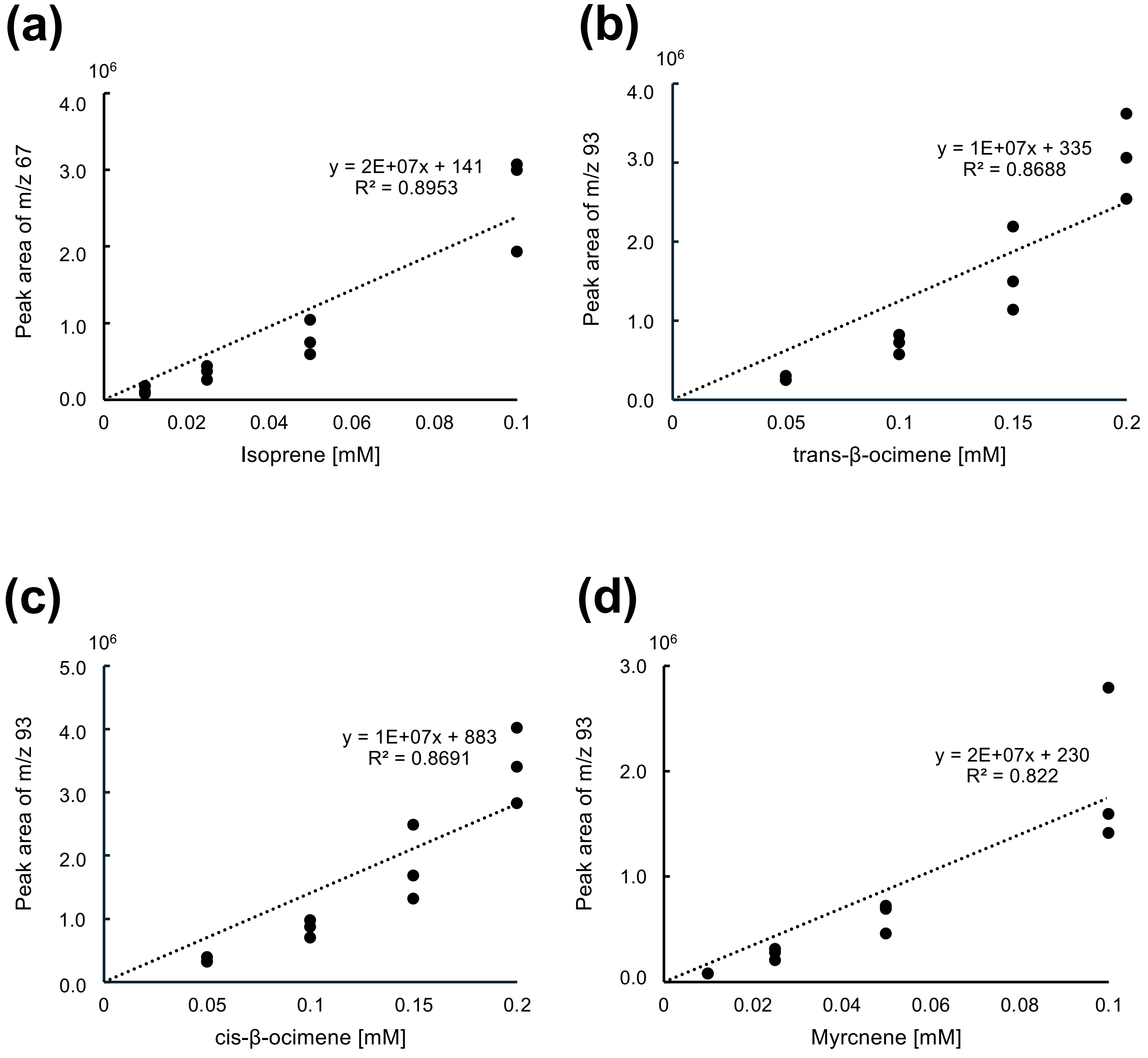


Supplementary Fig. S17. Standard curves of isoprene and monoterpenes analyzed in this study. Standard curves of isoprene (a), *trans*-β-ocimene (b), *cis*-β-ocimene (c), and myrcene (d).

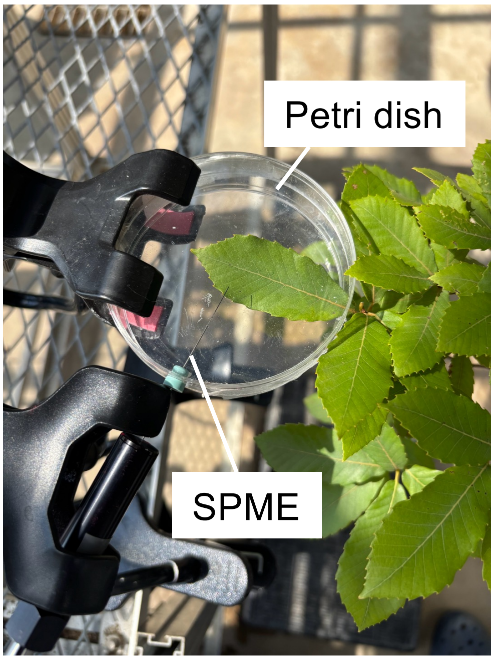


Supplementary Fig. S18. VOC trapping from leaves of *Quercus* trees. A leaf of either *Q. serrata* (the photo image) or *Q. glauca* was sandwiched with a Petri dish (90 mmφ x19 mm) through a hole that was sized for the leaf petiole. The SPME fiber was inserted from the other side of the hole. The VOCs in the headspace were trapped for 15 min.
